# A coral-linked adaptive radiation in butterflyfishes

**DOI:** 10.64898/2026.11.09.765357

**Authors:** Konstantinos Andriotis, Jennifer R. Hodge, Chase D. Brownstein, Michael E. Alfaro, Peter C. Wainwright, Thomas J. Near

## Abstract

Coral reefs harbour a disproportionate share of marine biodiversity, yet whether reef fish diversity reflects steady accumulation or rapid, ecologically driven diversification remains unclear. Butterflyfishes and bannerfishes (*Chaetodontidae*) include more than half of all coral-feeding fish species, making them a natural test of adaptive radiation on coral reefs. Here we present a time-calibrated phylogeny of 120 of the 138 species, built from 1,009 ultraconserved elements. The species-rich butterflyfishes (*Chaetodontinae*) are a recent radiation that originated in the Late Miocene (7.78 million years ago; 95% highest posterior density 7.13–8.49 million years ago), with much of their extant species diversity arising within the past 2 million years. The crown diversification rate of butterflyfishes is the highest among 29 comparator reef-fish lineages. Ancestral state reconstructions indicate that the ancestor of *Chaetodontinae* was a facultative corallivore, that a unique jaw joint arose before obligate corallivory, and that obligate corallivory then evolved repeatedly from facultative ancestors, alongside derived tooth morphologies. Together, these results suggest that Late Miocene coral reef turnover and the diversification of branching corals generated an ecological opportunity that functional innovations allowed butterflyfishes to exploit, extending the adaptive radiation framework to open marine seascapes beyond the island and lake systems where it has most often been applied.

---

Adaptive radiation can produce rapid ecological diversification when lineages gain access to newly available niche space, but most well-supported examples come from geographically bounded settings, such as islands^1,2^ and lakes^3,4^, where the physical boundaries of the system, and therefore the colonizing lineage’s history within it, are more readily delimited^5,6^. Coral reefs support some of the most species-rich vertebrate assemblages on Earth^7–10^, yet whether similarly rapid, ecologically driven diversification occurs on coral reefs remains unclear^11–13^. If ecological opportunity has shaped reef fish radiations, then major episodes of reef reorganization should be mirrored by accelerated diversification in coral-dependent consumers, coupled to predictable ecological shifts and associated functional innovations^14–16^.

Reef fish diversity has long been viewed as the product of steady lineage accumulation over tens of millions of years, an inference drawn from time-calibrated phylogenies that reflected a protracted tempo of diversification^9,17,18^. Recent fossil, genomic, and geological evidence instead points to a rapid period of coral reef growth and compositional turnover in the late Oligocene and Early Miocene^19–21^. These observations implicate reef habitat expansion and the proliferation of coral-dependent niche space as the major sources of ecological opportunity that produced present-day reef biodiversity^16,19–21^. Whether such opportunity translated into adaptive radiations of lineages reliant on living corals remains unclear.

*Chaetodontidae* is a globally-distributed lineage closely associated with reef corals^22^. The clade comprises 138 species in two lineages, the bannerfishes (*Heniochinae*; 28 species) and the butterflyfishes (*Chaetodontinae*; 110 species), and exhibits conspicuous diversity in feeding morphology, depth distribution, dietary specialization and coloration^11,23–27^.

Although *Chaetodontidae* represent a small fraction of overall reef-associated fish diversity, they account for more than half (54%) of all fish species reported to eat coral and 61% of obligate corallivores^26,28^. This outsized ecological role is matched by a suite of functional innovations that include the intramandibular joint (IMJ) between the dentary and articular bones, which permits independent flexion of the lower jaw and is thought to increase feeding efficiency on benthic prey^29,30^, coral-associated dental specialization^31–33^, gut elongation associated with coral-based diets^34^, and the capacity to bite with fully protruded jaws^29,35,36^ (Fig. 1).

**Fig. 1.**
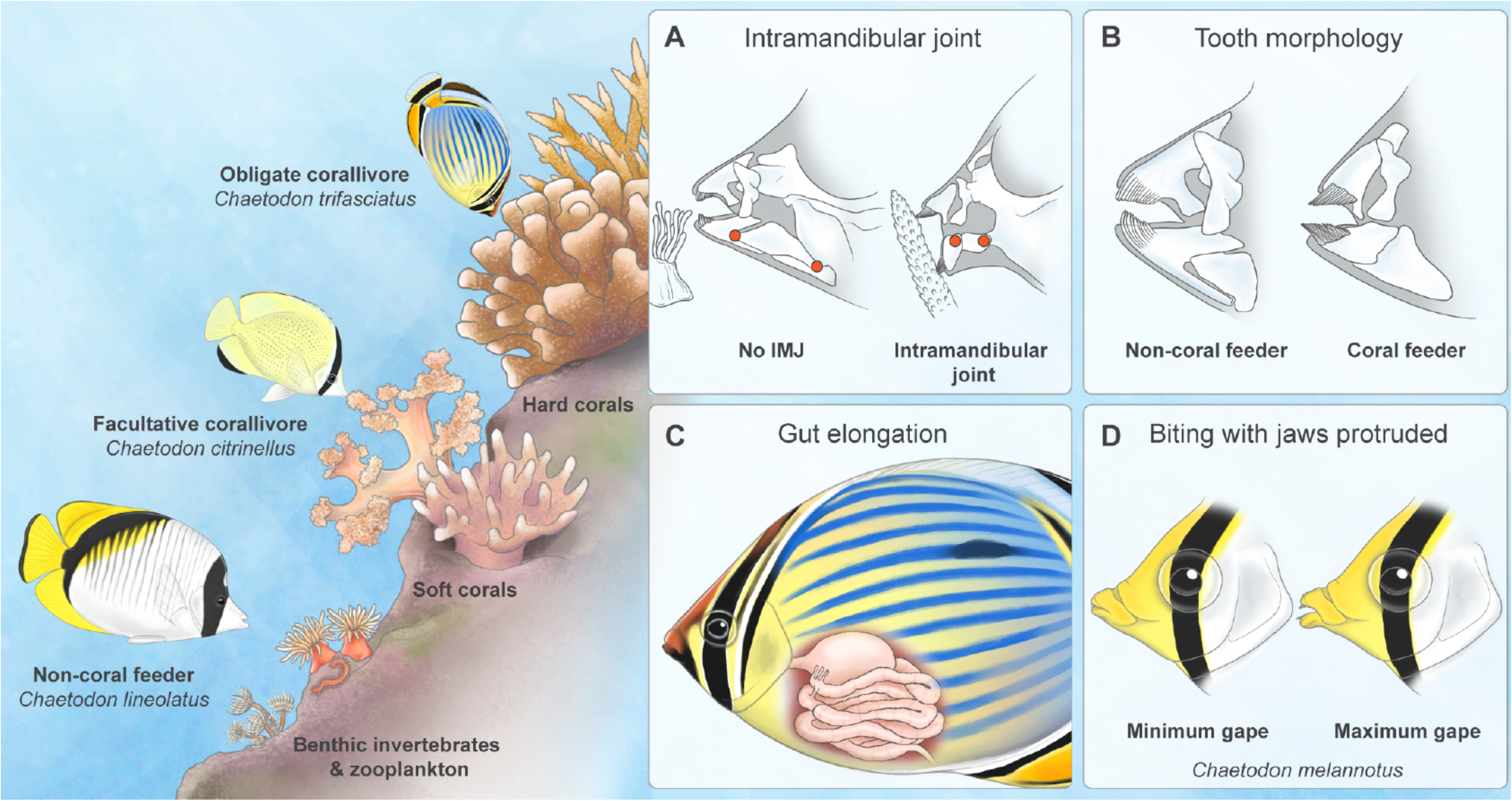
Functional innovations and feeding strategies of butterflyfishes. Left, feeding strategies on a reef: an obligate corallivore (*Chaetodon trifasciatus*), a facultative corallivore (*C. citrinellus*), and a non-coral feeder (*C. lineolatus*), with hard corals, soft corals, and benthic invertebrates and zooplankton as food resources. Right, four functional innovations associated with coral feeding: (A) the intramandibular joint (IMJ), absent (left) or present (right) in the lower jaw (red dots mark the lower-jaw joints); (B) the two extremes of tooth morphology in butterflyfishes: the basic condition (left), with a single row of short, slender teeth on each jaw, and the most complex condition (right), with dense pads of long, thin teeth at the front of the jaws; (C) gut elongation associated with the amount of coral consumption; and (D) biting with the fully protruded jaws, shown at minimum and maximum gape in the obligate corallivore *C. melannotus*. Illustrations by Julia Johnson (https://www.lifesciencestudios.com/).

Within *Chaetodontidae*, coral feeding is concentrated in butterflyfishes (*Chaetodontinae*), which contain every obligate corallivore and all but five species of facultative corallivores in our dataset, and the IMJ and coral-associated dental specializations occur only in *Chaetodon*. These specializations make *Chaetodontinae* a compelling system to test whether adaptive radiation can proceed on coral reefs without the strong geographic isolation that has characterized many classic examples^1–4,37,38^. Here, using a densely sampled, time-calibrated phylogeny of *Chaetodontidae*, we test whether *Chaetodontinae* is exceptionally species-rich for its age relative to the family as a whole and to other lineages of reef fishes, and whether ecological transitions and functional innovations align with this diversification within *Chaetodontinae*. Our results suggest that *Chaetodontinae* radiated more rapidly than nearly all other reef-fish lineages in the Late Miocene, coincident with turnover of reef-building corals^21^, and that obligate corallivory evolved repeatedly from facultative ancestors after the origin of the intramandibular joint, consistent with an adaptive radiation.

## Results

### *Chaetodontinae* is a Late Miocene radiation

We inferred the phylogeny of *Chaetodontidae* from 1,009 ultraconserved element (UCE) loci sampled across 218 species, comprising 257 specimens (Supplementary Table 1) representing 120 of 138 currently recognized species of *Chaetodontidae* (87% species coverage; Eschmeyer’s Catalog of Fishes, accessed October 2026) and 98 outgroup species. *Chaetodontinae*, the focal lineage of this study, is represented by 97 of 110 described species (88%). Bannerfishes (*Heniochinae*) and butterflyfishes (*Chaetodontinae*) are each resolved as monophyletic (Fig. 2). In *Chaetodontinae*, *Prognathodes* is paraphyletic with respect to *Roa* (Fig. 2; Supplementary Fig. 1; Supplementary Information). The phylogeny is largely consistent with previous studies (Supplementary Discussion 1).

**Fig. 2.**
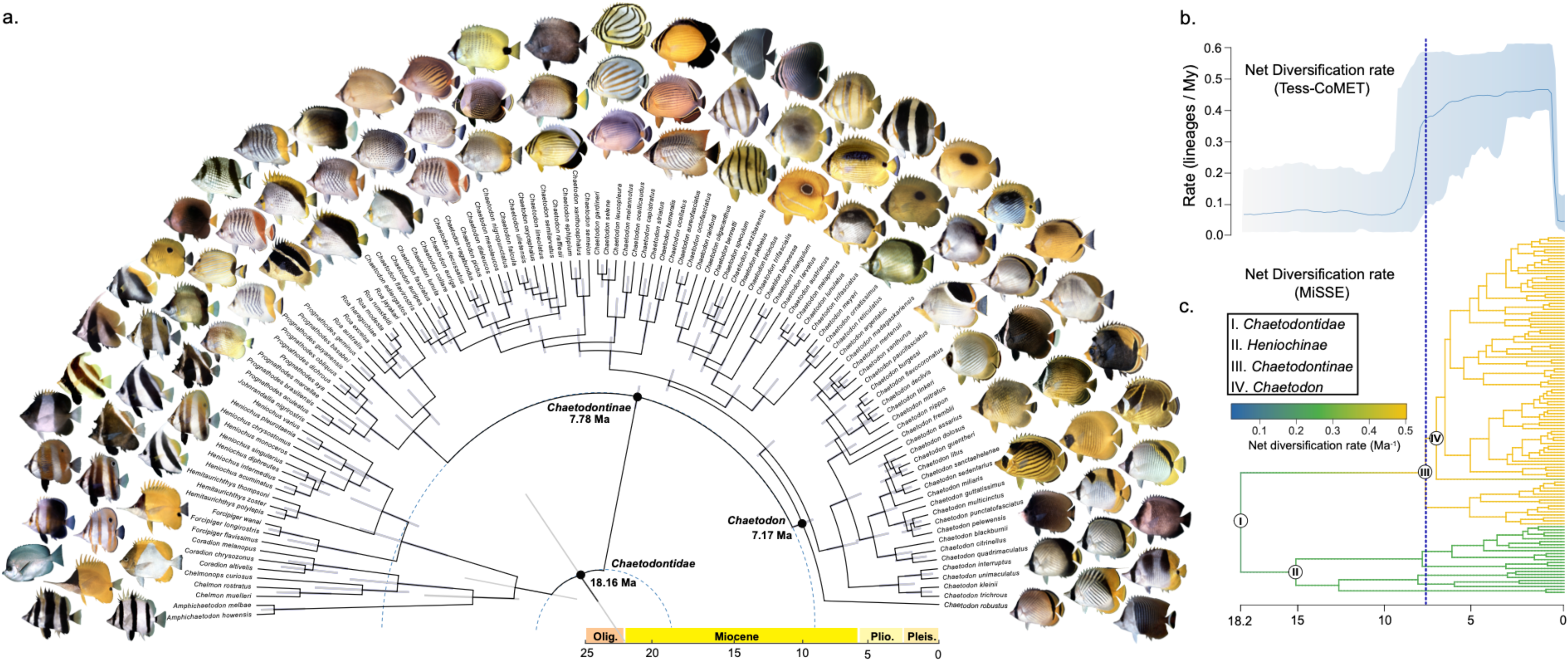
Phylogeny and timescale of butterflyfish diversification. (a) Time-calibrated phylogeny of *Chaetodontidae* (n=120 species), shown as a circular tree with representative species illustrations at the tips, inferred from 1,009 ultraconserved element (UCE) loci and dated in BEAST with the fossilized birth–death model and fossil calibrations. Grey bars are 95% highest posterior density (HPD) intervals of node ages. Black dots and dashed circles mark the crown ages of *Chaetodontidae* (∼18.2 Ma) and *Chaetodontinae* (∼7.8 Ma). The scale is in millions of years (Ma) before present, with geological epochs. (b) Net diversification rate through time estimated with TESS-CoMET (line, posterior mean; shading, 95% credible interval). (c) Net diversification rate along branches from the best-supported MiSSE model (Extended Data Fig. 3, Extended Data Table 2), pruned to *Chaetodontidae*. Dotted vertical lines in (b) and (c) mark the crown age of *Chaetodontinae* (7.78 Ma).

The age of the most recent common ancestor (MRCA) of *Chaetodontidae* is 18.16 Ma (95% highest posterior density [HPD] interval: 12.36–26.95 Ma) (Fig. 2; Extended Data Fig. 1). In addition, we estimated the age of the MRCA of *Chaetodontinae*, which contains most coral-feeding species in the genera *Prognathodes*, *Roa*, and *Chaetodon*, to be 7.78 Ma (95% HPD: 7.13–8.49 Ma) (Fig. 2, Supplementary Tables 2, 3). The age of the MRCA of *Chaetodon* is estimated at 7.17 Ma (95% HPD: 6.48–7.86 Ma) (Fig. 2).

The Early Miocene crown age estimate for *Chaetodontidae* is younger than prior estimates which placed the origin in the Oligocene or Eocene: 28.7 Ma (95% HPD: 18.26– 40.0 Ma)^7^; 32.8 Ma (95% HPD 24.9–40.9 Ma)^11^; 36.72 Ma^23^ and 40.52 Ma^39^. Similarly, our crown age estimate for *Chaetodontinae* (7.78 Ma) is also younger than prior estimates: 15.4 Ma^7^; 23.9 Ma^11^; 24.0 Ma^23^; and 28.10 Ma^39^.

### *Chaetodontinae* diversified exceptionally fast

To place the diversification of *Chaetodontinae* in a broader context of coral reef fish lineages, we estimated crown lineage diversification rates across several reef-associated clades under differing extinction fractions (ε = 0 and ε = 0.9) (Fig. 3a; Supplementary Table 4). Under both extinction fractions, *Chaetodontinae* showed the highest estimated crown diversification rate of all clades examined (0.515 Ma^-1^ at ε = 0 and 0.312 Ma^-1^ at ε = 0.9), exceeding rates for other species-rich reef-fish radiations including *Cirrhilabrus* (fairy wrasses), *Blenniidae* (blennies), *Pomacentridae* (clownfishes and damselfishes)*, Cirrhitidae* (hawkfishes), and *Gobiidae* (gobies) (Fig. 3a; Supplementary Table 4; Supplementary Discussion 2).

**Fig. 3.**
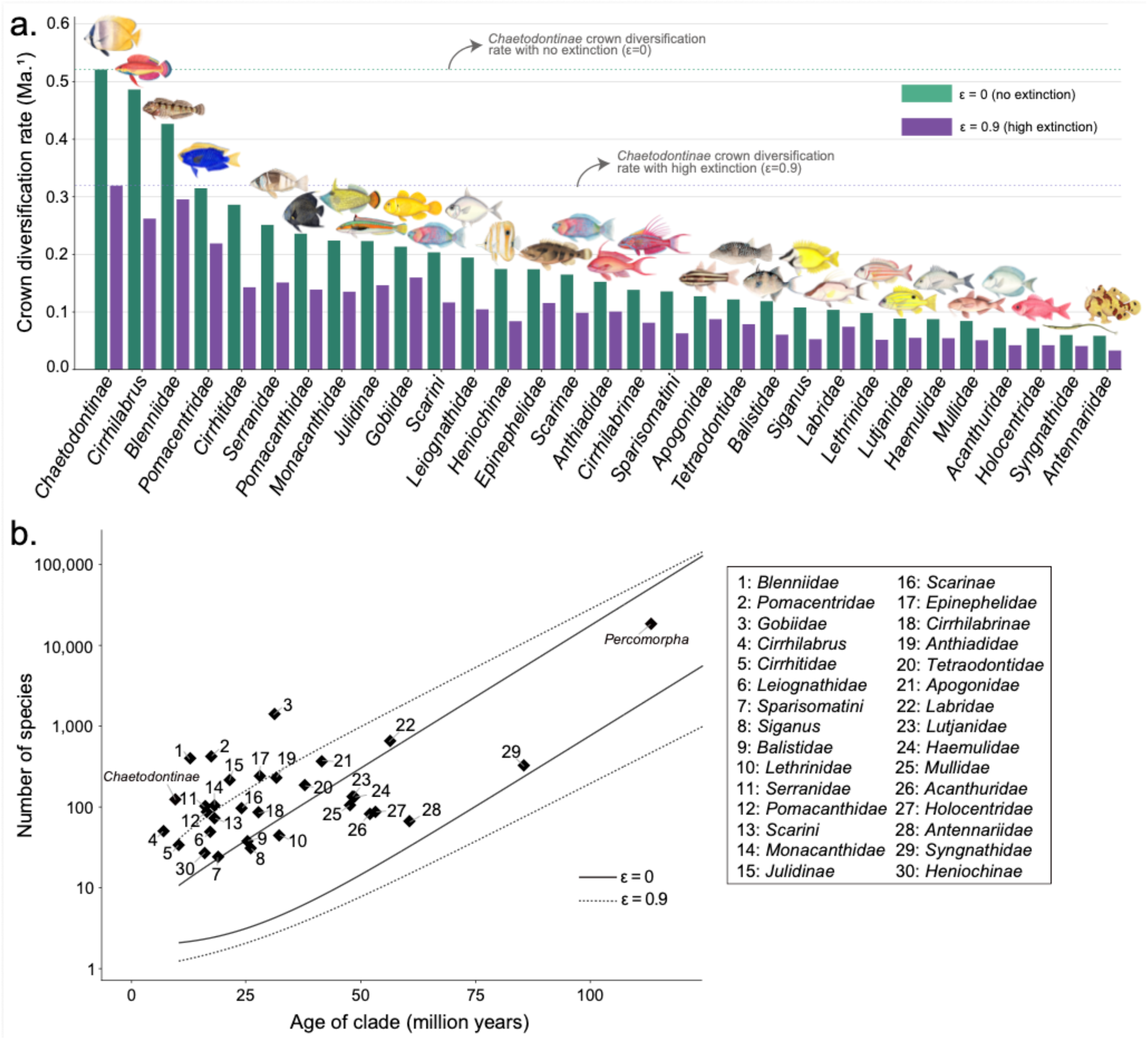
Elevated crown diversification rates in *Chaetodontinae* relative to reef-associated and marine fish lineages. (a) Crown lineage diversification rates for *Chaetodontinae, Heniochinae,* and 29 comparator reef fish lineages, estimated using a crown-age method-of-moments estimator under no extinction (ε=0, teal) and high extinction (ε=0.9, purple), sorted from highest to lowest under ε=0. The dashed horizontal lines mark the estimated lineage diversification rate of *Chaetodontinae* under each extinction scenario. (b) Number of species (log scale) versus crown age for *Chaetodontinae*, *Heniochinae*, the 29 comparator clades, and *Percomorpha*. Solid and dotted lines show the 95% confidence intervals for expected species richness given the background diversification rate of *Percomorpha* at ε=0 and ε=0.9, respectively; points falling above a given line exceed the richness expected for the background lineage diversification rate of *Percomorpha*. See Supplementary Tables 4 and 5a for precise values. Illustrations by Julia Johnson (https://www.lifesciencestudios.com/).

The species diversity of *Chaetodontinae* significantly exceeded the richness expected under the lineage diversification rate of 27 of the 29 comparator reef-fish lineages assuming no extinction (ε = 0: all except *Cirrhilabrus* and *Blenniidae*, the two lineages with the next-highest rates) and 26 of 29 under high extinction (ε = 0.9: all except *Cirrhilabrus, Pomacentridae,* and *Blenniidae*) (Extended Data Table 1).

A time-varying TESS-CoMET analysis reconstructed increased and sustained rates of lineage diversification in *Chaetodontidae*: net diversification stayed near 0.07-0.08 Ma^-1^ (95% CI: −0.06 to 0.19 Ma^-1^) from the MRCA of the lineage until approximately 8 Ma, rose sharply between approximately 8.5 and 7.5 Ma, overlapping with the origin of *Chaetodontinae* (7.78 Ma; 95% HPD 7.13–8.49) and then remained at approximately 0.44–0.47 Ma^-1^ (0.46, 95% CI 0.35–0.59 Ma^-1^ at 2 Ma) until approximately 0.7 Ma (Fig. 2b, Extended Data Fig. 2c). Extinction was low throughout (posterior mean 0.012–0.025 Ma⁻¹; Extended Data Fig. 2b; Supplementary Discussion 2) and poorly constrained. The apparent decline towards the present coincides with the absence of nodes younger than 0.5 Ma (2 ln BF = 5.7 for a speciation-rate shift at approximately 0.7–0.5 Ma), which we interpret as an artifact of the diversification model rather than a real slowdown.

Rate heterogeneity analyses using MiSSE independently localized the increase in diversification to *Chaetodontinae*. Models with rate heterogeneity fitted far better than a constant-rate model (ΔAICc = 27.9–37.0; Extended Data Fig. 3; Extended Data Table 2), and the two best-supported models (cumulative AICc weight = 0.91) both placed all 97 sampled *Chaetodontinae* in a single faster category (0.50 and 0.48 Ma⁻¹) that contains none of the 69 sampled bannerfishes (*Heniochinae*) and outgroup angelfishes (*Pomacanthidae*) (Fig. 2c; Extended Data Fig. 3). The crown lineage of *Chaetodontinae* was assigned to this category (*P* = 0.98 and 0.99) but the crown of the more inclusive *Chaetodontidae* was not (*P* = 0.03 and 0.00), placing a single shift on the branch leading to *Chaetodontinae* that contains species of *Chaetodon*, *Prognathodes*, and *Roa*. Model-averaged tip rates averaged 0.48 Ma⁻¹ in *Chaetodontinae* and 0.25 Ma⁻¹ in the other 69 species of bannerfishes and angelfishes, and every species of *Chaetodontinae* (≥0.40 Ma⁻¹) exceeded every other species of *Heniochinae* and *Pomacanthidae* (≤0.29 Ma⁻¹).

To test whether the elevated diversification rate of *Chaetodontinae* is exceptional relative to other clades, we compared the expected species diversity of *Chaetodontinae*, *Heniochinae*, and 29 other reef fish lineages using the background diversification rates of the inclusive clades *Percomorpha*, *Eupercaria*, *Perciformes*, and *Acanthuriformes*^40^. Under the background diversification rate of *Percomorpha* (r = 0.081 Ma^-1^ at ε=0 and r = 0.066 Ma^-1^ at ε = 0.9), *Chaetodontinae* was more species rich than expected under both extinction scenarios, with observed richness exceeding the upper confidence bound in each case (Fig. 3b, Extended Data Fig. 4, Supplementary Table 5a). Similarly, the species diversity of *Chaetodontinae* exceeded the upper ε=0.9 bound under the background diversification rates of *Eupercaria*, *Perciformes*, and *Acanthuriformes* (Extended Data Fig. 4, Supplementary Table 5b–d).

### Dietary specialization among corallivores

Compilation of *Chaetodontidae* feeding studies shows that dietary composition varies considerably both between non-corallivorous and corallivorous species, and within corallivores themselves^41–45^. Among species classified as obligate coral-feeders, hard scleractinian branching corals dominated the diet: most derived 100% of feeding records from scleractinian corals, with many others exceeding 90% (Fig. 4a, Supplementary Tables 2 and 3), indicating that hard-coral specialization is the dominant strategy among obligate corallivorous butterflyfishes. An exception is *Chaetodon melannotus*, in which soft corals constitute most of the dietary records (Fig. 4a; Supplementary Discussion 3).

**Fig. 4.**
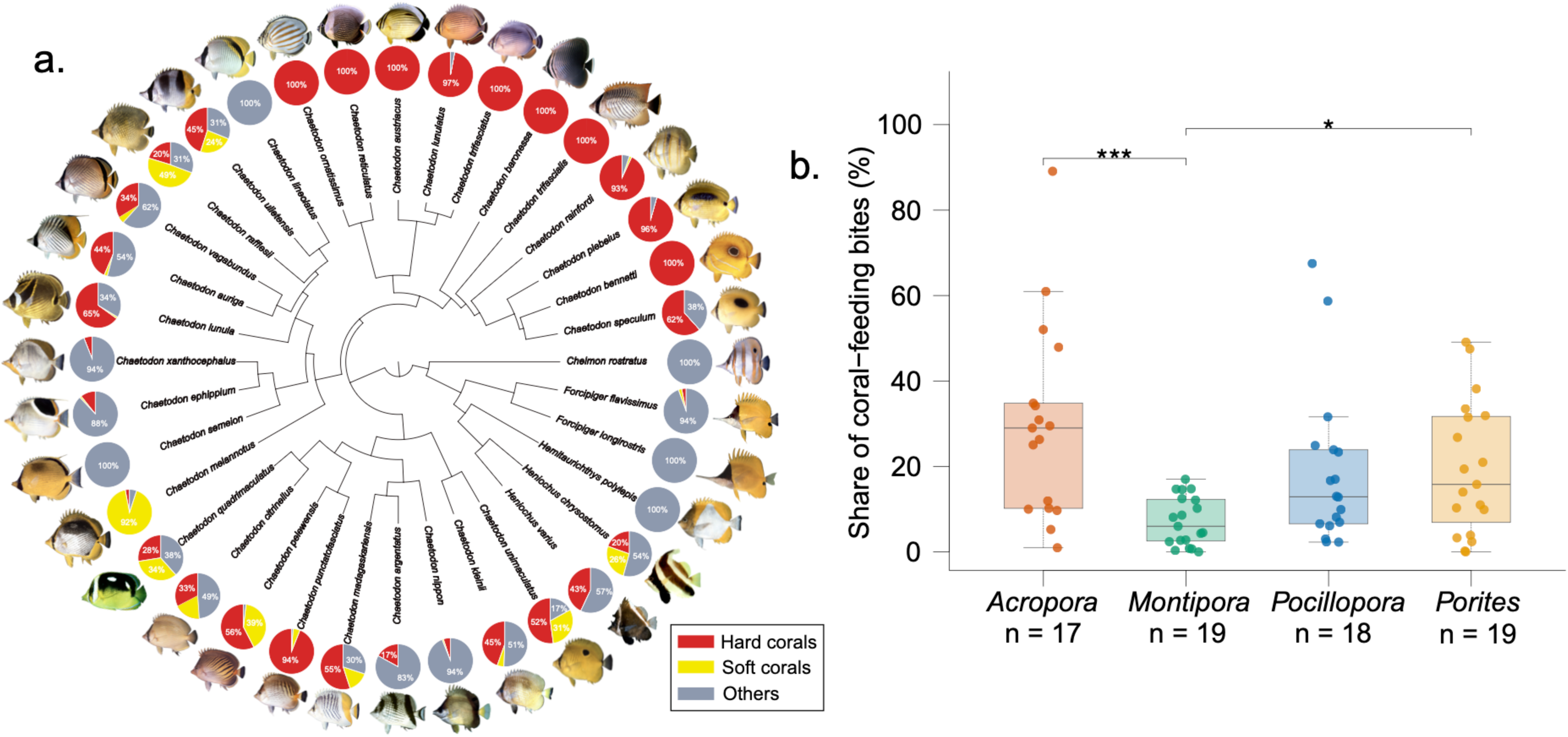
Dietary preferences of corallivorous Chaetodontidae. (a) Phylogeny of 36 *Chaetodontidae* species, with pie charts showing the proportion of dietary items (from bite observations and gut-content analyses) attributed to hard corals (red), soft corals (yellow), and other prey (grey) for each species. (b) Boxplots comparing the proportional use of four genera (*Acropora*, *Pocillopora*, *Porites*, *Montipora*) among obligate corallivores (points = species-level means; sample sizes given per genus). A Friedman test indicated a significant overall difference in genus use (χ²=10.8, d.f.=3, *P*=0.013; Kendall’s W=0.211); among pairwise comparisons, *Acropora* versus *Montipora* (\*\*\**P*<0.001) and *Porites* versus *Montipora* (\**P*<0.05) remained significant after Holm-Bonferroni correction (two-sided Wilcoxon signed-rank tests, indicated by asterisks). See Supplementary Tables 3 and 7.

Friedman’s test revealed a significant overall difference in coral genus use among 19 obligate corallivore species (χ² = 10.8, df = 3, p = 0.013), though the effect size was modest (W = 0.211, n = 17) (Fig. 4b). Of the six pairwise contrasts, two remained significant after Holm-Bonferroni correction: *Acropora* was consumed at higher rates than *Montipora* (*P*_adj_=0.00064), and *Porites* at higher rates than *Montipora* (*P*_adj_ = 0.010); the remaining four contrasts were non-significant (Fig. 4b).

### Asymmetric transitions in diet, depth and jaw innovation

To test whether the reconstructed shifts in lineage diversification in *Chaetodontidae* (Fig. 2b,c) coincided with evolution of ecologically relevant traits, we reconstructed ancestral states of dietary mode, the IMJ, depth category, and dentition on the time-calibrated phylogeny (Fig. 5, Extended Data Figs 5 and 6), and tested whether habitat depth is phylogenetically structured and whether corallivory is associated with habitat depth, as expected for consumers of light-dependent corals confined to shallow reefs.

**Fig. 5.**
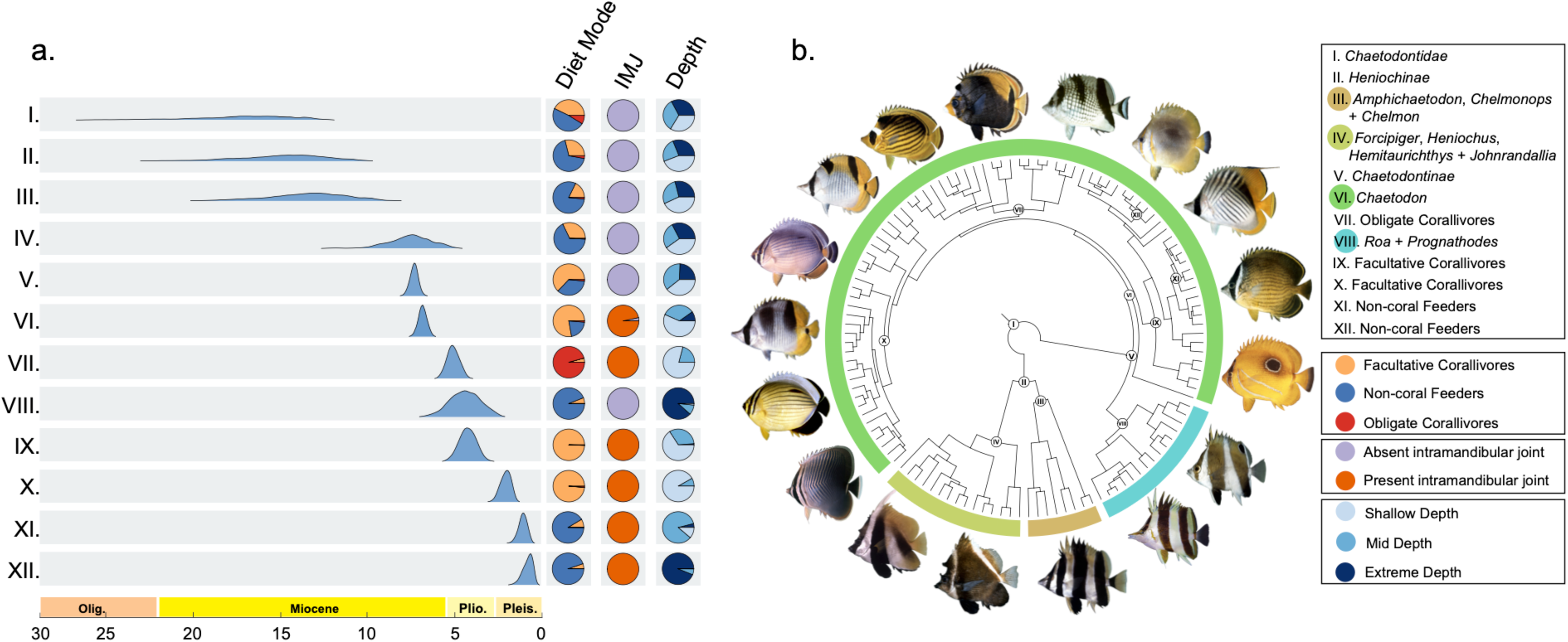
Ancestral state reconstruction of diet, jaw morphology and depth use across Chaetodontidae. (a) Posterior distribution of relaxed clock node-age estimates (Miocene-Pleistocene) for 12 focal nodes (I-XII), each paired with pie charts showing the reconstructed ancestral-state probabilities at the node for diet mode (non-coral feeder, facultative corallivore, obligate corallivore), intramandibular joint (IMJ) (absent, present), and depth category (shallow, mid, extreme) at the node. (b) Phylogeny of *Chaetodontidae* with the 12 focal nodes labeled and color-coded corresponding to the taxonomic grouping defined in the legend. See Supplementary Table 2 for information regarding how each species was classified.

Diet evolution favored an all-rates-different model (ARD; AIC = 160.51). The ancestors of *Chaetodontinae* and of *Chaetodon* were reconstructed as facultative corallivores (*P* = 0.63 and 0.78; Extended Data Fig. 5a; Supplementary Discussion 3), although the root state of *Chaetodontidae* was unresolved (posterior probabilities from stochastic mapping: non-coral feeding 0.49, facultative corallivory 0.43, obligate corallivory 0.09). Obligate corallivory occurred only within *Chaetodon* and arose at least four times (parsimony; stochastic mapping mean 4.7). Obligate corallivory is first reconstructed at the crown of a coral-feeding clade that includes 17 of the 23 species of obligate corallivores (node VII; 5.63 Ma, 95% HPD 4.83–6.34; *P* = 0.96). Notably, stochastic maps reconstructed no direct transitions from non-coral feeding to obligate corallivory; obligate corallivory was always entered through facultative corallivory, whereas facultative corallivory has a history of both losses (mean 16.6 transitions to non-coral feeding) and regains (mean 5.3 transitions from non-coral feeding).

The reconstructed history of IMJ evolution favored an irreversible gain-only model (GainOnly; AIC = 16.45). The ancestral state for the MRCA of *Chaetodontidae* was reconstructed as IMJ absent (*P* = 1.00), with a posterior mean of 1.03 independent gains across the phylogeny. Branches with IMJ present accounted for 54% of total tree length, representing all sampled species of *Chaetodon,* and all obligate corallivore clades ancestrally possessed an IMJ (Extended Data Fig. 5b). This reconstruction carries some uncertainty, as the IMJ has been scored in only a small number of species of *Chaetodontidae*^29^.

Habitat depth category was highly labile (mean = 128.47 changes), and non-coral feeders were concentrated in deep environments (26 of 53 species), whereas obligate corallivores were concentrated in shallow environments (16 of 23; Extended Data Fig. 5c). Depth showed significant phylogenetic signal when treated as a continuous trait (Pagel’s λ = 0.701, p = 4.2×10^−12^; Supplementary Fig. 2). Phylogenetic logistic regression confirmed a significant negative association between corallivory and depth (β = −0.0137 ± 0.0059 per m, p = 0.020), consistent with corallivory being confined to the shallow reef zones where light-dependent corals occur; the association was in the same direction but non-significant when depth was log10-transformed (β = −1.02 ± 0.57, p = 0.073).

We additionally reconstructed the evolutionary history of jaw and tooth morphology, coding this character following an osteological scheme for the *Chaetodontidae* dentition^46^ (Extended Data Fig. 6; Supplementary Discussion 3). Under an equal-rates transition model, the ancestral state at the most recent common ancestor of *Chaetodontidae* was reconstructed as having a basic dental pattern and lacking each of the derived dental character states (*P* ≥ 0.969 in every case). The simpler derived states (18:2–18:4) occur only in facultative corallivores or non-coral feeders outside *Chaetodon*, whereas the three more complex, coral-associated states (18:5–18:7) (teeth organized in more narrow and numerous bands, teeth straight with descending process edentulous, and very fine teeth coalescing into a brush) are found almost exclusively in species of *Chaetodon* that are obligate corallivores (Extended Data Fig. 6). The common ancestor of the coral-feeding clade of *Chaetodon* (node VII, Fig. 5), at which obligate corallivory is first reconstructed, is also reconstructed as having complex coral-associated teeth (*P* = 0.99; Extended Data Fig. 6).

### Morphology tracks depth more than diet

To test whether phenotypic diversification is tied to ecological differentiation, we quantified morphological variation and asked whether it covaries with depth and diet. Body shape covaried with the combined depth-and-diet ecology (two-block partial least squares, full trait set versus depth and diet: r = 0.390, permutation *P* = 0.008, n = 82; body traits: r = 0.398, *P* = 0.005; jaw traits: r = 0.264, *P* = 0.152), and depth rather than diet accounted for the association (phylogenetic generalized least squares with depth alone: R²=0.130, P<0.001 for all traits; with corallivory alone: all P>0.25; Extended Data Fig. 7 and Extended Data Table 3). Lower jaw length and mouth width evolved faster than any body-shape trait (Brownian-motion σ²=0.039 and 0.036 versus 0.0018–0.0092; Supplementary Table 6; Supplementary Discussion 4).

## Discussion

Our time-calibrated phylogenomic analyses indicate that *Chaetodontinae* comprises an exceptionally young and rapidly diversifying radiation that is consistent with an adaptive radiation framework in an open-marine system. We estimate a Late Miocene age of 7.78 Ma (95% HPD: 7.13–8.49 Ma) for *Chaetodontinae*, implying that 110 species, 80% of the species diversity of *Chaetodontidae*, have accumulated within the past 8 million years. In standardized richness–age comparisons across 29 reef-fish lineages and across progressively broader background clades, *Chaetodontinae* is consistently more species rich than expected, placing it among the fastest-diversifying reef-associated clades under a range of extinction assumptions (Fig. 3; Extended Data Fig. 4).

Although definitions of rapid diversification vary^5,47,48^, we assess diversification comparatively. On a standardized richness–age scale, *Chaetodontinae* is an outlier across reef-fish clades (Fig. 3). The diversification of the 110 species of *Chaetodontinae* within 7.78 Ma implies unusually fast lineage diversification, comparable to rates reported for several canonical adaptive radiations^37^. Importantly, we do not interpret the comparison clades as non-radiating nulls; several are themselves widely regarded as reef-fish radiations, and our comparison is instead intended to place *Chaetodontinae* on a common richness–age scale.

TESS-CoMET estimates low net diversification for roughly the first 8 Myr following the origin of *Chaetodontidae*, followed by a sharp increase and sustained high rate of lineage diversification that coincides with the origin of *Chaetodontinae* (Fig. 2b). Rate heterogeneity analyses using MiSSE assigned the highest-rate category to every sampled species of *Chaetodontinae* and to none of the 69 sampled bannerfishes and angelfishes, with a single shift on the branch leading to *Chaetodontinae* (Fig. 2c). The MiSSE estimated rate of lineage diversification (0.50 Ma⁻¹) is almost identical to the crown-group rate of *Chaetodontinae* from the method-of-moments estimator (0.515 Ma⁻¹ at ε = 0) and about twice that of *Chaetodontidae* as a whole (0.233 Ma⁻¹); three independent methods built on different assumptions agree that *Chaetodontinae* exhibits an exceptionally high rate of lineage diversification.

This pattern is consistent with the expectation that adaptive radiations may be temporally heterogeneous: young clades can exhibit elevated net diversification while ecological opportunity is abundant, whereas older clades may show reduced apparent net diversification after niche space fills, diversification slows, or extinction erodes earlier pulses of speciation^15,49^. Under this interpretation, the high species richness of *Chaetodontinae* given its relatively young Late Miocene crown age reflects a radiation that is recent and potentially still unfolding, nested within the older *Chaetodontidae* and sister to the more slowly diversifying *Heniochinae*, rather than implying that older clades in our comparison lacked adaptive radiations.

Interpreting this signature of adaptive radiation in *Chaetodontinae* is aided by historical context. The Early–Mid Miocene (23–11.6 Ma; peak 16–15 Ma) was marked by the largest increase in reef area and thickness of the past 66 million years^21,50^, when the Great Indo-Australian Miocene Reef System exceeded modern reefs in both seafloor area and thickness^51,52^ and coral lineages diversified across most growth forms^21^ (Supplementary Discussion 1). Notably, branching corals that dominate many present-day Indo-Pacific reefs, including *Acropora*, the genus taking the largest median share of bites in obligate corallivorous butterflyfishes (Fig. 4b), began to diversify later, in the Late Miocene and Pliocene (11.63-2.58 Ma)^8,21,53^. The rise of fast-growing, structurally complex branching corals likely created a rapidly diversifying and underexploited resource base capable of promoting trophic specialization^11^.

The crown age for *Chaetodontidae* (18.16 Ma, 95% HPD: 12.36–26.95 Ma) aligns with the onset of the earlier, early-mid Miocene reef-expansion phase, placing the origin of the lineage alongside that of other major reef-fish clades that radiated during the Early–Mid Miocene^19,20^. By contrast, our estimated crown age for *Chaetodontinae* (7.78 Ma, 95% HPD: 7.13–8.49 Ma) overlaps with the early portion of the Late Miocene reef reorganization and the onset of branching-coral diversification, and this temporal overlap is consistent with ecological turnover and the diversification of branching corals that created new niche space for coral-associated consumers, thereby contributing to rapid lineage diversification within *Chaetodontinae*. Trait reconstructions are consistent with this scenario. The most recent common ancestors of *Chaetodontinae* and *Chaetodon* were reconstructed as facultative corallivores (*P* = 0.63 and 0.78; Fig. 5), consistent with coral feeding emerging at the origin of *Chaetodontinae* and intensifying later within *Chaetodon*, and obligate corallivory arising only through facultative corallivory (Fig. 5). This contrasts with a previous reconstruction that resolved the root of *Chaetodontidae* as a facultative corallivore^54^; the difference likely reflects our denser taxon sampling and acanthuriform-specific fossil calibrations, which together shift the crown age of *Chaetodontinae* substantially younger and alter the relative branch lengths on which the reconstruction is based.

Independent trait-level evidence from depth distributions supports this ecological opportunity hypothesis. Phylogenetic logistic regression shows a significant negative association between corallivory and habitat depth, and the ancestral state reconstruction places the origin of obligate corallivory within shallow-water lineages of *Chaetodon* (*P* = 0.79; Fig. 5). This depth restriction mirrors the ecology of their primary resource: scleractinian branching corals are concentrated in shallow, well-lit reef zones and decline sharply in cover with depth^55,56^. The association between corallivory and shallow depth, the phylogenetic structuring of depth across *Chaetodontidae*, and the concentration of obligate corallivores in shallow reef zones converge on the same conclusion: the ecological opportunity that promoted diversification within *Chaetodontinae* was shallow, branching-coral reef habitat, the same habitat now most severely affected by thermal bleaching and the ongoing decline of branching *Acropora* and related genera.

Among candidate evolutionary innovations, the intramandibular joint (IMJ) is a plausible key innovation: adaptive radiation^15^ theory expects such traits to increase performance in a newly exploited habitat or resource. Although we did not measure performance, the IMJ bears on the performance-utility criterion for ecological adaptive radiation because earlier work links this discrete phenotypic change to a shift in feeding mechanics and resource use^29,30^ (Supplementary Discussion 5). The IMJ arose once, in the common ancestor of *Chaetodon*, and is absent from every other chaetodontid lineage, whereas obligate corallivory evolved later and only within a derived set of species of *Chaetodon*. The joint therefore appears to have functioned as a pre-adaptation in *Chaetodon*, reconstructed as present in the MRCA of the clade (Fig. 5, Extended Data Fig. 5b), but with its ecological potential only fully realized once particular lineages specialized on hard coral. Its gain-only pattern in ancestral character reconstruction is consistent with a key innovation model (Fig. 5), in which access to a resource or foraging mode reduces the selective incentive to revert to an ancestral, less effective configuration^57^. The gain-only pattern of the IMJ is consistent with a previous study^58^ that showed that chaetodontid lineages transitioning toward zooplanktivory evolve narrower bodies and a modified caudal fin associated with sustained, open-water swimming and the pursuit of fast-moving planktonic prey, which likely functionally constrain these species from transitioning back to coral-feeding.

The IMJ is not the only morphological feature consistent with specialization on coral-derived resources: derived tooth morphologies, including long thin teeth organized into pads and peripheral placement of robust teeth facilitating forceful biting^33,54^ and elongation of the digestive tract associated with coral-based diets^34^ (Fig. 1), further document functional differentiation aligned with hard-coral feeding. Our tooth reconstructions show that the most derived dental states (18:5–18:7) occur almost exclusively in obligate corallivores within *Chaetodon* (Extended Data Fig. 6), consistent with an earlier osteological analysis of *Chaetodontidae*^46^ and reinforcing the conclusion that hard-coral specialization is a derived capacity within *Chaetodon* rather than an attribute of *Chaetodontidae* as a whole.

Butterflyfishes (*Chaetodontinae*) meet key criteria of adaptive radiation^14,15,49^: a monophyletic clade that arose in the Late Miocene (7.78 Ma) and diversified faster than nearly all other reef-fish lineages, even under high extinction, with obligate corallivory arising repeatedly within *Chaetodon* from facultative ancestors after the single gain of the intramandibular joint^29,57^ and alongside derived tooth morphologies^31–33,54^. This radiation coincides with the Late Miocene reorganization of reef coral assemblages and the onset of branching-coral diversification^8,21,53^, suggesting that adaptive radiation can proceed on coral reefs, where opportunity arises from reef restructuring within an open seascape rather than from geographic isolation or post-extinction release. If the rise of branching corals helped generate this diversity, their ongoing loss^59–61^ may be dismantling the resource base on which butterflyfishes and other coral-dependent lineages rely.

## Methods

### Nomenclature

In this study, we follow a rank-free phylogenetic taxonomy of ray-finned fishes^40^, which follows the convention of the *PhyloCode*^62^. All formal taxonomic names, regardless of rank in the Linnaean system are italicized.

### Ultraconserved element dataset assembly

We extracted DNA from 257 specimens comprising 120 species in *Chaetodontidae* and sent samples to Arbor Biosciences for sequencing. We used Phyluce 1.7.3^63^ to align the contigs, extract ultraconserved elements using the probe set^64^, and produce a 75% complete matrix comprising 1,009 ultraconserved elements sampled across 218 species, including 120 species of *Chaetodontidae* (23 *Heniochinae* and 97 *Chaetodontinae*, of which 82 are *Chaetodon),* and 98 outgroups. The outgroup taxa include *Luvarus imperialis*, *Zanclus cornutus*, and species of *Acanthuridae*, *Leiognathidae*, and *Pomacanthidae*, which are resolved with Chaetodontidae in an inclusive lineage *Acanthuriformes* across multiple phylogenetic analyses^65–69^.

### Phylogenetic analyses and time calibration

We performed maximum-likelihood analyses of the 75%-complete dataset using IQ-TREE2^70^ that included all sampled specimens of *Chaetodontidae* and outgroup taxa. We calibrated the phylogeny of *Chaetodontidae* using a Bayesian node-dating approach implemented in BEAST v. 2.6.7^71,72^. First, for each sampled species of *Chaetodontidae* we included a single specimen and randomly partitioned 150 UCE sequence alignments into three sets of 50 UCEs, each of which we ran as an independent BEAST analysis. We used the log-normal relaxed clock model, and the BEAST2 implementation of the Fossilized Birth Death Model branching model^73,74^. We set the rho sampling parameter to 0.543, which is the proportion of living species in the most inclusive clade that includes the sampled outgroup taxa^70^, set the diversification rate prior to 0.05 (the approximate background diversification rate for *Acanthomorpha* fishes estimated in ^69^) with bounds of 0.0 and 1.0, and set the origin prior to 83.6 Ma, with bounds of 125.0 Ma and 66.02 Ma.

We estimated our time-calibrated phylogeny using 8 fossil node calibrations with the fossilized birth-death branching model (Extended Data Fig. 1; Supplementary Information). The ages of fossils and justifications for their placement in the phylogeny are in the Supplementary Information. For each fossil calibration (Supplementary Information), we adjusted the log-normal prior so that 97.5% of the distribution fell before the assigned age^75^. We fixed the tree topology to the phylogeny that we inferred by treating the UCEs as a single concatenated alignment with a single-partition scheme, and conducted two independent Markov Chain Monte Carlo (MCMC) runs for each set of 50 UCEs, each over 2.0 × 10^8^ generations with a 1.0 × 10^7^ burn-in. After confirming convergence of the posteriors and effective sample size values of 200 or more in Tracer v. 1.7.1^76^, we combined the top 25% of each posterior tree set (subsampling every 5000 generations) in LogCombiner v.2.6.6 and annotated the target tree (the phylogeny inferred in IQ-TREE2 using a single partition model on the concatenated UCE dataset) with median node heights in TreeAnnotator v.2.6.6^71,72^. The combined posterior sample contained 106,119 trees. To obtain posterior distributions of clade ages, we recorded the age of the most recent common ancestor of each named clade in every tree of this sample after discarding the first 10% of trees (95,508 trees retained). All comparative analyses used the single annotated target tree and therefore do not propagate uncertainty in divergence times.

### Diversification rates

We estimated the rate of lineage diversification for *Chaetodontinae*, its sister lineage *Heniochinae,* and 29 other lineages of reef fishes including *Blenniidae*, *Pomacentridae*, *Pomacanthidae*, *Monacanthidae*, *Julidinae*, *Leiognathidae*, *Gobiidae*, *Scarinae*, *Tetraodontidae*, *Balistidae*, *Mullidae*, *Syngnathidae*, *Apogonidae*, *Labridae*, *Scarini*, *Sparisomatini*, *Cirrhilabrinae*, *Cirrhilabrus*, *Anthiadidae*, *Epinephelidae*, *Serranidae*, *Acanthuridae*, *Holocentridae*, *Cirrhitidae*, *Lethrinidae*, *Lutjanidae*, *Haemulidae*, *Siganus*, and *Antennariidae*. Specifically, we estimated the crown-group net diversification rates for each lineage individually in the context of a birth-death model of diversification using the method-of-moments estimator of Magallón and Sanderson (2001; equation 4)^77^. Comparison clades were chosen because they are reef-associated^16^, have standing richness sufficient for stable richness–age comparisons, and are represented in published time-calibrated phylogenies with crown-age estimates; collectively, they span a range of trophic strategies and habitat affinities among percomorph reef fishes. Comparison clades are not intended to represent all reef-fish diversity, but to provide a broad, phylogenetically and ecologically diverse baseline of well-sampled reef-associated lineages. More specifically, we derived net diversification rate from each clade’s crown age, standing species richness, and an assumed relative extinction fraction (ε)^77^. Crown-age estimates were obtained from either our relaxed clock analysis or from previous studies (Supplementary Table 4). The total described species richness for each group was obtained mostly from Eschmeyer’s Catalog of Fishes^78^ (accessed June 2026), and rates were calculated under both a pure-birth (ε = 0) and high-extinction (ε = 0.9) scenario using bd.ms() in the R package geiger^79,80^. To formally test whether *Chaetodontinae’s* observed species richness exceeds that predicted under a comparison clade’s diversification rate, we used crown.limits() (geiger) to compute the 95% confidence interval on expected richness for a clade with the crown age estimate of *Chaetodontinae’s* diversifying at each of the comparison clade’s rate, under both extinction scenarios.

We explored the expected species richness of *Chaetodontinae, Heniochinae,* and 29 other lineages of reef fishes given the background rate of diversification of the inclusive lineages *Percomorpha* (*N* = 19,110 species; crown age = 113.0 Ma), *Eupercaria* (*N* = 7,134 species; crown age = 87.7 Ma), *Perciformes* (*N* = 3,213 species; crown age = 51.1 Ma), and *Acanthuriformes* (*N* = 2,423 species; crown age = 80.9 Ma) following the approach presented by Magallón and Sanderson^16,77^. We calculated each lineage’s diversification rate using equation 4 in Magallón and Sanderson^77^, and we calculated the 95% confidence interval on expected species richness under the estimated diversification rate through time under a birth-death process diversifying at that rate, using the crown.limits() (geiger) (Cl = 0.95). We then plotted *Chaetodontinae* and the other 29 lineages against these two confidence bounds as a function of crown age, for each of the inclusive lineages. Following Magallón and Sanderson^77^, we classified lineages falling above the upper bound of the high extinction scenario (ε = 0.9) as containing more species than expected given the background lineage diversification rate of *Percomorpha*. We also classified lineages falling below the lower bound of the same interval as fewer than expected number of species. We estimated lineage diversification for *Chaetodontidae*^7^ using TESS-CoMET^81^, a Bayesian episodic birth–death model in which speciation and extinction rates change at times shared by all lineages and mass extinctions can occur. We set the sampling probability to 0.87 based on the proportion of *Chaetodontidae* species sampled relative to the total valid species count for the *Chaetodontidae* clade in Eschmeyer’s Catalog of Fishes (Accessed June 2026)^78^. We set the expected survival probability to 0.90, reflecting minimal extinction in the Cenozoic teleost fish record^82–84^ and the expectation that lineage loss within *Chaetodontidae* has been negligible relative to speciation over the family’s evolutionary history, with one expected mass extinction event specified to avoid being overly restrictive. Speciation and extinction rates had lognormal priors with means of 0.10 and 0.02 (standard deviation 0.05 for both), and ten rate changes were expected *a priori*. We ran MCMC for up to 2.0 × 10^6^ iterations with a burn-in of 3.0 × 10^4^ generations using an adaptive proposal mechanism. The run stopped automatically once its effective-sample-size criterion was met, after 63,000 iterations for *Chaetodontidae*. Samples were stored every 100 iterations, the first 25% of stored samples were discarded, and rates were summarized over 100 time intervals. Support for rate changes and mass extinctions was assessed with Bayes factors (2 ln BF thresholds of 2, 6 and 10). Convergence of the posterior distribution was confirmed both by visual inspection of the trace plot and by effective sample size (ESS), which exceeded 330 for all monitored parameters (log-posterior ESS = 336; number of speciation-rate categories ESS = 473; number of extinction-rate categories ESS = 473; number of mass extinctions ESS = 473).

To test for heterogeneity in diversification rates that is independent of the traits we measured, we fitted MiSSE models^85^ in the R package hisse to the *Chaetodontidae* tree together with its closely related lineage of *Pomacanthidae* (166 tips: 120 *Chaetodontidae* and 46 *Pomacanthidae*), so that shifts within *Chaetodontidae* can be detected relative to close relatives. We used a single global sampling fraction of 0.73 (166 of 228 described species: 138 *Chaetodontidae* and 90 *Pomacanthidae*). We fitted the six-model set produced by MiSSEGreedy(), which allows turnover and the extinction fraction to vary among one to three hidden rate categories (generateMiSSEGreedyCombinations() with max.param = 4 and vary.both = TRUE), and compared the models, including the constant-rate null, by AICc. We reconstructed the marginal probability of each hidden rate category on every branch with MarginReconMiSSE() and plotted the net diversification rates of the best-supported model.

### Coral diet composition

For 36 of the *Chaetodontidae* species represented in the phylogeny, dietary composition data were compiled from the published literature, drawing specifically on gut-content analyses and field observations^41–44,86^. Where more than one source reported data for a given species, the percentage composition of each prey category was averaged across studies. Prey categories reported in the original sources were then collapsed into three broad groups: hard corals (*Scleractinia*), soft corals, and other prey (including planktivorous and non-coral invertebrate prey items). The resulting per-species dietary proportions are displayed as pie charts positioned at the corresponding tip of the phylogeny (Fig. 4a; Supplementary Table 3).

To further understand the coral genus preferences of obligate coral-feeding butterflyfishes, data for 19 species were compiled^26^ from a report of the proportion of coral-feeding bites taken from *Acropora*, *Montipora*, *Pocillopora*, and *Porites*. For each butterflyfish species, values were averaged across all reported locations to yield a single mean per combination of butterflyfish species and coral genus, avoiding pseudoreplication from butterflyfish species sampled at multiple sites. Differences in mean proportional use among genera were tested with a Friedman test^87^ on the species with a value for all four coral genera (17 of the 19 obligate species). To quantify the effect size, we used Kendall’s coefficient of concordance (W)^88^, indicating how consistently butterflyfish species agree on the relative ranking of the four genera. Pairwise comparisons between each genus pair were then conducted using two-sided Wilcoxon signed-rank tests^89^ on all species with data for both genera. Because six pairwise comparisons were run, p-values were adjusted for multiple testing using the Holm-Bonferroni method^90^. All analyses were performed in R (version 4.5.0), using the friedman.test, wilcox.test, and p.adjust functions from the base stats package, with statistical significance set at α = 0.05 (Supplementary Table 7).

### Ancestral state reconstruction

We reconstructed evolutionary histories of three traits across the time-calibrated phylogeny: the presence and absence of the intramandibular joint (IMJ)^29,30^; the feeding mode, coded as a three-state trait: non-coral feeders; facultative corallivores (≥5% of the diet from coral); and obligate corallivores (approximately >95% of the diet from coral)^26,28,41–44,86,91,92^; and depth, binned from average depth occurrence^93^ into three categories (shallow, equal to or less than 20 meters; mid, 21–50 meters; and extreme, greater than 50 meters). For each discrete trait we fit four candidate continuous-time Markov models; equal-rates (ER), all-rates-different (ARD), and two directional models constraining transitions to occur in a single direction only. We used fitMk() in R package phytools^94^, selected the best-supported model by AIC, and used it to estimate marginal ancestral state probabilities at internal nodes via ace() in ape^95^. Evolutionary trajectories were visualized using stochastic character mapping (1,000 simulations per trait)^96,97^ via make.simmap() in phytools, from which we summarized the posterior mean number of state changes and the marginal probability of each state at the root. For average depth as a continuous trait, ancestral states were additionally estimated under a Brownian motion model using fastAnc() in phytools.

To determine whether depth occurrence is phylogenetically structured, and thereby justify the use of phylogenetically corrected methods in subsequent depth-related analyses, we quantified phylogenetic signal in average depth using Pagel’s λ^98^, implemented via phylosig() in the R package phytools^94^. Significance was evaluated with a likelihood-ratio test comparing the fitted model against a model with λ constrained to zero.

### Phylogenetic logistic regression

To test whether corallivory (facultative or obligate versus non-coral feeding) varies with depth, we fitted a phylogenetic logistic regression with phyloglm() in the R package phylolm^99^ with average depth entered in meters and, because depth is right-skewed, as log10(depth). To test whether corallivory is associated with morphology, we fitted a separate univariate model for each of nine log-transformed measurements (total weight, standard length and the seven linear traits; Supplementary Table 8). Each model was fitted to the species with data for both variables and to the tree pruned to those species. All eleven models converged with the default setting, although the raw-depth model reported that the boundary of the linear predictor had been reached. P-values for the nine morphological predictors were adjusted within each response with the Holm–Bonferroni method.

### Tooth morphology

To assess evolutionary changes in dentition among *Chaetodontidae*, we reconstructed the evolutionary history of seven binary tooth characters described by Blum (1988) (Characters 18:1–18:7)^46^. We used only species for which the cleared and stained specimens were examined and excluded species that had no data or only had radiographs examined^46^.

The finalized character matrix covered 55 *Chaetodontidae* species, most of which were species of *Chaetodon*. We reconstructed all seven characters independently. To compare dentition with diet at the focal nodes of Fig. 5, we also assigned each species to one of four tooth classes (complex, if it carried any of 18:5–18:7; otherwise simple 18:2/18:3; otherwise simple 18:4; otherwise basic only) and reconstructed the class under the same equal-rates model. For each character, maximum-likelihood ancestral states at internal nodes were estimated under an equal-rates (ER) transition model using the ace() function (ape)^95^; an equal-rates model was used given the sparse and often strongly skewed distribution of several of these characters (present in only a small number of species), for which a more heavily parameterized asymmetric-rates model risked being poorly identified. For each character, feeding strategy (facultative corallivore, obligate corallivore, or non-coral feeder) was additionally displayed alongside the reconstructed tooth-character states on the same tree, to allow visual comparison between dental morphology and diet.

### Morphospace and ecospace analysis

To evaluate whether morphological diversification tracks ecological diversification, following the approach of similar papers^100^, we constructed morphospaces from the seven size-corrected morphological traits. Traits were log-transformed and size-corrected as the residuals of a regression on log standard length, using the 82 species with a complete set of measurements. The traits were analyzed separately as (i) all combined, (ii) jaw traits only (head depth, lower jaw length, mouth width), and (iii) body traits only (body depth, fish width, caudal peduncle depth, caudal peduncle width). Principal component analysis (PCA) was used to visualize morphospace occupation (Extended Data Fig. 7a).

Association between each morphospace and an ecology was tested using two-block partial least squares analysis (2B-PLS)^101^, with the cross-covariance matrix between blocks decomposed by singular value decomposition and significance assessed by permutation (999 iterations). We first tested each morphospace against a combined ecospace comprising average depth and an ordinary corallivory score (0 = non-coral feeder, 1 = facultative corallivore, 2 = obligate corallivore). In addition, to determine whether depth or diet was independently responsible for any observed association, we subsequently repeated the PLS procedure with depth and corallivory score tested as separate single-variable ecology blocks against each of the three morphospaces, rather than combined. In all cases, resulting PLS scores were then analyzed by phylogenetic generalized least squares (pGLS), implemented via pgls(), in the R package caper^102^, with λ estimated by maximum likelihood, to account for phylogenetic non-independence among species.

### Rate heterogeneity among morphological traits

To characterize rate heterogeneity in morphological evolution, we estimated the Brownian motion rate parameter (σ²) for each of seven size-corrected, log-transformed linear morphological traits (maximum body depth, maximum fish width, head depth, lower jaw length, mouth width, minimum caudal peduncle depth, minimum caudal peduncle width) via fitContinuous() in the R package geiger^79,80^. Morphological traits were extracted from previously published data^103^, which provided coverage for 82 of the 120 Chaetodontidae species in our phylogeny.

## Data availability

The UCE sequence data generated in this study are deposited in NCBI [database and accession numbers to be confirmed]; specimen vouchers, localities and accession numbers are listed in Supplementary Table 1. Sequence alignments, time-calibrated trees, trait datasets and the data underlying all figures and tables are available at [Dryad/Zenodo DOI to be added].

## Code availability

The R scripts used for all analyses are available at [to be added].

## Acknowledgements

The authors thank members of the Near and Muñoz Labs for discussions related to this manuscript, as well as the following people and museum collections for their assistance and generosity with tissue samples or genetic material: C. Atta, R. Pyle, C. Ka’apu-Lyons Copus, and the Bernice Pauahi Bishop Museum (Honolulu); D. Bray, and the Museum of Victoria; D. Bellwood; B. Bowen; M. Erdmann; A. Friedlander; S. Longo; C. McMahan, M. Westneat, and the Field Museum of Natural History (Chicago); D. Pitassy, K. Murphy, J. Williams, C. Baldwin, and the Smithsonian National Museum of Natural History (Washington, DC); C. Rocha, L. Rocha, M. Hoang, and the California Academy of Sciences (San Francisco); H. Motomura and the Kagoshima University Museum (Korimoto); A.C. Bentley and the University of Kansas Biodiversity Institute (Lawrence). We thank G. Watkins-Colwell, Yale Peabody Museum (New Haven) for help with tissue collection processing. We thank C. Atta and the Bernice Pauahi Bishop Museum (Honolulu) for permission to use John E. Randall’s butterflyfish images and J. Johnson and the Life Science Studios for the butterflyfish illustrations.

## Author contributions

Conceptualization, K.A. and T.J.N.; methodology, K.A., T.J.N., C.D.B., M.E.A., P.C.W. and J.R.H.; validation, K.A., T.J.N., C.D.B., M.E.A., P.C.W. and J.R.H.; formal analysis, K.A.; investigation, K.A.; resources, J.R.H., P.C.W. and T.J.N.; writing original draft, K.A.; writing-review and editing, K.A., T.J.N., C.D.B., M.E.A., P.C.W. and J.R.H.; visualization, K.A.; supervision, K.A. and T.J.N.; project administration, T.J.N.; funding acquisition, P.C.W. and T.J.N.

---

**Extended Data Table 1 Tests of *Chaetodontinae* species richness against comparator diversification rates.** For each comparator reef-fish lineage (29 lineages plus *Heniochinae*), the 95% confidence interval on the species richness expected for a clade of the crown age of *Chaetodontinae* (7.78 Ma) diversifying at that lineage’s crown-group rate (method-of-moments estimator), under no extinction (ε=0) and high extinction (ε=0.9), and whether the observed richness of *Chaetodontinae* (110 species) exceeded the upper bound (Supplementary Table 4 gives the rates).

**Extended Data Table 2 MiSSE model comparison.** Six models with one to three hidden rate categories for turnover and/or the extinction fraction were fitted to 166 tips (single global sampling fraction, 0.73) and ranked by AICc. k, number of free parameters; ln L, log-likelihood; ΔAICc, relative to the best model; AICc weights are relative to the six models; AICc gain is the AICc of the constant-rate model minus that of each model; the last two columns summarise the marginal reconstruction of each model.

**Extended Data Table 3 Morphology–ecology associations in Chaetodontidae.** Two-block PLS and pGLS results for three trait sets (all seven traits; body traits; jaw traits) against a combined ecospace (depth and corallivory score) and, separately, against depth and corallivory alone (n=82 per test). PLS r, R² and permutation P show whether morphology and ecology covary more than expected by chance; pGLS R², P and λ re-test the relationship after accounting for shared ancestry.

## Extended Data legends

**Extended Data Fig. 1.**
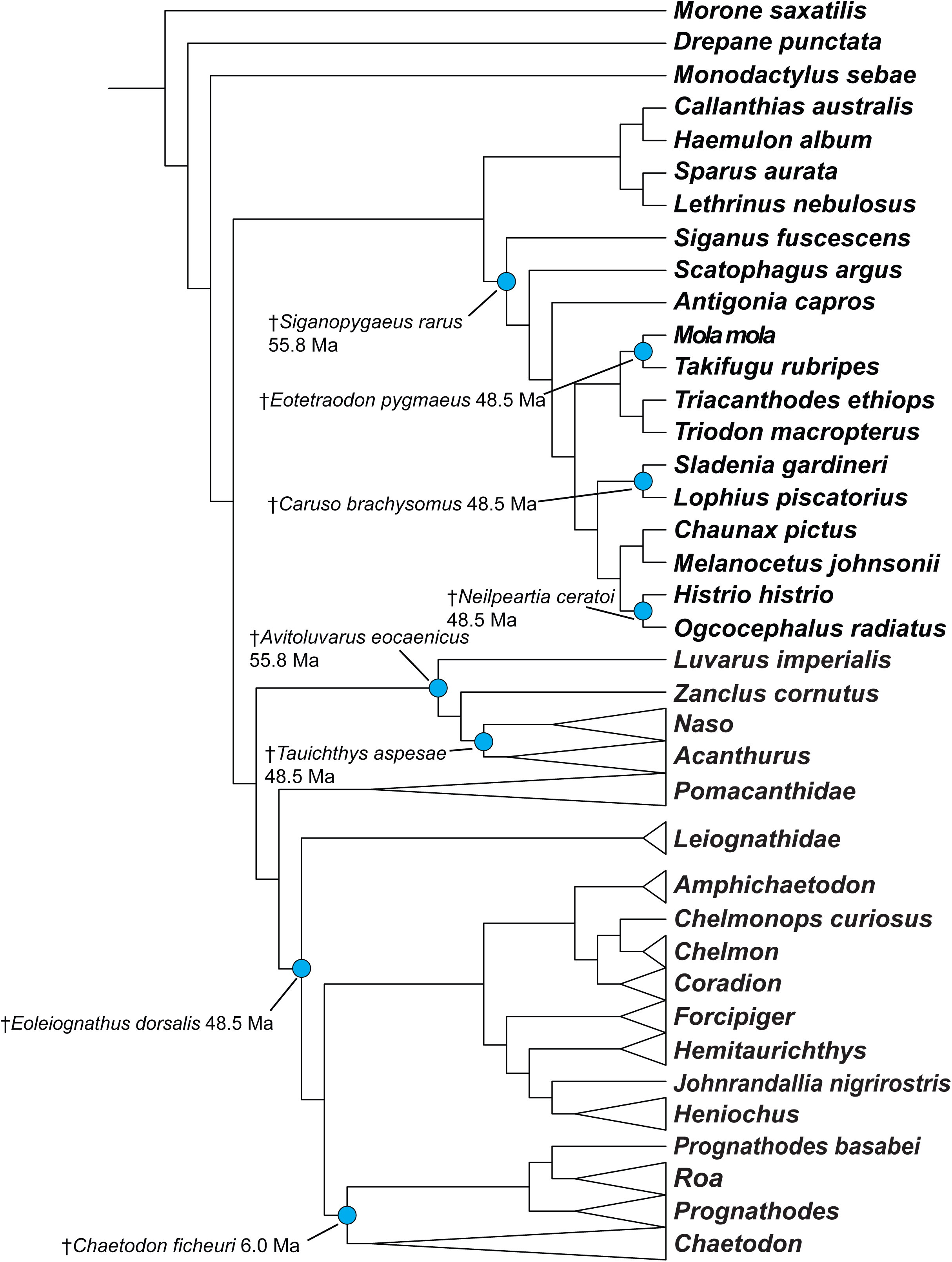
Fossil calibration scheme for the time-calibrated *Chaetodontidae* phylogeny. Phylogeny of *Chaetodontidae* and outgroups inferred from UCE loci in this study. Blue circles mark the phylogenetic placement of each fossil calibration point used in the BEAST fossilized birth-death relaxed molecular clock analyses. Fossil taxon names (†) and their estimated ages (Ma) are given beside each calibration point. Major ingroup and outgroup clades are collapsed for clarity.

**Extended Data Fig. 2.**
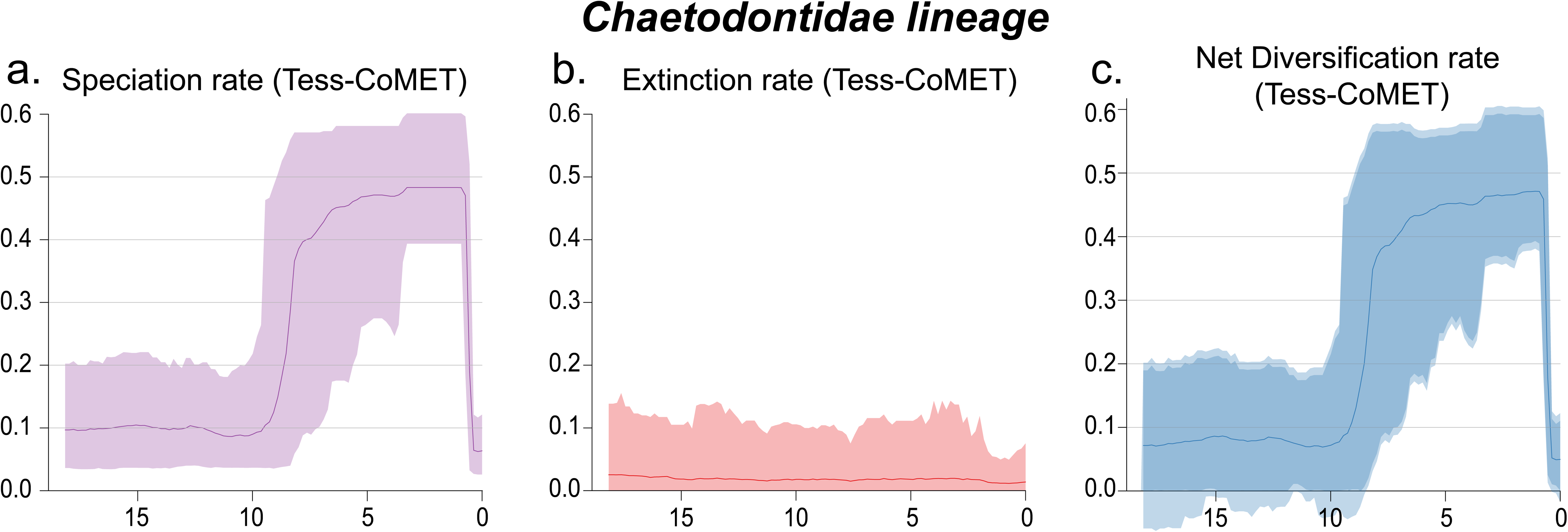
Temporal dynamics of speciation, extinction and net diversification in *Chaetodontidae*. Bayesian estimates from episodic birth–death models for *Chaetodontidae* (a–c): speciation rate (a), extinction rate (b), and net diversification rate (c). Lines show posterior medians, and shading shows the 95% credible intervals. Horizontal axes show time in millions of years before present.

**Extended Data Fig. 3.**
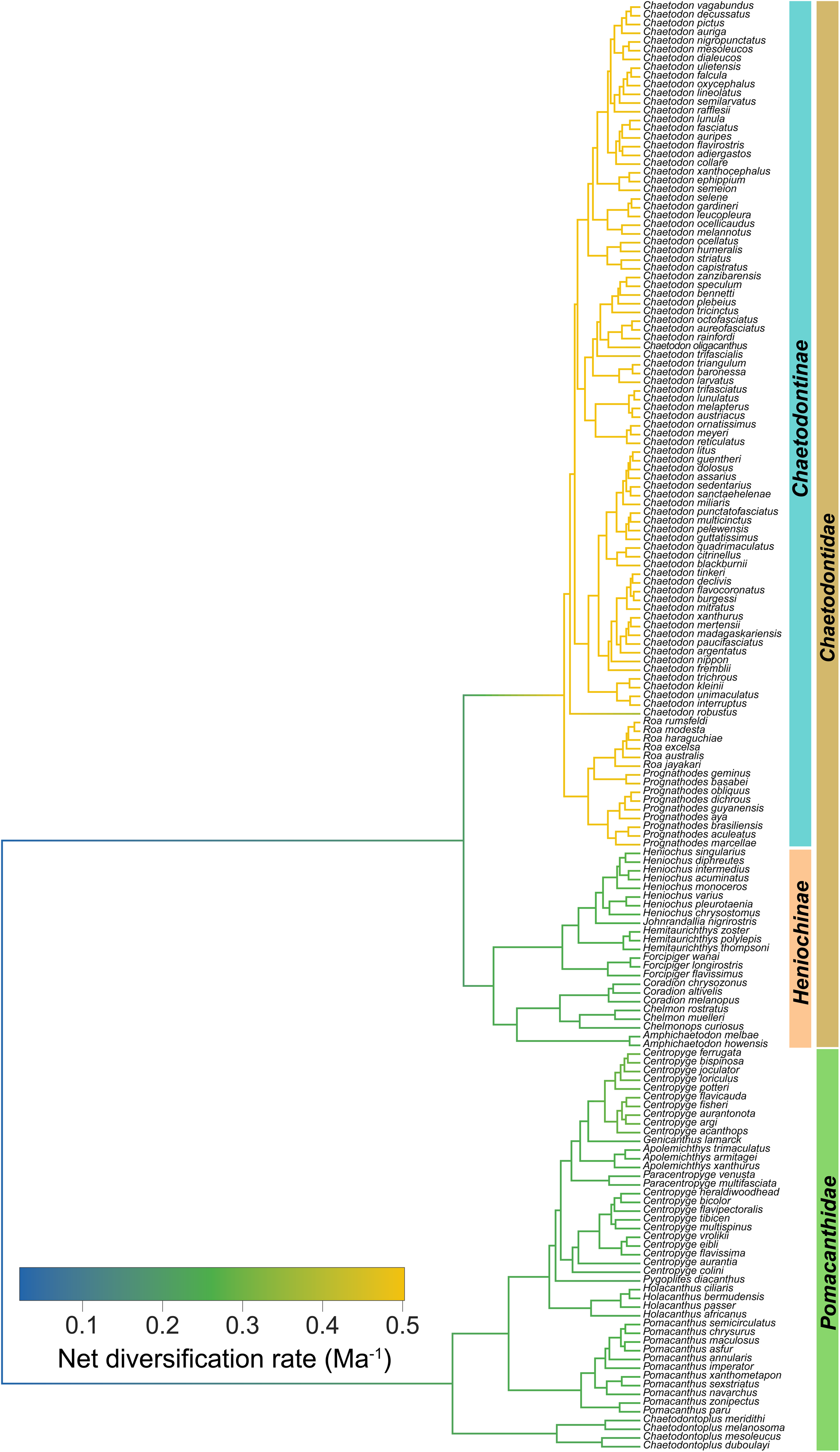
MiSSE reconstruction of net diversification rate across *Chaetodontidae* and *Pomacanthidae*. Time-calibrated phylogeny of 166 species: 120 *Chaetodontidae* (23 *Heniochinae* and 97 *Chaetodontinae*) and 46 species of the sister family *Pomacanthidae*, included so that rate shifts within *Chaetodontidae* can be detected relative to close relatives. Branches are coloured by net diversification rate (speciation minus extinction; Ma⁻¹) under the best-supported MiSSE model, a hidden-state model that uses no trait data; rates are averaged over hidden categories weighted by their marginal probabilities and interpolated along branches. Coloured bars mark *Chaetodontinae*, *Heniochinae*, *Pomacanthidae* and *Chaetodontidae*.

**Extended Data Fig. 4.**
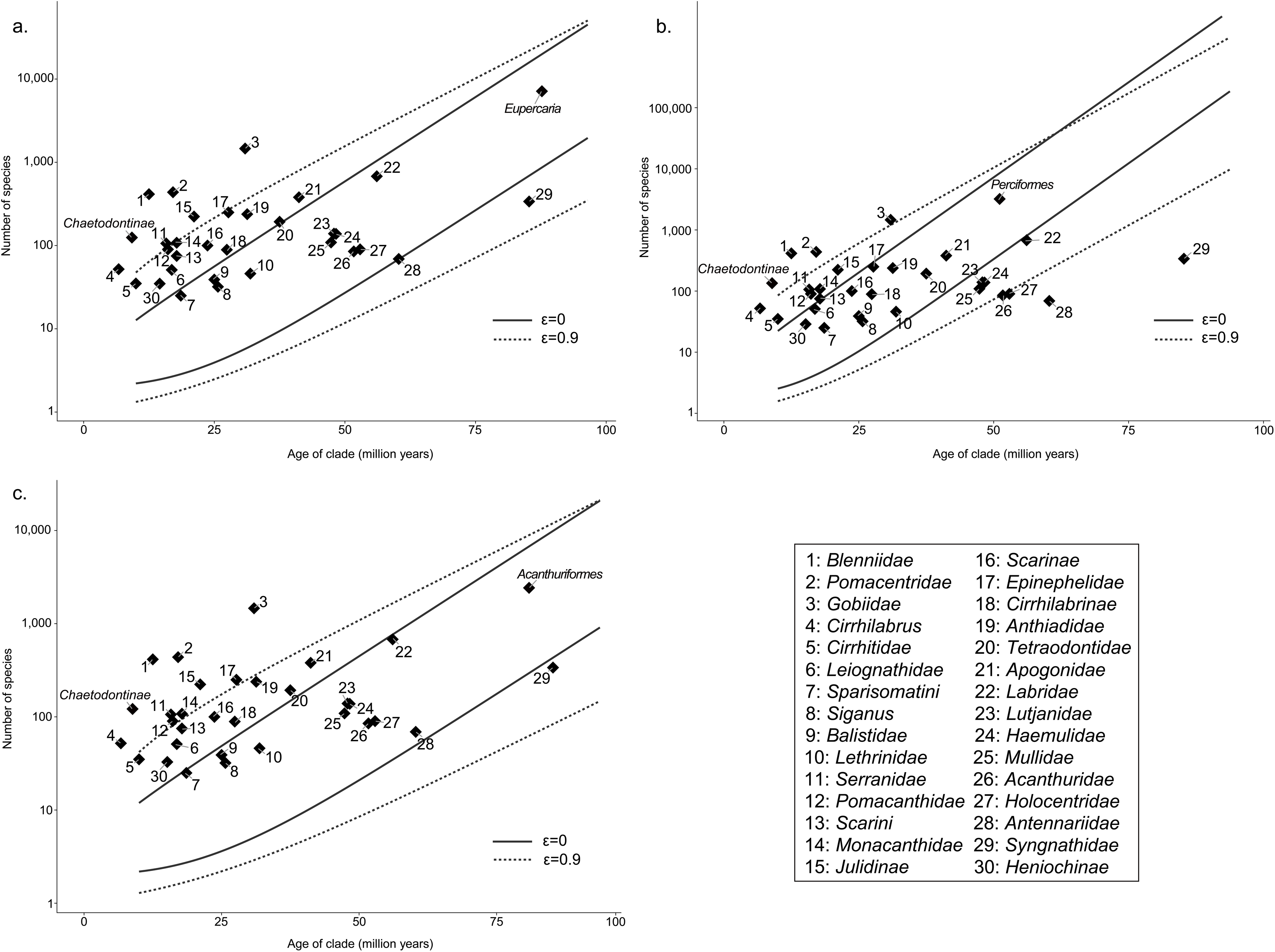
*Chaetodontinae* diversification relative to progressively broader background rates. Number of species (log scale) versus crown age for *Chaetodontinae* and the 29 comparator reef-fish lineages, plotted against the 95% confidence intervals for expected richness under the background diversification rate of *Eupercaria* (a), *Perciformes* (b) and *Acanthuriformes* (c) at ε=0 (solid) and ε=0.9 (dotted). Numbered points follow the key at right, shared across panels; points above a line exceed the richness expected for that background rate. The corresponding analysis for *Percomorpha* is Fig. 3b.

**Extended Data Fig. 5.**
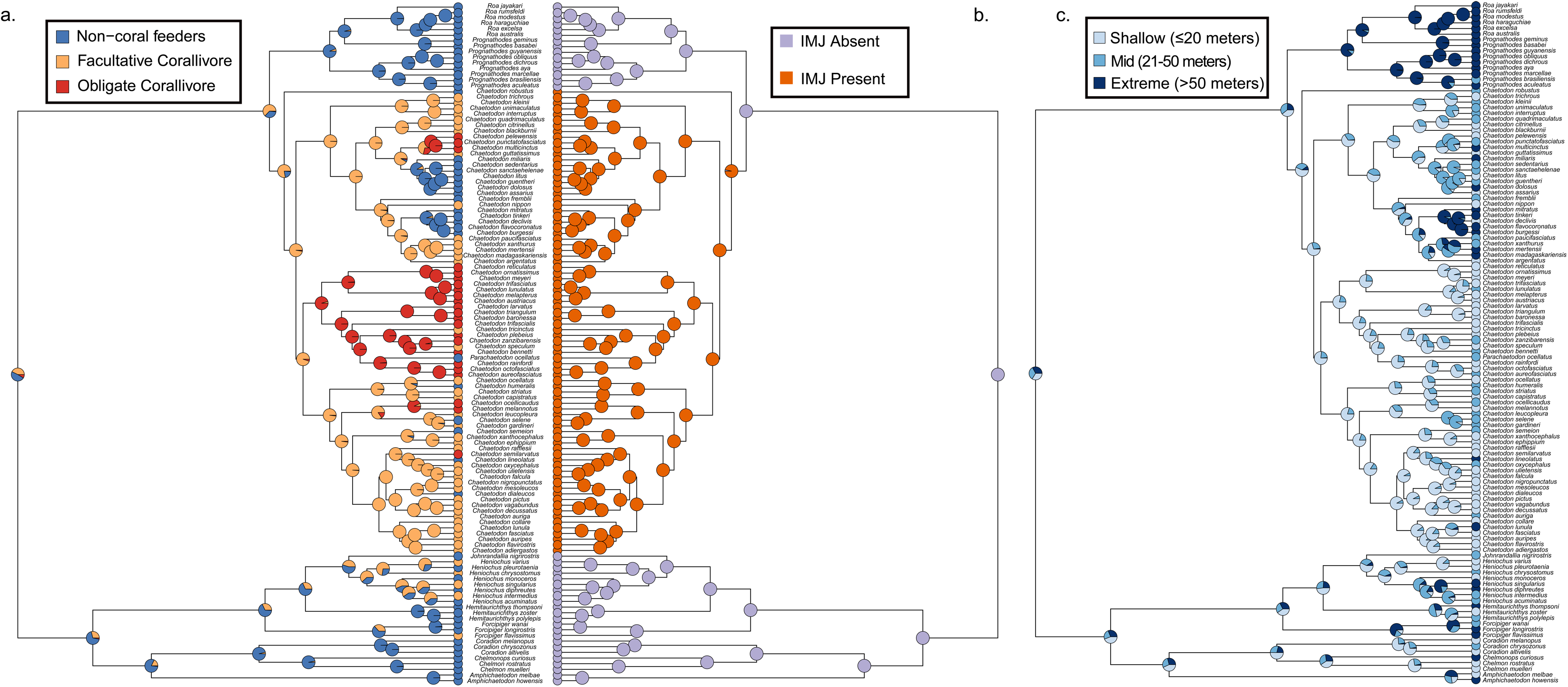
Ancestral state reconstruction of feeding strategy, jaw joint and depth category in *Chaetodontidae*. Maximum-likelihood ancestral states across the 120-species phylogeny for (a) feeding strategy (non-coral feeder, facultative corallivore, obligate corallivore), (b) presence or absence of the intramandibular joint (IMJ) and (c) depth category (shallow, ≤20 m; mid, 21–50 m; extreme, >50 m). Pie charts at nodes show the marginal probability of each ancestral state; tip colours show the observed state.

**Extended Data Fig. 6.**
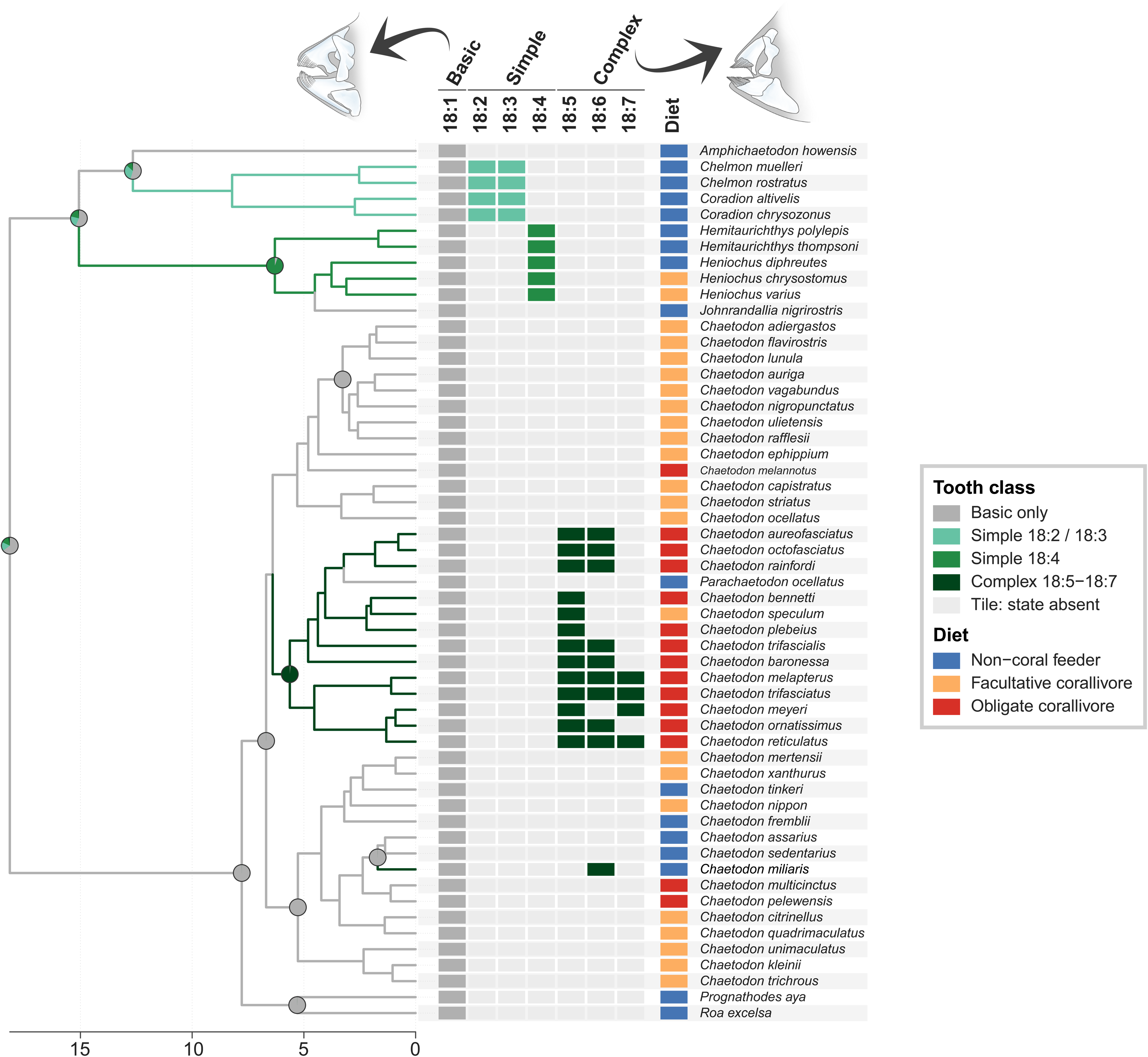
Ancestral states of tooth characters in *Chaetodontidae*. Time-calibrated phylogeny of the 55 species with tooth data. Tiles show the observed state of seven binary tooth characters (18:1–18:7; 18:1 is the basic state present in all species), coloured when present and light grey when absent; the Diet column gives each species’ feeding strategy (non-coral feeder, facultative corallivore or obligate corallivore). Species were assigned to four tooth classes (complex if 18:5–18:7 is present; otherwise simple 18:2/18:3; otherwise simple 18:4; otherwise basic only) and the class was reconstructed under an equal-rates model.

**Extended Data Fig. 7.**
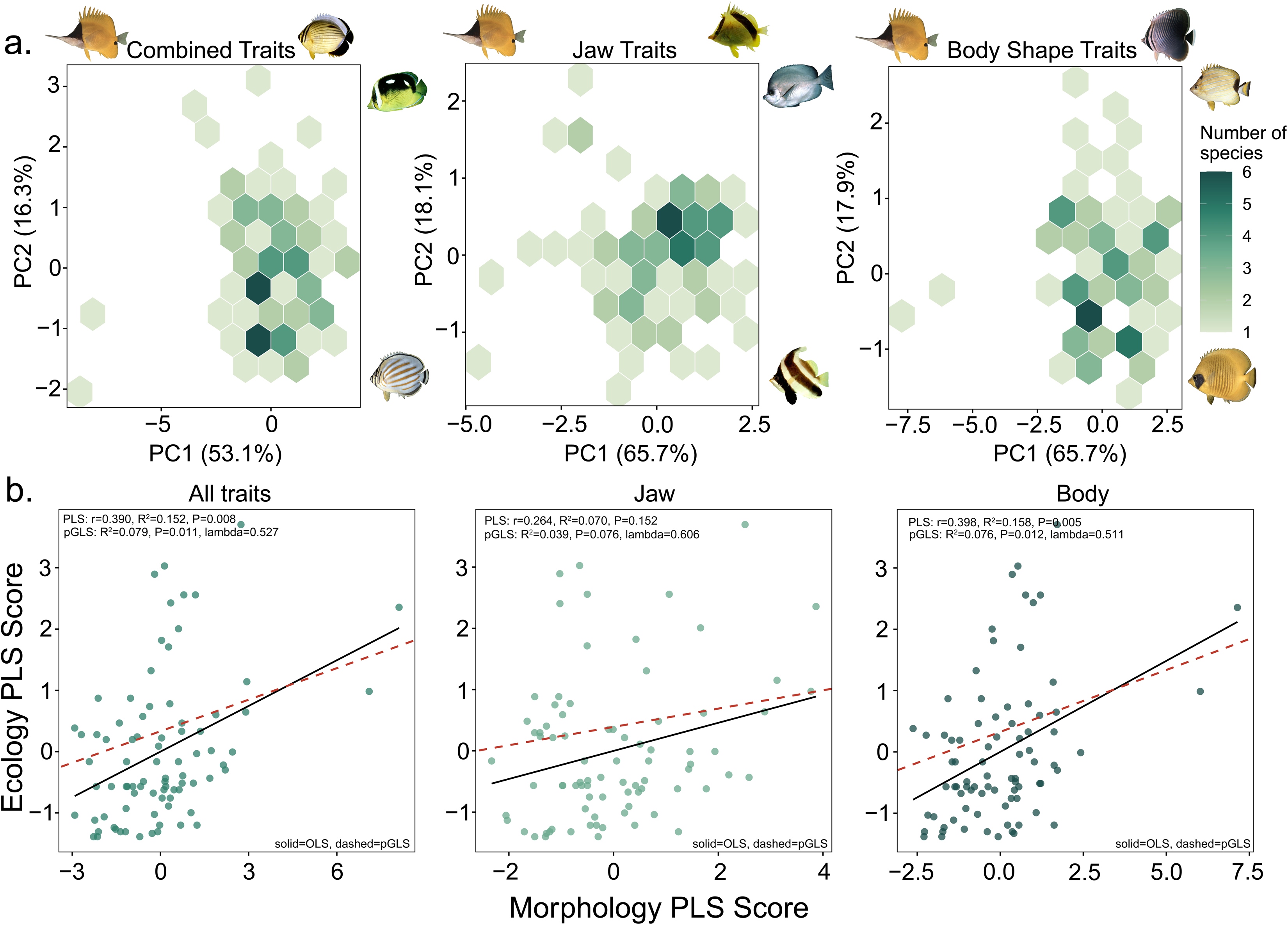
Morphospace occupation and morphology–ecology covariation in *Chaetodontidae*. (a) Density of species (n=82) in principal-component space of size-corrected traits: all seven traits combined, jaw traits (head depth, lower jaw length, mouth width) and body traits (maximum body depth, maximum fish width, minimum caudal peduncle depth and width); percentages are the variance explained by each axis. (b) Two-block partial least squares (PLS) and phylogenetic generalized least squares (pGLS) relating ecology scores (depth and corallivory) to morphology scores for the three trait sets (n=82): solid lines, ordinary least squares fit; dashed lines, pGLS fit. PLS r, R² and permutation P, and pGLS R², P and λ are given on each panel (Extended Data Table 3).

**Extended Data Table 1.** Results of the formal significance test comparing the diversification rate of *Chaetodontinae* against 29 comparator lineages and its sister lineage, *Heniochinae*, following the crown-age method-of-moments framework of Magallón and Sanderson (2001). For each comparator lineage and extinction-fraction scenario (ε=0, no extinction; ε=0.9, high extinction), we calculated the 95% confidence interval on the species richness expected for a clade of *Chaetodontinae’s* crown age (7.78 Ma) diversifying at that comparator’s estimated rate (see Supplementary Table 4). The observed richness of *Chaetodontinae* (110 species) was considered significantly elevated relative to any given clade when it exceeded the 95% upper bound of the estimated diversification rate of that clade.

| **Comparator lineage** | **Expected richness 95% CI (ε=0)** | **Significant** | **Expected richness 95% CI (ε=0.9)** | **Significant** |
|---|---|---|---|---|
| *Heniochinae* | 2.5 – 20.4 | Yes | 1.3 – 40.0 | Yes |
| *Scarinae* | 2.4 – 18.7 | Yes | 1.3 – 49.4 | Yes |
| *Pomacanthidae* | 3.0 – 33.6 | Yes | 1.5 – 80.1 | Yes |
| *Tetraodontidae* | 2.2 – 12.9 | Yes | 1.3 – 37.4 | Yes |
| *Monacanthidae* | 2.9 – 30.5 | Yes | 1.5 – 76.9 | Yes |
| *Pomacentridae* | 4.2 – 63.3 | Yes | 2.2 – 179.0 | **No** |
| *Balistidae* | 2.2 – 12.6 | Yes | 1.2 – 28.0 | Yes |
| *Gobiidae* | 2.8 – 27.9 | Yes | 1.7 – 100.3 | Yes |
| *Blenniidae* | 8.1 – 152.7 | **No** | 3.4 – 352.0 | **No** |
| *Mullidae* | 2.1 – 9.2 | Yes | 1.2 – 23.5 | Yes |
| *Syngnathidae* | 2.1 – 7.2 | Yes | 1.1 – 19.2 | Yes |
| *Leiognathidae* | 2.6 – 23.4 | Yes | 1.4 – 52.4 | Yes |
| *Julidinae* | 2.9 – 30.3 | Yes | 1.6 – 87.0 | Yes |
| *Apogonidae* | 2.2 – 13.5 | Yes | 1.3 – 42.5 | Yes |
| *Labridae* | 2.2 – 11.0 | Yes | 1.2 – 35.1 | Yes |
| *Scarini* | 2.7 – 25.8 | Yes | 1.4 – 62.0 | Yes |
| *Sparisomatini* | 2.3 – 14.6 | Yes | 1.2 – 29.1 | Yes |
| *Cirrhilabrinae* | 2.3 – 14.9 | Yes | 1.3 – 38.9 | Yes |
| *Cirrhilabrus* | 12.0 – 243.7 | **No** | 2.8 – 263.9 | **No** |
| *Anthiadidae* | 2.3 – 16.8 | Yes | 1.3 – 50.8 | Yes |
| *Epinephelidae* | 2.5 – 20.2 | Yes | 1.4 – 61.2 | Yes |
| *Serranidae* | 3.2 – 38.0 | Yes | 1.6 – 91.7 | Yes |
| *Acanthuridae* | 2.1 – 8.2 | Yes | 1.1 – 19.7 | Yes |
| *Holocentridae* | 2.1 – 8.1 | Yes | 1.1 – 19.7 | Yes |
| *Cirrhitidae* | 3.7 – 50.3 | Yes | 1.6 – 83.8 | Yes |
| *Lethrinidae* | 2.1 – 10.4 | Yes | 1.2 – 23.9 | Yes |
| *Lutjanidae* | 2.1 – 9.5 | Yes | 1.2 – 25.4 | Yes |
| *Haemulidae* | 2.1 – 9.5 | Yes | 1.2 – 25.1 | Yes |
| Siganus | 2.2 – 11.4 | Yes | 1.2 – 24.4 | Yes |
| Antennariidae | 2.1 – 7.1 | Yes | 1.1 – 16.0 | Yes |

**Extended Data Table 2.**
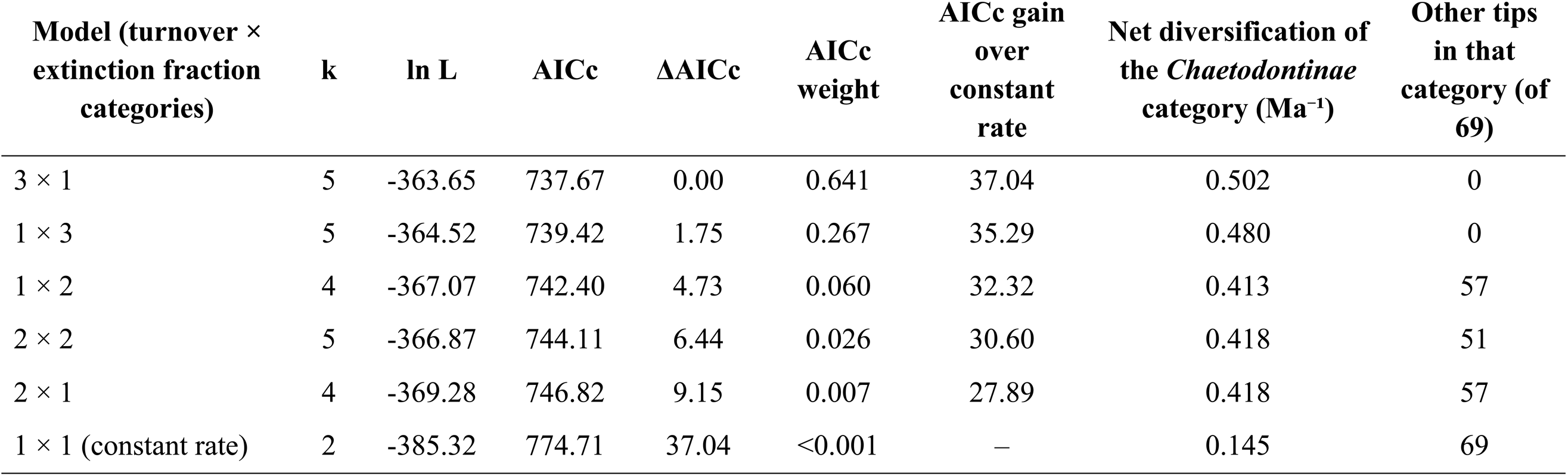
MiSSE model comparison for *Chaetodontidae* and its sister family *Pomacanthidae* (166 tips). Six models with one to three hidden rate categories for turnover and/or the extinction fraction were fitted with a single global sampling fraction (0.73) and ranked by AICc. Models are labelled by their number of turnover × extinction-fraction categories; k = number of free parameters; ln L = log-likelihood; ΔAICc is relative to the best model and AICc weights are relative to the six models; “AICc gain over constant rate” is the AICc of the constant-rate model minus the AICc of each model. The last two columns summarise the marginal reconstruction of each model.

**Extended Data Table 3.** Two-block PLS and phylogenetic generalized least squares (pGLS) results testing morphology–ecology associations for three trait sets: All traits (max body depth, max fish width, head depth, lower jaw length, mouth width, min caudal peduncle depth, and min caudal peduncle width); Body (max body depth, max fish width, min caudal peduncle depth, and min caudal peduncle width); and Jaw (head depth, lower jaw length, and mouth width). Using a combined ecospace (depth and corallivory score) and, separately, depth and corallivory tested as independent single-variable ecology blocks (n = 82 per test). PLS r, R², and P measure how strongly morphology and ecology co-vary and whether that covariation exceeds chance (permutation test), while pGLS R², P, and λ re-test that same relationship after correcting for phylogenetic non-independence among species, so it reflects the ecology signal that remains once shared ancestry is accounted for.

| **Trait set** | **Ecology** | **n** | **PLS r** | **PLS R<sup>2</sup>** | **PLS P** | **pGLS R<sup>2</sup>** | **pGLS P** | **pGLS λ** |
|---|---|---|---|---|---|---|---|---|
| All traits | Combined | 82 | 0.390 | 0.152 | 0.008 | 0.079 | 0.011 | 0.527 |
| Body | Combined | 82 | 0.398 | 0.158 | 0.005 | 0.076 | 0.012 | 0.511 |
| Jaw | Combined | 82 | 0.264 | 0.070 | 0.152 | 0.039 | 0.076 | 0.606 |
| All traits | Depth only | 82 | 0.398 | 0.158 | 0.010 | 0.130 | <0.001 | 0.161 |
| Body | Depth only | 82 | 0.402 | 0.161 | 0.002 | 0.130 | <0.001 | 0.139 |
| Jaw | Depth only | 82 | 0.275 | 0.076 | 0.047 | 0.067 | 0.019 | 0.243 |
| All traits | Corallivory only | 82 | 0.297 | 0.088 | 0.075 | 0.009 | 0.406 | 1.000 |
| Body | Corallivory only | 82 | 0.292 | 0.085 | 0.032 | 0.001 | 0.786 | 1.000 |
| Jaw | Corallivory only | 82 | 0.212 | 0.045 | 0.203 | 0.016 | 0.259 | 1.000 |

## References

1. McGreevy, T. J., Crawford, N. G., Legreneur, P. & Schneider, C. J. Influence of geographic isolation and the environment on gene flow among phenotypically diverse lizards. Heredity 133, 317–330 (2024).

2. Grant, P. R. & Grant, B. R. Evolution of Character Displacement in Darwin’s Finches. Science 313, 224–226 (2006).

3. Reno, P. L. Genetic and developmental basis for parallel evolution and its significance for hominoid evolution. Evolutionary Anthropology: Issues, News, and Reviews 23, 188– 200 (2014).

4. Brawand, D. et al. The genomic substrate for adaptive radiation in African cichlid fish. Nature 513, 375–381 (2014).

5. Stroud, J. T. & Losos, J. B. Ecological Opportunity and Adaptive Radiation. Annual Review of Ecology, Evolution, and Systematics 47, 507–532 (2016).

6. Gillespie, R. G. et al. Comparing Adaptive Radiations Across Space, Time, and Taxa. J Hered 111, 1–20 (2020).

7. DiBattista, J. D. et al. Ice ages and butterflyfishes: Phylogenomics elucidates the ecological and evolutionary history of reef fishes in an endemism hotspot. Ecology and Evolution 8, 10989–11008 (2018).

8. Siqueira, A. C., Kiessling, W. & Bellwood, D. R. Fast-growing species shape the evolution of reef corals. Nature Communications 13, 2426 (2022).

9. Siqueira, A. C., Morais, R. A., Bellwood, D. R. & Cowman, P. F. Trophic innovations fuel reef fish diversification. Nat Commun 11, 2669 (2020).

10. Tea, Y.-K. et al. Phylogenomic Analysis of Concatenated Ultraconserved Elements Reveals the Recent Evolutionary Radiation of the Fairy Wrasses (Teleostei: Labridae: Cirrhilabrus). Syst Biol 71, 1–12 (2022).

11. Bellwood, D. R. et al. Evolutionary history of the butterflyfishes (f: Chaetodontidae) and the rise of coral feeding fishes. J. Evol. Biol. 23, 335–349 (2010).

12. Price, S. A., Holzman, R., Near, T. J. & Wainwright, P. C. Coral reefs promote the evolution of morphological diversity and ecological novelty in labrid fishes. Ecology Letters 14, 462–469 (2011).

13. Hundt, P. J., Iglésias, S. P., Hoey, A. S. & Simons, A. M. A multilocus molecular phylogeny of combtooth blennies (Percomorpha: Blennioidei: Blenniidae): Multiple invasions of intertidal habitats. Molecular Phylogenetics and Evolution 70, 47–56 (2014).

14. Simpson, G. G. The Major Features of Evolution. (New York, Columbia University Press, 1953).

15. Schluter, D. The Ecology of Adaptive Radiation. (Oxford University Press, Oxford, New York, 2000).

16. Price, S. A., Claverie, T., Near, T. J. & Wainwright, P. C. Phylogenetic insights into the history and diversification of fishes on reefs. Coral Reefs 34, 997–1009 (2015).

17. Burress, E. D. & Wainwright, P. C. Adaptive radiation in labrid fishes: A central role for functional innovations during 65 My of relentless diversification. Evol 73, 346–359 (2019).

18. Cowman, P. F., Bellwood, D. R. & van Herwerden, L. Dating the evolutionary origins of wrasse lineages (Labridae) and the rise of trophic novelty on coral reefs. Molecular Phylogenetics and Evolution 52, 621–631 (2009).

19. Tian, S. Y. et al. Cenozoic history of the tropical marine biodiversity hotspot. Nature 632, 343–349 (2024).

20. Brownstein, C. D. et al. Phylogenomics establishes an Early Miocene reconstruction of reef vertebrate diversity. Science Advances 11, eadu6149 (2025).

21. Siqueira, A. C., Kiessling, W., Raja, N. B. & Bellwood, D. R. The rise and fall of the world’s greatest marine biodiversity hotspot. Science Advances 12, eaec7264 (2026).

22. Siqueira, A. C., Muruga, P. & Bellwood, D. R. On the evolution of fish–coral interactions. Ecology Letters 26, 1348–1358 (2023).

23. Fessler, J. L. & Westneat, M. W. Molecular phylogenetics of the butterflyfishes (Chaetodontidae): Taxonomy and biogeography of a global coral reef fish family. Molecular Phylogenetics and Evolution 45, 50–68 (2007).

24. Hemingson, C. R., Cowman, P. F., Hodge, J. R. & Bellwood, D. R. Colour pattern divergence in reef fish species is rapid and driven by both range overlap and symmetry. Ecology Letters 22, 190–199 (2019).

25. Kelley, J. L., Fitzpatrick, J. L. & Merilaita, S. Spots and stripes: ecology and colour pattern evolution in butterflyfishes. Proc Biol Sci 280, 20122730 (2013).

26. Pratchett, M. S., Berumen, M. L. & Kapoor, B. G. Biology of Butterflyfishes. Biology of Butterflyfishes 352 (CRC Press, 2013).

27. Tea, Y.-K., Blum, S. D., Rocha, L. A., Parkinson, K. & Stanley, E. L. A New Deepwater Species of Butterflyfish (Teleostei: Chaetodontidae: Prognathodes) from Mozambique, Africa, with a Systematic Review of Chaetodontid Relationships on the Basis of Osteology. Ichthyology & Herpetology 114, 474–494 (2026).

28. Cole, A. J., Pratchett, M. S. & Jones, G. P. Diversity and functional importance of coral-feeding fishes on tropical coral reefs. Fish and Fisheries 9, 286–307 (2008).

29. Konow, N., Bellwood, D. R., Wainwright, P. C. & Kerr, A. M. Evolution of novel jaw joints promote trophic diversity in coral reef fishes. Biol J Linn Soc 93, 545–555 (2008).

30. Konow, N., Price, S., Abom, R., Bellwood, D. & Wainwright, P. Decoupled diversification dynamics of feeding morphology following a major functional innovation in marine butterflyfishes. Proc. R. Soc. B 284, 20170906 (2017).

31. Motta, P. J. Functional morphology of the feeding apparatus of ten species of Pacific butterflyfishes (Perciformes, Chaetodontidae): an ecomorphological approach. Environ Biol Fish 22, 39–67 (1988).

32. Motta, P. J. Functional morphology of the head of Hawaiian and Mid-Pacific butterflyfishes (Perciformes, Chaetodontidae). Environ Biol Fish 13, 253–276 (1985).

33. Motta, P. J. Dentition patterns among Pacific and Western Atlantic butterflyfishes (Perciformes, Chaetodontidae): relationship to feeding ecology and evolutionary history. Environ Biol Fish 25, 159–170 (1989).

34. Berumen, M. L., Pratchett, M. S. & Goodman, B. A. Relative gut lengths of coral reef butterflyfishes (Pisces: Chaetodontidae). Coral Reefs 30, 1005–1010 (2011).

35. Motta, P. J. Functional morphology of the head of the inertial suction feeding butterflyfish, Chaetodon miliaris (Perciformes, Chaetodontidae). Journal of Morphology 174, 283–312 (1982).

36. Copus, J. M. & Gibb, A. C. A forceful upper jaw facilitates picking-based prey capture: biomechanics of feeding in a butterflyfish, Chaetodon trichrous. Zoology 116, 336–347 (2013).

37. Baldwin, B. G. & Sanderson, M. J. Age and rate of diversification of the Hawaiian silversword alliance (Compositae). Proceedings of the National Academy of Sciences 95, 9402–9406 (1998).

38. Near, T. J. et al. Phylogenetic analysis of Antarctic notothenioids illuminates the utility of RADseq for resolving Cenozoic adaptive radiations. Molecular Phylogenetics and Evolution 129, 268–279 (2018).

39. Rabosky, D. L. et al. An inverse latitudinal gradient in speciation rate for marine fishes. Nature 559, 392–395 (2018).

40. Near, T. J. & Thacker, C. E. Phylogenetic Classification of Living and Fossil Ray-Finned Fishes (Actinopterygii). Bull Peabody Museum Natural History 65, 3–302 (2024).

41. Bouchon-Navaro, Y. Partitioning of food and space resources by chaetodontid fishes on coral reefs. Journal of Experimental Marine Biology and Ecology 103, 21–40 (1986).

42. Harmelin-Vivien, M. L. & Bouchon-Navaro, Y. Feeding diets and significance of coral feeding among Chaetodontid fishes in Moorea (French Polynesia). Coral Reefs 2, 119– 127 (1983).

43. Harmelin-Vivien, M. L. Implications of feeding specialization on the recruitment processes and community structure of butterflyfishes. Environ Biol Fish 25, 101–110 (1989).

44. Sano, M. Feeding habits of Japanese butterflyfishes (Chaetodontidae). Environ Biol Fish 25, 195–203 (1989).

45. Randall, J. E. Food habits of reef fishes of the West Indies. Studies in Tropical Oceanography 5, 665–847 (1967).

46. Blum, S. D. The Osteology and Phylogeny of the Chaetodontidae (Teleostei: Perciformes). (University of Hawaii, Honolulu, HI, 1988). doi:10.5281/zenodo.3340185.

47. Givnish, T. J. Adaptive radiation versus ‘radiation’ and ‘explosive diversification’: why conceptual distinctions are fundamental to understanding evolution. New Phytologist 207, 297–303 (2015).

48. Olson, M. E. & Arroyo-Santos, A. Thinking in continua: beyond the “adaptive radiation” metaphor. BioEssays 31, 1337–1346 (2009).

49. Gavrilets, S. & Losos, J. B. Adaptive Radiation: Contrasting Theory with Data. Science 323, 732–737 (2009).

50. Kiessling, W., Simpson, C. & Foote, M. Reefs as Cradles of Evolution and Sources of Biodiversity in the Phanerozoic. Science 327, 196–198 (2010).

51. McCaffrey, J. C., Wallace, M. W. & Gallagher, S. J. A Cenozoic Great Barrier Reef on Australia’s North West shelf. Global and Planetary Change 184, 103048 (2020).

52. Mathew, M. et al. The emergence of Miocene reefs in South China Sea and its resilient adaptability under varying eustatic, climatic and oceanographic conditions. Sci Rep 10, 7141 (2020).

53. Renema, W. et al. Are coral reefs victims of their own past success? Science Advances 2, e1500850 (2016).

54. Marrable, I., Peoples, N., Kolmann, M. A., Huie, J. M. & Figueira, W. F. The Evolutionary History and Morphological Divergence of Corallivorous Fishes on Coral Reefs. Integr Org Biol 8, obag032 (2026).

55. Tamir, R., Eyal, G., Kramer, N., Laverick, J. H. & Loya, Y. Light environment drives the shallow-to-mesophotic coral community transition. Ecosphere 10, e02839 (2019).

56. Muir, P., Wallace, C., Bridge, T. C. L. & Bongaerts, P. Diverse Staghorn Coral Fauna on the Mesophotic Reefs of North-East Australia. PLOS ONE 10, e0117933 (2015).

57. Miller, A. H., Stroud, J. T. & Losos, J. B. The ecology and evolution of key innovations. Trends in Ecology & Evolution 38, 122–131 (2023).

58. Hodge, J. R. et al. Constraints on the Ecomorphological Convergence of Zooplanktivorous Butterflyfishes. Integr Org Biol 3, obab014 (2021).

59. Clark, T. R. et al. U-Th dating reveals regional-scale decline of branching Acropora corals on the Great Barrier Reef over the past century. Proceedings of the National Academy of Sciences 114, 10350–10355 (2017).

60. Tkachenko, K. S., Wu, B.-J., Fang, L.-S. & Fan, T.-Y. Dynamics of a coral reef community after mass mortality of branching Acropora corals and an outbreak of anemones. Mar Biol 151, 185–194 (2007).

61. Anderson, K. D., Heron, S. F. & Pratchett, M. S. Species-specific declines in the linear extension of branching corals at a subtropical reef, Lord Howe Island. Coral Reefs 34, 479–490 (2015).

62. Queiroz, K. de & Cantino, P. International Code of Phylogenetic Nomenclature (PhyloCode). (CRC Press, Boca Raton, 2020). doi:10.1201/9780429446320.

63. Faircloth, B. C. PHYLUCE is a software package for the analysis of conserved genomic loci. Bioinformatics 32, 786–788 (2016).

64. Alfaro, M. E. et al. Explosive diversification of marine fishes at the Cretaceous– Palaeogene boundary. Nat Ecol Evol 2, 688–696 (2018).

65. Near, T. J. et al. Phylogeny and tempo of diversification in the superradiation of spiny-rayed fishes. Proceedings of the National Academy of Sciences 110, 12738–12743 (2013).

66. Near, T. J. et al. Resolution of ray-finned fish phylogeny and timing of diversification. Proceedings of the National Academy of Sciences 109, 13698–13703 (2012).

67. Betancur-R. R., et al. The Tree of Life and a New Classification of Bony Fishes. PLoS Curr 5, ecurrents.tol.53ba26640df0ccaee75bb165c8c26288 (2013).

68. Hughes, L. C. et al. Comprehensive phylogeny of ray-finned fishes (Actinopterygii) based on transcriptomic and genomic data. Proceedings of the National Academy of Sciences 115, 6249–6254 (2018).

69. Ghezelayagh, A. et al. Prolonged morphological expansion of spiny-rayed fishes following the end-Cretaceous. Nat Ecol Evol 6, 1211–1220 (2022).

70. Nguyen, L.-T., Schmidt, H. A., von Haeseler, A. & Minh, B. Q. IQ-TREE: A Fast and Effective Stochastic Algorithm for Estimating Maximum-Likelihood Phylogenies. Mol Biol Evol 32, 268–274 (2015).

71. Bouckaert, R. et al. BEAST 2: A Software Platform for Bayesian Evolutionary Analysis. PLOS Computational Biology 10, e1003537 (2014).

72. Bouckaert, R. et al. BEAST 2.5: An advanced software platform for Bayesian evolutionary analysis. PLOS Computational Biology 15, e1006650 (2019).

73. Gavryushkina, A. et al. Bayesian Total-Evidence Dating Reveals the Recent Crown Radiation of Penguins. Syst Biol 66, 57–73 (2017).

74. Capobianco, A., Darlim, G. & Höhna, S. How to date a molecular phylogeny: comparison of effective priors between node calibration and fossilized birth–death. Proc Biol Sci 293, 20253255 (2026).

75. Yang, Z. & Rannala, B. Bayesian Estimation of Species Divergence Times Under a Molecular Clock Using Multiple Fossil Calibrations with Soft Bounds. Mol Biol Evol 23, 212–226 (2006).

76. Rambaut, A., Drummond, A. J., Xie, D., Baele, G. & Suchard, M. A. Posterior Summarization in Bayesian Phylogenetics Using Tracer 1.7. Syst Biol 67, 901–904 (2018).

77. Magallon, S. & Sanderson, M. J. Absolute diversification rates in angiosperm clades. Evol 55, 1762–1780 (2001).

78. Fricke, R., Eschmeyer, W. N. & Van der Laan, R. Eschmeyer’s catalog of fishes: genera, species, references. http://researcharchive.calacademy.org/research/ichthyology/catalog/fishcatmain.asp (2026).

79. Pennell, M. W. et al. geiger v2.0: an expanded suite of methods for fitting macroevolutionary models to phylogenetic trees. Bioinformatics 30, 2216–2218 (2014).

80. Harmon, L. J., Weir, J. T., Brock, C. D., Glor, R. E. & Challenger, W. GEIGER: investigating evolutionary radiations. Bioinformatics 24, 129–131 (2008).

81. Höhna, S., May, M. R. & Moore, B. R. TESS: an R package for efficiently simulating phylogenetic trees and performing Bayesian inference of lineage diversification rates. Bioinformatics 32, 789–791 (2016).

82. Sibert, E. C., Zill, M. E., Frigyik, E. T. & Norris, R. D. No state change in pelagic fish production and biodiversity during the Eocene–Oligocene transition. Nature Geoscience 13, 238–242 (2020).

83. Sibert, E. C., Hull, P. M. & Norris, R. D. Resilience of Pacific pelagic fish across the Cretaceous/Palaeogene mass extinction. Nature Geosci 7, 667–670 (2014).

84. Sibert, E. C. & Norris, R. D. New Age of Fishes initiated by the Cretaceous−Paleogene mass extinction. Proceedings of the National Academy of Sciences 112, 8537–8542 (2015).

85. Vasconcelos, T., O’Meara, B. C. & Beaulieu, J. M. A flexible method for estimating tip diversification rates across a range of speciation and extinction scenarios. Evolution 76, 1420–1433 (2022).

86. Pratchett, M. S. Dietary overlap among coral-feeding butterflyfishes (Chaetodontidae) at Lizard Island, northern Great Barrier Reef. Marine Biology 148, 373–382 (2005).

87. Friedman, M. The Use of Ranks to Avoid the Assumption of Normality Implicit in the Analysis of Variance. Journal of the American Statistical Association 32, 675–701 (1937).

88. Kendall, M. G. & Smith, B. B. The Problem of m Rankings. The Annals of Mathematical Statistics 10, 275–287 (1939).

89. Wilcoxon, F. Individual Comparisons by Ranking Methods. Biometrics Bulletin 1, 80–83 (1945).

90. Holm, S. A Simple Sequentially Rejective Multiple Test Procedure. Scandinavian Journal of Statistics 6, 65–70 (1979).

91. Nagelkerken, I., Van Der Velde, G., Wartenbergh, S. L. J., Nugues, M. M. & Pratchett, M. S. Cryptic dietary components reduce dietary overlap among sympatric butterflyfishes (Chaetodontidae). Journal of Fish Biology 75, 1123–1143 (2009).

92. Pratchett, M. S. Dietary selection by coral-feeding butterflyfishes (Chaetodontidae) on the Great Barrier Reef, Australia. (2007).

93. Froese, R. & Pauly, D. FishBase. https://www.fishbase.org/.

94. Revell, L. J. phytools: an R package for phylogenetic comparative biology (and other things). Methods in Ecology and Evolution 3, 217–223 (2012).

95. Paradis, E. & Schliep, K. ape 5.0: an environment for modern phylogenetics and evolutionary analyses in R. Bioinformatics 35, 526–528 (2019).

96. Huelsenbeck, J. P., Nielsen, R. & Bollback, J. P. Stochastic Mapping of Morphological Characters. Syst Biol 52, 131–158 (2003).

97. Bollback, J. P. SIMMAP: Stochastic character mapping of discrete traits on phylogenies. BMC Bioinformatics 7, 88 (2006).

98. Pagel, M. Inferring the historical patterns of biological evolution. Nature 401, 877–884 (1999).

99. Ives, A. R. & Garland, T., Jr. Phylogenetic Logistic Regression for Binary Dependent Variables. Syst Biol 59, 9–26 (2010).

100. Ronco, F. et al. Drivers and dynamics of a massive adaptive radiation in cichlid fishes. Nature 589, 76–81 (2021).

101. Rohlf, F. J. & Corti, M. Use of Two-Block Partial Least-Squares to Study Covariation in Shape. Syst Biol 49, 740–753 (2000).

102. Orme, D. et al. caper: Comparative Analyses of Phylogenetics and Evolution in R. 5, 1–36 (2013).

103. Price, S. A. et al. FishShapes v1: Functionally relevant measurements of teleost shape and size on three dimensions. Ecology 103, e3829 (2022).

