## Supplementary Information for "A coral-linked adaptive radiation in butterflyfishes"

**Table of Contents.**

Fossil Calibration Justification List ………. Pg. 2
Supplementary Results and Discussion …... Pg. 5
References…………………………………..Pg. 15
Supplementary Figure Captions…………….Pg. 22

**Fossil Calibration Justification List.**

†*Neilpeartia ceratoi*

**Node calibrated:** †*Neilpeartia ceratoi*[1] calibrates the node representing the MRCA of the clade containing *Histrio* and *Ogcocephalus* in our analysis. †*Eophryne barbutii* may be an equivalent fossil calibration[2], but †*Neilpeartia ceratoi* represents the most likely member of this crown clade among the Monte Bolca forms[1]. †*Neilpeartia ceratoi* is identified as an antennariid based on: ectopterygoid triradiate, endopterygoid present, epural present, spatulate post-maxillary process of the premaxilla, reduced opercle, dorsoventrally deep body shape, and enlarged third dorsal fin spine. [S1] Phylogenetic analysis of morphological characters placed †*Neilpeartia ceratoi* sister to the genus *Fowlerichthys*[1].

**Stratigraphy**: Monte Bolca, near Verona, Italy: early-late Ypresian, Eocene, Paleogene 48.96 to 48.5 Ma[3].

**Fossil tip age**: 48.5 Ma. The node calibration was created such that 97.5% of the probability distribution fell before this date.

†*Caruso brachysomus*

**Node calibrated:** †*Caruso brachysomus* calibrates the node representing the MRCA of the clade containing *Lophius* and *Sladenia* (=crown *Lophiidae*)[4]. †*Eosladenia caucasica* from the Middle Eocene of the northern Caucasus provides a penecontemporaneous fossil calibration[4,5]. The assignment of †*Caruso brachysomus* to total clade Lophiidae is based on: absence of a mesethmoid, an autogenous ascending process of the premaxilla, fusion of the ectopterygoid and endopterygoid, prominent anterodorsal process of the subopercle articulates with anteroventral margin of opercle via connective tissue, restriction of the dentition on the fifth ceratobranchial to discrete rows on lateral and medial margins, presence of a cleithral spine, skin unornamented, and eight caudal fin rays. †*Caruso brachysomus* is identified as a crown lophiid based on the presence of a strongly bifurcated opercle, and as sister to *Sladenia* by a narrow interorbital skull roof and a posteriorly directed expanded distal end of the posteriormost dorsal fin pterygiophore. These character combinations are based on a comprehensive phylogenetic analysis of lophiids using morphological data[4].

**Stratigraphy**: Monte Bolca, near Verona, Italy: early-late Ypresian, Eocene, Paleogene 48.96 to 48.5 Ma[3].

**Fossil tip age**: 48.5 Ma. The node calibration was created such that 97.5% of the probability distribution fell before this date.

†*Eotetraodon pygmaeus*

**Node calibrated:** †*Eotetraodon pygmaeus* calibrates the node representing the MRCA of the clade containing *Mola* and *Takifugu*. The placement of †*Eotetraodon pygmaeus* follows the phylogenetic analysis conducted in previous studies[6] and comparative anatomical assignments in Bannikov and Tyler[7], who principally identified this species as a pan-tetraodontid based on the division of the jaws into right and left halves.

**Stratigraphy**: Monte Bolca, near Verona, Italy: early-late Ypresian, Eocene, Paleogene 48.96 to 48.5 Ma[3].

**Fossil tip age**: 48.5 Ma. The node calibration was created such that 97.5% of the probability distribution fell before this date.

†*Siganopygaeus rarus*

**Node calibrated:** †*Siganopygaeus rarus* calibrates the node representing the MRCA of *Scatophagus argus* (*Scatophagidae*) and *Siganus spinus* (*Siganidae*) in our analysis. The justification for this fossil calibration as a member of Pan-*Siganidae* follows previous studies[8,9] and the phylogenetic analysis presented in a study[10].

**Stratigraphy**: Danatinsk Suite, Uylya-Kushlyuk locality, Turkmenistan: Thanetian-Ypresian, Paleocene-Eocene, 55.8 Ma[8,11].

**Fossil tip age**: 55.8 Ma. The node calibration was created such that 97.5% of the probability distribution fell before this date.

†*Avitoluvarus eocaenicus*

**Node calibrated:** †*Avitoluvarus eocaenicus* is a member of Pan-*Luvarus*[12] and calibrates the node representing the MRCA of *Luvarus imperialis* and *Zanclus cornutus* in our analysis. The placement of †*A. eocaenicus* in Pan-Luvarus as a member of †*Avitoluvarus* is supported by the following combination of characters: dentition reduced or lost in adults; median pterygial truss around most of body; pterygial truss shallow and not extensively interdigitated; two or fewer spines in dorsal fin; anal spines absent; all soft dorsal and anal fin rays unsegmented; distal end of first anal fin pterygiophore anteriorly elongated; anteriorly displaced anus; fusion of first four hypurals; broad overlap between caudal fin rays and hypural plate; pelvic fin rudimentary with increasing body size; proximal shafts of anal fin pterygiophores situation in first two interhaemal spaces[12,13].

**Stratigraphy**: Uylya-Kushlyuk locality, 2 km northeast of Uylya-Kushlyuk village, Turkmenistan; Danata Formation, uppermost Thanetian-lowermost Ypresian (Paleocene to Eocene)[8,12,14]. Following Ghezelayagh et al.[8], we use an age of 55.8 Ma.

**Fossil tip age**: 55.8 Ma.

†*Tauichthys aspesae*

**Node calibrated:** †*Tauichthys aspesae* is a member of Pan-*Acanthurinae*[15,16] and calibrates the node representing the MRCA of *Ctenochaetus striatus* and *Naso annulatus* in our analysis. The placement of †*T. aspesae* in *Acanthuridae* is supported by the following combination of characters: hypurals unconsolidated; three epurals; first dorsal and anal fin spines elongated such that they protrude prominently outward; absence of laterally expanded pterygial shields around first dorsal and anal fin spine bases; five branchiostegal rays[16].

**Stratigraphy**: Monte Bolca, near Verona, Italy: Ypresian, Eocene, 48.96 to 48.5 Ma[3].

**Fossil tip age**: 48.5 Ma. The node calibration was created such that 97.5% of the probability distribution fell before this date.

†*Eoleiognathus dorsalis*

**Node calibrated:** †*Eoleiognathus dorsalis* is a member of Pan-*Leiognathidae*[17,18] and calibrates the node representing the MRCA of *Chaetodon unimaculatus* and *Gazza minuta* in our analysis. The placement of †*E. dorsalis* in Pan-*Leiognathidae* is supported by the following combination of characters: small, protractile mouth; high, pointed supraoccipital crest; origin of dorsal fin anteriorly shifted; 24 vertebrae; non-oligomerized caudal skeleton; iliac process well developed; and meristic counts of the dorsal and anal fins[17].

**Stratigraphy**: Monte Bolca, near Verona, Italy: Ypresian, Eocene, 48.96 to 48.5 Ma[3].

**Fossil tip age**: 48.5 Ma. The node calibration was created such that 97.5% of the probability distribution fell before this date.

†*Chaetodon ficheuri*

**Node calibrated:** †*Chaetodon ficheuri* is a member of Pan-*Chaetodon*[19] and calibrates the node representing the MRCA of *Chaetodon unimaculatus* and *Forcipiger longirostris* in our analysis. The placement of †*C. ficheuri* in *Chaetodontidae* as a member of Pan-*Chaetodon* is supported by the following combination of characters: supraoccipital crest tall; frontals sutured to epiotics; second infraorbital bone excluded from orbit; supracleithral lateral line canal incomplete medially; 24 vertebrae; ctenoid scales arranged in ascending rows from the lower anterior to upper posterior; predorsal formula 0/0+2/1/1; pelvic fin spine and basipterygium interlocked via two small flanges at proximal end of pelvic spine fused through pelvic articular foramen; lateral line scales disposed as smooth arch below the dorsal fin and extend almost to last dorsal fin ray; caudal fin skeleton structure and composition[19].

**Stratigraphy**: Massive diatomites at Les Planteurs, Raz-el-Aïn, Saint-Denis du Sig, and St. Eugène, Chelif Basin, northwestern Algeria; early Messinian, ~7 to 6 Ma[19].

**Fossil tip age**: 6.0 Ma.

### <u>Supplementary Results and Discussion.</u>

***Taxonomic Findings.-*** Bannerfishes (*Heniochinae*) and butterflyfishes (*Chaetodontinae*) are each resolved as reciprocally monophyletic (Fig. 2). In *Chaetodontinae*, *Prognathodes* is paraphyletic with respect to *Roa* (Fig. 2; Supplementary Fig. 1) as Orange Margin Butterflyfish *P. basabei* Pyle and Kosaki and Twin Butterflyfish *P. geminus* Copus, Pyle, Greene, and Randall are a clade that is resolved as the sister lineage of *Roa* with high node support. The UCE phylogeny resolves the Sixspine Butterflyfish, *Chaetodon oligacanthus* Bleeker, nested within *Chaetodon* and a sister species of a clade containing *Chaetodon aureofasciatus*, *C. octofasciatus*, and *C. rainfordi* (Fig. 2). This taxon was previously classified as *Parachaetodon ocellatus* (Cuvier)[20,21]. Within *Chaetodon*, the phylogenomic analysis resolved Ghana Butterflyfish *C. robustus* as the sister species to all other species in the clade (Fig. 2; Supplementary Fig. 1).

***Feeding Strategy Classification.-*** Species-level feeding strategies (non-coral feeder, facultative corallivore, obligate corallivore) were compiled from the published literature on butterflyfish dietary ecology, primarily field-based bite-rate and stomach-content studies[22–28,28–31]. In some cases where classifications conflicted between sources, we gave precedence to studies based on gut content analysis over those based on field observations of feeding behavior alone, as direct examination of stomach contents provides more definitive evidence of realized diet than inferred feeding activity. The only species for which this created a discrepancy with prior classification was *Heniochus singularius*. This species has previously been classified as obligate corallivore due to the field observations of only two individuals reported by Sano (1989). Due to the location of the species on the tree, the lack of additional observations and stomach content analysis and sources reporting hat this species also consumes algae and non-coral invertebrates we classify the species here as a facultative corallivore. All species, their classification used in this paper and the sources used for each one can be found in Supplementary Table 2.

***Depth Occurrence Data.-*** Depth occurrence range data for each species were obtained from FishBase[32], which reports a minimum and maximum depth of occurrence per species. Average depth was calculated as the midpoint of these two values and used directly as the continuous depth trait in all continuous-trait analyses. For the discrete depth trait analysis, each species was assigned to one of three ordinal categories (shallow, ≤20 m; mid, 21–50 m; deep, >50 m) according to which category its average depth value fell into.

### Supplementary Discussion 1. Phylogeny, Divergence Times and Reef History.

***Congruence with Previous Studies.-*** The topology we inferred is broadly concordant with previous studies based on smaller taxon and sequence datasets[33–36,37].

***Timing of the Crown Ages.-*** The age of the most recent common ancestor (MRCA) of *Chaetodontidae* is 18.16 Ma (95% highest posterior density [HPD] interval: 12.36–26.95 Ma) (Fig. 2), indicating that the original diversification of the extant lineages of *Chaetodontidae* occurred during the early Miocene reef-expansion zenith implicated in the assembly of modern reef biodiversity[38–40], at a timing matched by the origination and diversification of other major reef-fish lineages[38,39]. In addition, we estimated the age of the MRCA of *Chaetodontinae*, which contains the majority of coral-feeding species in the genera *Prognathodes*, *Roa*, and *Chaetodon*, to be 7.78 Ma (95% HPD: 7.13–8.49 Ma) (Fig. 2, Supplementary Tables 2, 3), indicating that most extant chaetodontid species diversity, and in particular the lineage specialized on coral feeding, arose long after the reef-expansion zenith, but in the Late Miocene.

***Prior Estimates of Crown Age.-*** The Early Miocene crown age estimate for *Chaetodontidae* is younger than prior estimates which placed the origin in the Oligocene or Eocene: 28.7 Ma (95% HPD: 18.26–40.0 Ma)[33]; 32.8 Ma (95% HPD 24.9–40.9 Ma)[41]; 36.72 Ma[34] and 40.52 Ma[42]. Similarly, our crown age estimate for *Chaetodontinae* (7.78 Ma) is also younger than prior estimates: 15.4 Ma[33]; 23.9 Ma[41]; 24.0 Ma[34]; and 28.10 Ma[42]. Differences among studies likely reflect variation in taxon sampling, sequence data type (targeted Sanger loci[34,42,37,41] versus phylogenomic datasets[33]), the fossil calibration schemes, and the relaxed-clock implementations (BEAST here versus MCMCTREE[33] or penalized likelihood in r8s[34]).

***Reef Expansion and Contraction.-*** The Early–Mid Miocene (23–11.6 Ma; peak 16–15 Ma) was marked by the largest increase in reef area and thickness of the past 66 million years[40,43], when the Great Indo-Australian Miocene Reef System exceeded modern reefs in both seafloor area and thickness[44,45] and coral lineages diversified across most growth forms[40]. This expansion ended by 10 Ma, when many Indo-Pacific reefs contracted or disappeared amid climate-driven change, shifting regional seascapes toward fragmentation and ecological restructuring[40,44,46].

### Supplementary Discussion 2. Diversification-Rate Analyses.

***Comparison with Other Reef-Fish Lineages.-*** To place the diversification of *Chaetodontinae* in a broader context of coral reef fish lineages, we estimated crown lineage diversification rates across several reef-associated clades under extinction fractions of $\varepsilon=0$ and $\varepsilon=0.9$ (Fig. 3a; Supplementary Table 4). Because the crown ages of these comparative reef fish lineages span more than an order of magnitude (6.7–85.25 Ma; Supplementary Table 4), we interpret each rate as a standardized richness–age summary under shared assumptions rather than as evidence that diversification proceeded at a constant rate throughout any individual clade's history. Under both extinction fractions, *Chaetodontinae* showed the highest estimated crown diversification rate of all clades examined (0.515 $Ma^{-1}$ at $\varepsilon=0$ and 0.312 $Ma^{-1}$ at $\varepsilon=0.9$), exceeding rates for other species-rich reef-fish radiations including *Cirrhilabrus* (fairy wrasses), *Blenniidae* (blennies), *Pomacentridae* (clownfishes and damselfishes), *Cirrhitidae* (hawkfishes), and *Gobiidae* (gobies) (Fig. 3a; Supplementary Table 4). Although absolute rate estimates decreased under high extinction fraction, *Chaetodontinae* remained an outlier relative to other reef lineages across the full comparison (Fig. 3a). Notably, only *Cirrhilabrus* and *Blenniidae* approach the rate of lineage diversification in *Chaetodontinae* under ε=0, and *Chaetodontinae* remains the fastest-diversifying clade using a high fraction of extinction (ε=0.9), indicating that elevated diversification is robust to extinction-fraction assumptions across a broad comparative context.

***Species-Richness Tests.-*** In addition, the species diversity of *Chaetodontinae* significantly exceeded the richness expected under the lineage diversification rate of 27 of the 29 lineages of reef fishes assuming no extinction (ε=0: all lineages except *Cirrhilabrus* and *Blenniidae*; all upper 95% CI bounds ≤63.3 species for every other significant clade comparison), and 26 of 29 under high extinction (ε=0.9: all except *Cirrhilabrus, Pomacentridae,* and *Blenniidae*) (Extended Data Table 1). *Cirrhilabrus* and *Blenniidae* were the only two lineages of reef fishes for which the species diversity of *Chaetodontinae* did not significantly exceed the expected number of species under either extinction scenario (*Cirrhilabrus:* 95% CI upper bound = 243.7 species at ε=0, 263.9 at ε=0.9; *Blenniidae:* 152.7 species at ε=0, 352.0 at ε=0.9), consistent with these two lineages having the highest and second-highest estimated diversification rates among the 29 reef fish lineages (*Cirrhilabrus* r=0.486, *Blenniidae* r=0.427 at ε=0). *Pomacentridae* additionally became non-significant under the high-extinction scenario alone (95% CI upper bound = 179.0 at ε=0.9, versus 110 species of *Chaetodontinae*), despite being significant at ε=0 (upper bound = 63.3). Overall, the diversification rate of *Chaetodontinae* is high relative to most reef-fish lineages examined, regardless of extinction assumption (Extended Data Table 1).

***Background Diversification Rates.-*** We additionally compared the expected species diversity of *Chaetodontinae, Heniochinae,* and 29 other reef fish lineages using the background diversification rates of the inclusive clades *Percomorpha*, *Eupercaria*, *Perciformes*, and *Acanthuriformes*[47]. Under the background diversification rate of *Percomorpha* (r=0.081 $Ma^{-1}$ at ε=0 and r=0.066 $Ma^{-1}$ at ε=0.9), we found that *Chaetodontinae* are more species rich than expected, with observed richness falling above the upper bound of both the ε=0 and ε=0.9

confidence intervals (Fig. 3b, Extended Data Fig. 4, Supplementary Table 5a). Other young, comparatively species-rich lineages of reef fishes, including *Blenniidae*, *Pomacentridae*, *Gobiidae*, and *Julidinae* are also more species rich given the background diversification of *Percomorpha* (Fig. 3b, Extended Data Fig. 4, Supplementary Table 5a). Several older lineages of reef fishes with lower estimated diversification rates, including *Syngnathidae* (seahorses), *Antennariidae* (frogfishes) and *Holocentridae* (squirrelfishes) fell within the expected range of species richness (Fig. 3b, Supplementary Table 5a). Similarly, the species diversity of *Chaetodontinae* exceeded the upper ε=0.9 bound under the background diversification rates of *Eupercaria* (r=0.093 $Ma^{-1}$ at ε=0 and r=0.074 $Ma^{-1}$ at ε=0.9), *Perciformes* (r=0.145 $Ma^{-1}$ at ε=0 and r=0.112 $Ma^{-1}$ at ε=0.9), and *Acanthuriformes* (r=0.088 $Ma^{-1}$ at ε=0 and r=0.067 $Ma^{-1}$ at ε=0.9) (Extended Data Fig. 4, Supplementary Table 5b–d).

***Time-Varying Rates (TESS-CoMET).-*** The estimated rate of extinction was low and constant through the analysis (posterior mean 0.012–0.025 $Ma^{-1}$; Extended Data Fig. 2b), although over most of the time axis its posterior is close to the prior, so extinction is poorly constrained, no change in the extinction rate and no mass extinction was supported. Net diversification then declined to approximately 0.05 $Ma^{-1}$ at the present in the analysis with positive support for a shift in the speciation rate at approximately 0.7–0.5 Ma (2 ln BF = 5.7 in *Chaetodontidae*; Fig. 2b, Extended Data Fig. 2a,c). Because time-dependent diversification methods can be sensitive to incomplete sampling, we interpret the slowdown of lineage accumulation towards the present as an artifact of the diversification model, as there are no nodes in the phylogeny younger than 500,000 years (the mean age of the youngest node in the relaxed clock analysis is 0.53 Ma) and the mean ages of only four nodes, all within *Chaetodontinae*, are younger than 700,000 years. Because TESS-CoMET fits a single rate to all lineages at a given time, it cannot say which lineages carry the increase; however, a hidden-state model can.

### Supplementary Discussion 3. Feeding Ecology and Ancestral-State Reconstructions.

***Dietary Composition of Corallivores.-*** Among species classified as obligate coral-feeders, scleractinian (hard) branching corals universally dominated the diet: most (e.g., *Chaetodon austriacus*, *C. baronessa*, *C. bennetti*, *C. ornatissimus*, *C. reticulatus*, *C. trifascialis*, *C.*

*trifasciatus*) derived 100% of feeding records from scleractinian corals, with many others exceeding 90% (*C. lunulatus*, 97%; *C. plebeius*, 96%; *C. rainfordi*, 93%) (Fig. 4a, Supplementary Tables 2 and 3). Several facultative corallivore species, including *C. auriga*, *C. lunula*, *C. vagabundus*, *C. kleinii*, *C. argentatus*, *Heniochus chrysostomus*, and *H. varius*, relied more heavily on non-coral trophic sources such as plankton, algae, and other invertebrate prey (Fig. 4a, Supplementary Tables 2 and 3).

***Coral Genus Use.-*** When analyzing the diversity of corals comprising the diet of 19 obligate corallivores, Friedman's test revealed a significant overall difference in coral genus use among species of *Chaetodon* ($\chi^2$=10.8, df=3, p=0.013), though the effect size was modest (W=0.211, n=17) (Fig. 4b). Post-hoc pairwise comparisons showed that two contrasts remained significant after Holm-Bonferroni correction: *Acropora* was consumed at higher rates than *Montipora* (p_holm=0.00064), and *Porites* at higher rates than *Montipora* (p_holm=0.010) (Fig. 4b). None of the other four genus pairs were significant after correction. Including the three non-obligate species in the source table (n=20) gave the same overall result ($\chi^2$=10.2, p=0.017) and the same *Acropora-Montipora* difference (p_holm=0.00038), but the *Porites-Montipora* difference was not significant (p_holm=0.064; Supplementary Table 7).

***Ancestral States of Diet.-*** Stochastic mapping inferred a mean of 30.96 diet changes, dominated by transitions from facultative corallivory to non-coral feeding (mean = 16.6). Transitions between facultative and obligate corallivory were bidirectional (means = 4.66 and 3.69). Transitions from obligate corallivory to non-coral feeding were rare (mean=0.7) (Extended Data Fig. 5a).

***Depth-Category Evolution.-*** Habitat depth-category evolution favored an ARD model (AIC = 242.77) with weak root-state resolution (shallow = 0.338, mid = 0.334, extreme = 0.328). Stochastic mapping inferred high lability (mean = 128.47 changes), dominated by shallow–mid depth transitions (means = 52.8 and 45.2; Extended Data Fig. 5c). Direct shallow-to-extreme depth transitions were not reconstructed, whereas there were extreme-to-shallow depth transitions (mean = 3.59), indicating deep-habitat colonization proceeded through the mid-depth category (mid-to-extreme = 18.4).

***Tooth Characters.-*** Under an equal-rates transition model, the ancestral state at the most recent common ancestor of *Chaetodontidae* was reconstructed as having a basic dental pattern, which is present in all species, and lacking each of the derived dental character states, with high confidence in every case ($P$ = 0.994 for Character 18:2, 0.994 for 18:3, 0.969 for 18:4, 0.997 for 18:5, 0.971 for 18:6, and 0.997 for 18:7). This indicates that each of these six tooth characters represents a derived condition that arose later within *Chaetodontidae*, rather than being part of the ancestral state for the lineage. Three of the least complex derived character states (18:2–18:4) are restricted to species outside *Chaetodon*: angled teeth in more than three rows per band (18:2) and nearly vertical teeth in six bands (18:3) occur in *Chelmon* and *Coradion*, and short, straight teeth in two bands (18:4) occur in *Hemitaurichthys* and *Heniochus*. All species with these states are facultative corallivores or non-coral feeders. By contrast, three more complex (18:5-18:7), coral-associated character states (teeth organized in more narrow and numerous bands, teeth straight with descending process edentulous, and very fine teeth coalescing into a brush) are found almost exclusively in species of *Chaetodon* that are obligate corallivorous (Extended Data Fig. 6).

**<u>Supplementary Discussion 4. Morphological Disparity and Ecological Covariation.</u>**

***Multivariate Morphology and Ecology.-*** To test whether phenotypic diversification is tied to ecological differentiation, as expected under ecological adaptive radiation[48], we quantified morphological variation and asked whether it covaries with depth and diet, expecting feeding-related (jaw) traits to be most strongly associated with diet. Principal component analysis of the size-corrected traits showed that the first two axes explained 69% of the variance in the full trait set (PC1 53%, PC2 16%) and 84% in each of the jaw and body sets (Extended Data Fig. 7a). Two-block partial least squares (PLS) analysis supported a significant multivariate correlation between the full morphological trait set and combined ecological patterns of depth and diet ($r=0.390$, $R^2=0.152$, permutation $P=0.008$, $n=82$; Extended Data Table 3). This correlation was driven by body-shape traits ($r=0.398$, $R^2=0.158$, $P=0.005$), whereas jaw traits were weaker and non-significant ($r=0.264$, $R^2=0.070$, $P=0.152$) (Extended Data Fig. 7b; Extended Data Table 3). Phylogenetic generalized least squares (pGLS) similarly supported associations for the full trait set ($R^2$=0.079, P=0.011, λ=0.527) and body traits ($R^2$=0.076, P=0.012, λ=0.511), but not for jaw traits ($R^2$=0.039, P=0.076, λ=0.606) (Extended Data Table 3).

***Depth and Diet Tested Separately.-*** When determining whether depth or diet was independently responsible for these associations, we repeated the analysis with depth and corallivory tested as separate variables. Depth alone was significantly associated with all three morphological trait sets (all traits, body and jaw; the jaw set comprises head depth, lower jaw length and mouth width), in both the permutation-based (all traits: r=0.398, P=0.010; body: r=0.402, P=0.002; jaw: r=0.275, P=0.047) and phylogenetically-corrected tests (all traits: $R^2$=0.130, P<0.001; body: $R^2$=0.130, P<0.001; jaw: $R^2$=0.067, P=0.019), indicating that depth alone accounts for the jaw-morphology association that was undetectable when combined with diet. Corallivory alone, by contrast, showed no phylogenetically corrected association with any morphological traits (all pGLS P>0.25); the one nominally significant permutation result (body traits, r=0.292, P=0.032) did not survive phylogenetic correction (pGLS $R^2$=0.001, P=0.786, λ=1), indicating this signal reflects phylogenetic clustering of diet and body shape rather than a robust independent relationship (Extended Data Table 3). These results indicate that the combined ecospace association reported above is driven predominantly by depth rather than diet.

***Rates of Trait Evolution.-*** Brownian motion rate estimates ($\sigma^2$) for the seven morphological traits, all log-transformed linear measurements corrected for body size, revealed substantial rate heterogeneity. Lower jaw length ($\sigma^2$=0.039) and mouth width ($\sigma^2$=0.036) had the highest rates, exceeding all four body-shape traits ($\sigma^2$=0.0018–0.0092), indicating relatively high rates of evolution in specific feeding-related dimensions despite the limited jaw morphology and ecology association in regression-based tests (Supplementary Table 6). Together, these results suggest that depth-related habitat use is a primary axis linking morphology to ecology across *Chaetodontidae*, whereas the feeding-related traits examined here may be evolving rapidly along trophic or microhabitat dimensions that our diet categories do not fully capture, such as prey type breadth within corallivory, foraging mode, and fine-scale reef structure.

***Lineage Accumulation and Morphological Disparity.-*** Across reef fish lineages, species richness is not associated with greater body shape disparity, suggesting that rapid lineage accumulation and morphological diversification can be decoupled at macroevolutionary scales[49].

**Supplementary Discussion 5. Jaw Mechanics and Digestive Specialization.**

***The Intramandibular Joint.-*** Among candidate evolutionary innovations, the intramandibular joint (IMJ) is a plausible key innovation: adaptive radiation[48] theory expects such traits to increase performance in a newly exploited habitat or resource. Although we did not measure performance, the IMJ bears on the performance-utility criterion for ecological adaptive radiation because earlier work links this discrete phenotypic change to a shift in feeding mechanics and resource use[50,51]. More specifically, the IMJ facilitates a biomechanical shift from ram-suction feeding toward benthic, coral-based foraging, with maximal joint flexion in species that scrape mucus from the surface of corals[50]. Its gain-only pattern in ancestral character reconstruction is consistent with a key innovation model (Fig. 5), in which access to a resource or foraging mode reduces the selective incentive to revert to an ancestral, less effective configuration[52]. This interpretation is reinforced by the tempo of phenotypic evolution: lower jaw length and mouth width evolved at roughly ten times the rate of body-shape traits in *Chaetodontidae* (Supplementary Table 6).

***Digestive Physiology and Jaw Protrusion.-*** Additional morphological features that are consistent with specialization on coral-derived resources include long thin teeth organized into pads or peripheral placement of teeth facilitating various corallivore feeding modes[53,54] and elongation of the digestive tract associated with coral-based diets[55] (Fig. 1). Elevated rates of gut-length evolution in *Chaetodon* lineages occur alongside relatively slower cranial diversification[51], suggesting that once the IMJ enables coral feeding, subsequent specialization may proceed through digestive physiology as much as through further cranial remodeling. *Chaetodontidae* can bite with fully protruded jaws, a novelty shared with *Pomacanthidae* enabled by extreme jaw protrusion and mandibular extension at the IMJ[50,56] (Fig. 1). Because only a minority of pomacanthids consume coral and none are obligate corallivores, this shared mechanism is unlikely to be sufficient on its own; instead, the disproportionate reliance on coral as a dietary resource within *Chaetodon* appears to reflect lineage-specific ecological and evolutionary amplification of a broader ancestral biomechanical capability present at the origin of *Chaetodon*.

**References.**

1. Carnevale, G., Pietsch, T. W., Bonde, N., Leal, M. E. C. & Marramà, G. †Neilpeartia ceratoi, gen. et sp. nov., a new frogfish from the Eocene of Bolca, Italy. *Journal of Vertebrate Paleontology* **40**, e1778711 (2020).
2. Carnevale, G. & Pietsch, T. W. An Eocene Frogfish from Monte Bolca, Italy: The Earliest Known Skeletal Record for the Family. *Palaeontology* **52**, 745–752 (2009).
3. Friedman, M. & Carnevale, G. The Bolca Lagerstätten: shallow marine life in the Eocene. *Journal of the Geological Society* **175**, 569–579 (2018).
4. Carnevale, G. & Pietsch, T. W. †Caruso, a new genus of anglerfishes from the Eocene of Monte Bolca, Italy, with a comparative osteology and phylogeny of the teleost family Lophiidae. *Journal of Systematic Palaeontology* **10**, 47–72 (2012).
5. Bannikov, A. The first discovery of an anglerfish (Teleostei, Lophiidae) in the Eocene of the Northern Caucasus. *Paleonotological Journal* **38**, 420–425 (2004).
6. Close, R. A., Johanson, Z., Tyler, J. C., Harrington, R. C. & Friedman, M. Mosaicism in a new Eocene pufferfish highlights rapid morphological innovation near the origin of crown tetraodontiforms. *Palaeontology* **59**, 499–514 (2016).
7. Bannikov, A. & Tyler, J. A new species of the pufferfish Eotetraodon (Tetraodontiformes, Tetraodontidae) from the Eocene of the Northern Caucasus. *Paleontological Journal* **42**, 526–530 (2008).
8. Ghezelayagh, A. *et al.* Prolonged morphological expansion of spiny-rayed fishes following the end-Cretaceous. *Nat Ecol Evol* **6**, 1211–1220 (2022).
9. Near, T. J. *et al.* Resolution of ray-finned fish phylogeny and timing of diversification. *Proc. Natl. Acad. Sci. U.S.A.* **109**, 13698–13703 (2012).

10. Tyler, J. C. & Bannikov, A. F. Relationships of the fossil and recent genera of rabbitfishes (Acanthuroidei: Siganidae). http://repository.si.edu/xmlui/handle/10088/2000 (1997).

11. Gradstein, F. M., Ogg, J. G., Schmitz, M. & Ogg, G. *The Geologic Time Scale 2020*. (Elsevier Science, Amsterdam, The Netherlands, 2021).

12. Bannikov, A. & Tyler, J. A new species of the luvarid fish genus †Avitoluvarus (Acanthuroidei, Perciformes) from the Eocene of the Caucasus in Southwest Russia. *Proceedings of the Biological Society of Washington* **114**, 579–588 (2001).

13. Bannikov, A. F. & Tyler, J. C. *Phylogenetic Revision of the Fish Families Luvaridae and †Kushlukiidae (Acanthuroidei), with a New Genus and Two New Species of Eocene Luvarids*. (Smithsonian Institution, 1995).

14. Prokofiev, A. M. A remarkable new genus of Carangidae fron1 the Upper Paleocene of Turkmenistan (Osteichthyes: Perciformes). https://www.zin.ru/Journals/zsr/content/2002/zr_2002_11_1_Prokofiev_2.pdf (2002).

15. Siqueira, A. C., Bellwood, D. R. & Cowman, P. F. Historical biogeography of herbivorous coral reef fishes: The formation of an Atlantic fauna. *Journal of Biogeography* **46**, 1611–1624 (2019).

16. Tyler, J. & Bannikov, A. A new species of the surgeon fish genus Tauichthys from the Eocene of Monte Bolca, Italy (Perciformes, Acanthuridae). *Bollettino del Museo Civico di Storia Naturale di Verona Geologia Paleontologia Preistoria* **24**, 29–36 (2000).

17. Bannikov, A. The new genus Eoleiognathus for the percoid fish Pygaeus dorsalis Agassiz from the Eocene of Bolca in northern Italy, a putative ponyfish (Perciformes, Leoignathidae). *Studi e ricerche sui giacimenti terziari di Bolca* **15**, (2014).

18. Kovalchuk, O. M., Świdnicka, E. & Stefaniak, K. Early Miocene Ponyfishes (Acanthuriformes, Leiognathidae) of the Carpathian Basin. *Paleontol. J.* **55**, 421–428 (2021).

19. Carnevale, G. Morphology and biology of the Miocene butterflyfish Chaetodon ficheuri (Teleostei: Chaetodontidae). *Zoological Journal of the Linnean Society* **146**, 251–267 (2006).

20. Fricke, R., Eschmeyer, W. N. & Van der Laan, R. Eschmeyer's catalog of fishes: genera, species, references. http://researcharchive.calacademy.org/research/ichthyology/catalog/fishcatmain.asp (2026).

21. Smith, Wm. L., Webb, J. F. & Blum, S. D. The evolution of the laterophysic connection with a revised phylogeny and taxonomy of butterflyfishes (Teleostei: Chaetodontidae). *Cladistics* **19**, 287–306 (2003).

22. Pratchett, M. S. Dietary selection by coral-feeding butterflyfishes (Chaetodontidae) on the Great Barrier Reef, Australia. (2007).

23. Pratchett, M. S. Dietary overlap among coral-feeding butterflyfishes (Chaetodontidae) at Lizard Island, northern Great Barrier Reef. *Marine Biology* **148**, 373–382 (2005).

24. Cole, A. J., Pratchett, M. S. & Jones, G. P. Diversity and functional importance of coral-feeding fishes on tropical coral reefs. *Fish and Fisheries* **9**, 286–307 (2008).

25. Sano, M. Feeding habits of Japanese butterfyfishes (Chaetodontidae). *Environ Biol Fish* **25**, 195–203 (1989).

26. Bouchon-Navaro, Y. Partitioning of food and space resources by chaetodontid fishes on coral reefs. *Journal of Experimental Marine Biology and Ecology* **103**, 21–40 (1986).

27. Bouchon-Navaro, Y. & Bouchon, C. Correlations between chaetodontid fishes and coral communities of the Gulf of Aqaba (Red Sea). *Environ Biol Fish* **25**, 47–60 (1989).

28. Harmelin-Vivien, M. L. & Bouchon-Navaro, Y. Feeding diets and significance of coral feeding among Chaetodontid fishes in Moorea (French Polynesia). *Coral Reefs* **2**, 119–127 (1983).

29. Nagelkerken, I., Van Der Velde, G., Wartenbergh, S. L. J., Nugues, M. M. & Pratchett, M. S. Cryptic dietary components reduce dietary overlap among sympatric butterflyfishes (Chaetodontidae). *Journal of Fish Biology* **75**, 1123–1143 (2009).

30. Randall, J. E. Food habits of reef fishes of the West Indies. *Studies in Tropical Oceanography* **5**, 665–847 (1967).

31. Pratchett, M. S., Berumen, M. L. & Kapoor, B. G. *Biology of Butterflyfishes*. *Biology of Butterflyfishes* 352 (CRC Press, 2013).

32. Froese, R. & Pauly, D. FishBase. https://www.fishbase.org/.

33. DiBattista, J. D. *et al.* Ice ages and butterflyfishes: Phylogenomics elucidates the ecological and evolutionary history of reef fishes in an endemism hotspot. *Ecology and Evolution* **8**, 10989–11008 (2018).

34. Fessler, J. L. & Westneat, M. W. Molecular phylogenetics of the butterflyfishes (Chaetodontidae): Taxonomy and biogeography of a global coral reef fish family. *Molecular Phylogenetics and Evolution* **45**, 50–68 (2007).

35. Cowman, P. F. & Bellwood, D. R. Coral reefs as drivers of cladogenesis: expanding coral reefs, cryptic extinction events, and the development of biodiversity hotspots. *J. Evol. Biol.* **24**, 2543–2562 (2011).

36. Ghezelayagh, A. *et al.* Prolonged morphological expansion of spiny-rayed fishes following the end-Cretaceous. *Nat Ecol Evol* **6**, 1211–1220 (2022).

37. Hodge, J. R., van Herwerden, L. & Bellwood, D. R. Temporal evolution of coral reef fishes: global patterns and disparity in isolated locations. *Journal of Biogeography* **41**, 2115–2127 (2014).

38. Tian, S. Y. *et al.* Cenozoic history of the tropical marine biodiversity hotspot. *Nature* **632**, 343–349 (2024).

39. Brownstein, C. D. *et al.* Phylogenomics establishes an Early Miocene reconstruction of reef vertebrate diversity. *Science Advances* **11**, eadu6149 (2025).

40. Siqueira, A. C., Kiessling, W., Raja, N. B. & Bellwood, D. R. The rise and fall of the world's greatest marine biodiversity hotspot. *Science Advances* **12**, eaec7264 (2026).

41. Bellwood, D. R. *et al.* Evolutionary history of the butterflyfishes (f: Chaetodontidae) and the rise of coral feeding fishes. *J. Evol. Biol.* **23**, 335–349 (2010).

42. Rabosky, D. L. *et al.* An inverse latitudinal gradient in speciation rate for marine fishes. *Nature* **559**, 392–395 (2018).

43. Kiessling, W., Simpson, C. & Foote, M. Reefs as Cradles of Evolution and Sources of Biodiversity in the Phanerozoic. *Science* **327**, 196–198 (2010).

44. McCaffrey, J. C., Wallace, M. W. & Gallagher, S. J. A Cenozoic Great Barrier Reef on Australia's North West shelf. *Global and Planetary Change* **184**, 103048 (2020).

45. Mathew, M. *et al.* The emergence of Miocene reefs in South China Sea and its resilient adaptability under varying eustatic, climatic and oceanographic conditions. *Sci Rep* **10**, 7141 (2020).

46. Leprieur, F., Pellissier, L., Mouillot, D. & Gaboriau, T. Influence of historical changes in tropical reef habitat on the diversification of coral reef fishes. *Sci Rep* **11**, 20731 (2021).

47. Near, T. J. & Thacker, C. E. Phylogenetic Classification of Living and Fossil Ray-Finned Fishes (Actinopterygii). *Bull Peabody Museum Natural History* **65**, 3–302 (2024).

48. Schluter, D. *The Ecology of Adaptive Radiation*. (Oxford University Press, Oxford, New York, 2000).

49. Price, S. A., Claverie, T., Near, T. J. & Wainwright, P. C. Phylogenetic insights into the history and diversification of fishes on reefs. *Coral Reefs* **34**, 997–1009 (2015).

50. Konow, N., Bellwood, D. R., Wainwright, P. C. & Kerr, A. M. Evolution of novel jaw joints promote trophic diversity in coral reef fishes. *Biol J Linn Soc* **93**, 545–555 (2008).

51. Konow, N., Price, S., Abom, R., Bellwood, D. & Wainwright, P. Decoupled diversification dynamics of feeding morphology following a major functional innovation in marine butterflyfishes. *Proc. R. Soc. B* **284**, 20170906 (2017).

52. Miller, A. H., Stroud, J. T. & Losos, J. B. The ecology and evolution of key innovations. *Trends in Ecology & Evolution* **38**, 122–131 (2023).

53. Motta, P. J. Dentition patterns among Pacific and Western Atlantic butterflyfishes (Perciformes, Chaetodontidae): relationship to feeding ecology and evolutionary history. *Environ Biol Fish* **25**, 159–170 (1989).

54. Marrable, I., Peoples, N., Kolmann, M. A., Huie, J. M. & Figueira, W. F. The Evolutionary History and Morphological Divergence of Corallivorous Fishes on Coral Reefs. *Integr Org Biol* **8**, obag032 (2026).

55. Berumen, M. L., Pratchett, M. S. & Goodman, B. A. Relative gut lengths of coral reef butterflyfishes (Pisces: Chaetodontidae). *Coral Reefs* **30**, 1005–1010 (2011).

56. Motta, P. J. Functional morphology of the head of the inertial suction feeding butterflyfish, Chaetodon miliaris (Perciformes, Chaetodontidae). *Journal of Morphology* **174**, 283–312 (1982).

**<u>Supplementary Figure/ Tables.</u>**

**Supplementary Figure 1a. Maximum-Likelihood Phylogeny of *Chaetodontidae* and Outgroup Lineages (panel 1 of 5).** Maximum-likelihood tree inferred in IQ-TREE from a concatenated multi-locus dataset, sampling 260 *Chaetodontidae* specimens plus outgroups. Node support is shown as bootstrap support (BS: light grey circles, 72–92%; dark grey circles, 93–99%; unlabeled nodes, 100%); branch lengths are in substitutions per site. This panel shows the outgroup lineages and the root of the tree; it continues in Supplementary Figure 1b. The inset thumbnail shows the full tree, with the region displayed on this panel highlighted in grey.

**Supplementary Figure 1b. Maximum-Likelihood Phylogeny of *Chaetodontidae* and Outgroup Lineages (panel 2 of 5).** Continued from Supplementary Figure 1a; continues in Supplementary Figure 1c. Node support and branch length conventions as in Supplementary Figure 1a. The inset thumbnail shows the full tree, with the region displayed on this panel highlighted in grey.

**Supplementary Figure 1c. Maximum-Likelihood Phylogeny of *Chaetodontidae* and Outgroup Lineages (panel 3 of 5).** Continued from Supplementary Figure 1b; continues in Supplementary Figure 1d. Node support and branch length conventions as in Supplementary Figure 1a. The inset thumbnail shows the full tree, with the region displayed on this panel highlighted in grey.

**Supplementary Figure 1d. Maximum-Likelihood Phylogeny of *Chaetodontidae* and Outgroup Lineages (panel 4 of 5).** Continued from Supplementary Figure 1c; continues in Supplementary Figure 1e. Node support and branch length conventions as in Supplementary Figure 1a. The inset thumbnail shows the full tree, with the region displayed on this panel highlighted in grey.

**Supplementary Figure 1e. Maximum-Likelihood Phylogeny of *Chaetodontidae* and Outgroup Lineages (panel 5 of 5).** Continued from Supplementary Figure 1d, showing the genus *Chaetodon* through to the final tips of the tree. Node support and branch length conventions as in Supplementary Figure 1a. The inset thumbnail shows the full tree, with the region displayed on this panel highlighted in grey.

**Supplementary Figure 2. Continuous Ancestral Reconstruction of Average Depth Across *Chaetodontidae*.** Maximum-likelihood ancestral state reconstruction of average depth (m), mapped as a continuous trait along the branches of the *Chaetodontidae* phylogeny. Branch color indicates the reconstructed depth value at each point on the tree, from shallow (yellow, 5 m) to deep (dark purple, 165 m).

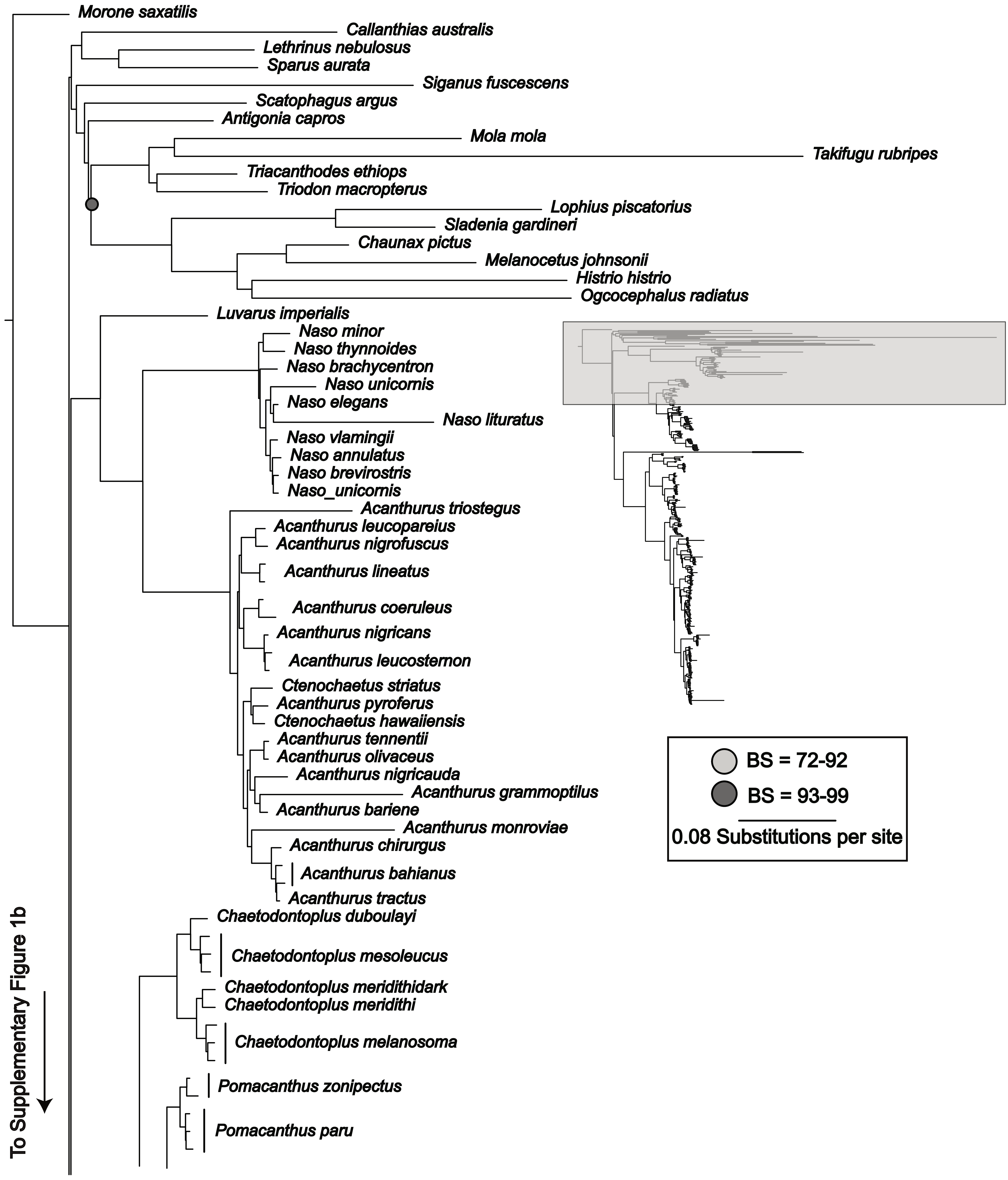
Morone saxatilis
Callanthias australis
Lethrinus nebulosus
Sparus aurata
Siganus fuscescens
Scatophagus argus
Antigonia capros
Mola mola
Takifugu rubripes
Triacanthodes ethiops
Triodon macropterus
Lophius piscatorius
Sladenia gardineri
Chaunax pictus
Melanocetus johnsonii
Histrio histrio
Ogcocephalus radiatus
Luvarus imperialis
Naso minor
Naso thynnoides
Naso brachycentron
Naso unicornis
Naso elegans
Naso lituratus
Naso vlamingii
Naso annulatus
Naso brevirostris
Naso_unicornis
Acanthurus triostegus
Acanthurus leucopareius
Acanthurus nigrofuscus
Acanthurus lineatus
Acanthurus coeruleus
Acanthurus nigricans
Acanthurus leucosternon
Ctenochaetus striatus
Acanthurus pyroferus
Ctenochaetus hawaiiensis
Acanthurus tennentii
Acanthurus olivaceus
Acanthurus nigricauda
Acanthurus grammoptilus
Acanthurus bariene
Acanthurus monroviae
Acanthurus chirurgus
Acanthurus bahianus
Acanthurus tractus
Chaetodontoplus duboulayi
Chaetodontoplus mesoleucus
Chaetodontoplus meridithidark
Chaetodontoplus meridithi
Chaetodontoplus melanosoma
Pomacanthus zonipectus
Pomacanthus paru
BS = 72-92
BS = 93-99
0.08 Substitutions per site
To Supplementary Figure 1b

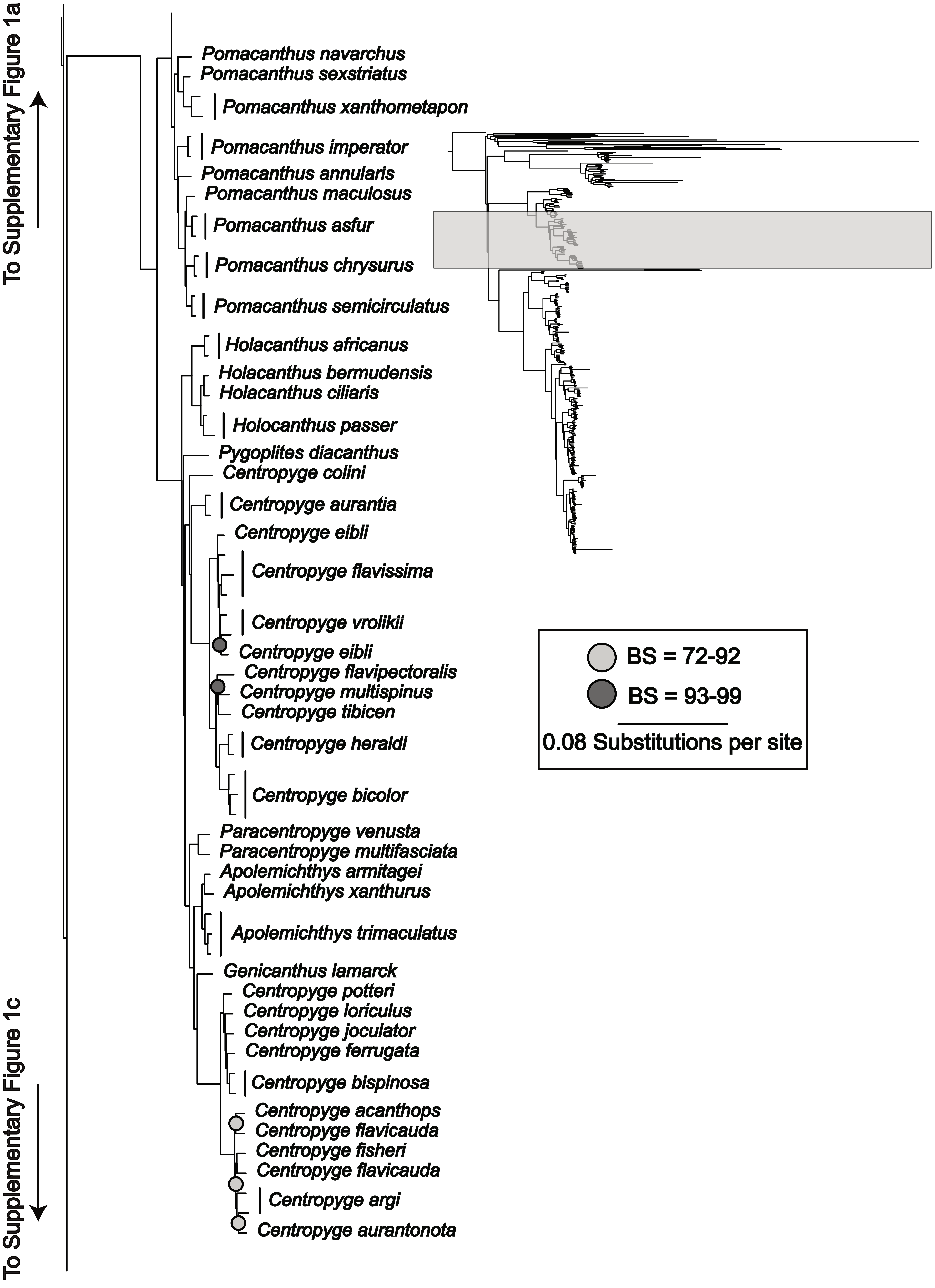

To Supplementary Figure 1a
Pomacanthus navarchus
Pomacanthus sexstriatus
Pomacanthus xanthometapon
Pomacanthus imperator
Pomacanthus annularis
Pomacanthus maculosus
Pomacanthus asfur
Pomacanthus chrysurus
Pomacanthus semicirculatus
Holacanthus africanus
Holacanthus bermudensis
Holacanthus ciliaris
Holocanthus passer
Pygoplites diacanthus
Centropyge colini
Centropyge aurantia
Centropyge eibli
Centropyge flavissima
Centropyge vrolikii
Centropyge eibli
Centropyge flavipectoralis
Centropyge multispinus
Centropyge tibicen
Centropyge heraldi
Centropyge bicolor
Paracentropyge venusta
Paracentropyge multifasciata
Apolemichthys armitagei
Apolemichthys xanthurus
Apolemichthys trimaculatus
Genicanthus lamarck
Centropyge potteri
Centropyge loriculus
Centropyge joculator
Centropyge ferrugata
Centropyge bispinosa
Centropyge acanthops
Centropyge flavicauda
Centropyge fisheri
Centropyge flavicauda
Centropyge argi
Centropyge aurantonota
BS = 72-92
BS = 93-99
0.08 Substitutions per site
To Supplementary Figure 1c

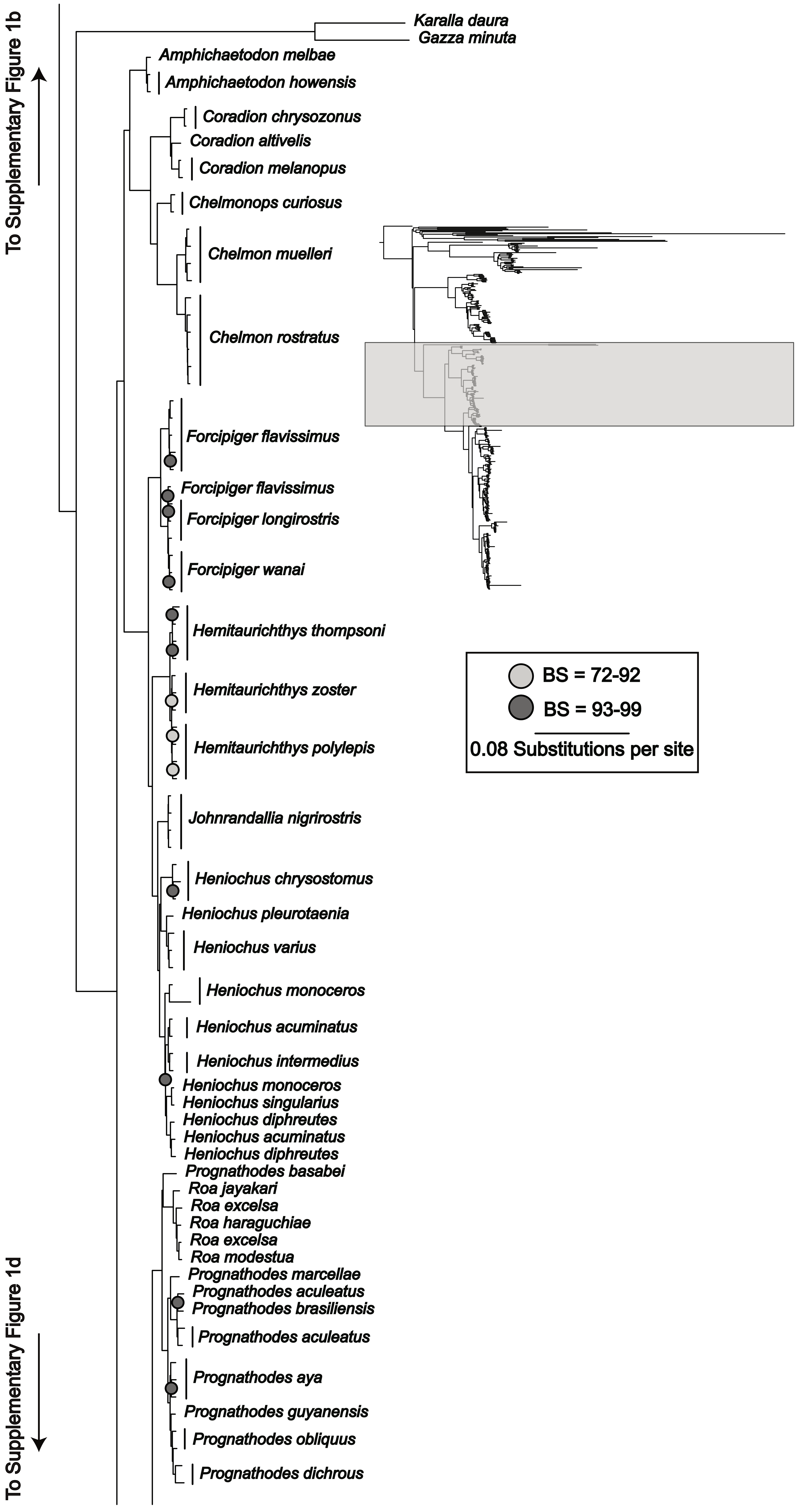
To Supplementary Figure 1b
Karalla daura
Gazza minuta
Amphichaetodon melbae
Amphichaetodon howensis
Coradion chrysozonus
Coradion altivelis
Coradion melanopus
Chelmonops curiosus
Chelmon muelleri
Chelmon rostratus
Forcipiger flavissimus
Forcipiger flavissimus
Forcipiger longirostris
Forcipiger wanai
Hemitaurichthys thompsoni
Hemitaurichthys zoster
Hemitaurichthys polylepis
Johnrandallia nigrirostris
Heniochus chrysostomus
Heniochus pleurotaenia
Heniochus varius
Heniochus monoceros
Heniochus acuminatus
Heniochus intermedius
Heniochus monoceros
Heniochus singularius
Heniochus diphreutes
Heniochus acuminatus
Heniochus diphreutes
Prognathodes basabei
Roa jayakari
Roa excelsa
Roa haraguchiae
Roa excelsa
Roa modestua
Prognathodes marcellae
Prognathodes aculeatus
Prognathodes brasiliensis
Prognathodes aculeatus
Prognathodes aya
Prognathodes guyanensis
Prognathodes obliquus
Prognathodes dichrous
BS = 72-92
BS = 93-99
0.08 Substitutions per site
To Supplementary Figure 1d

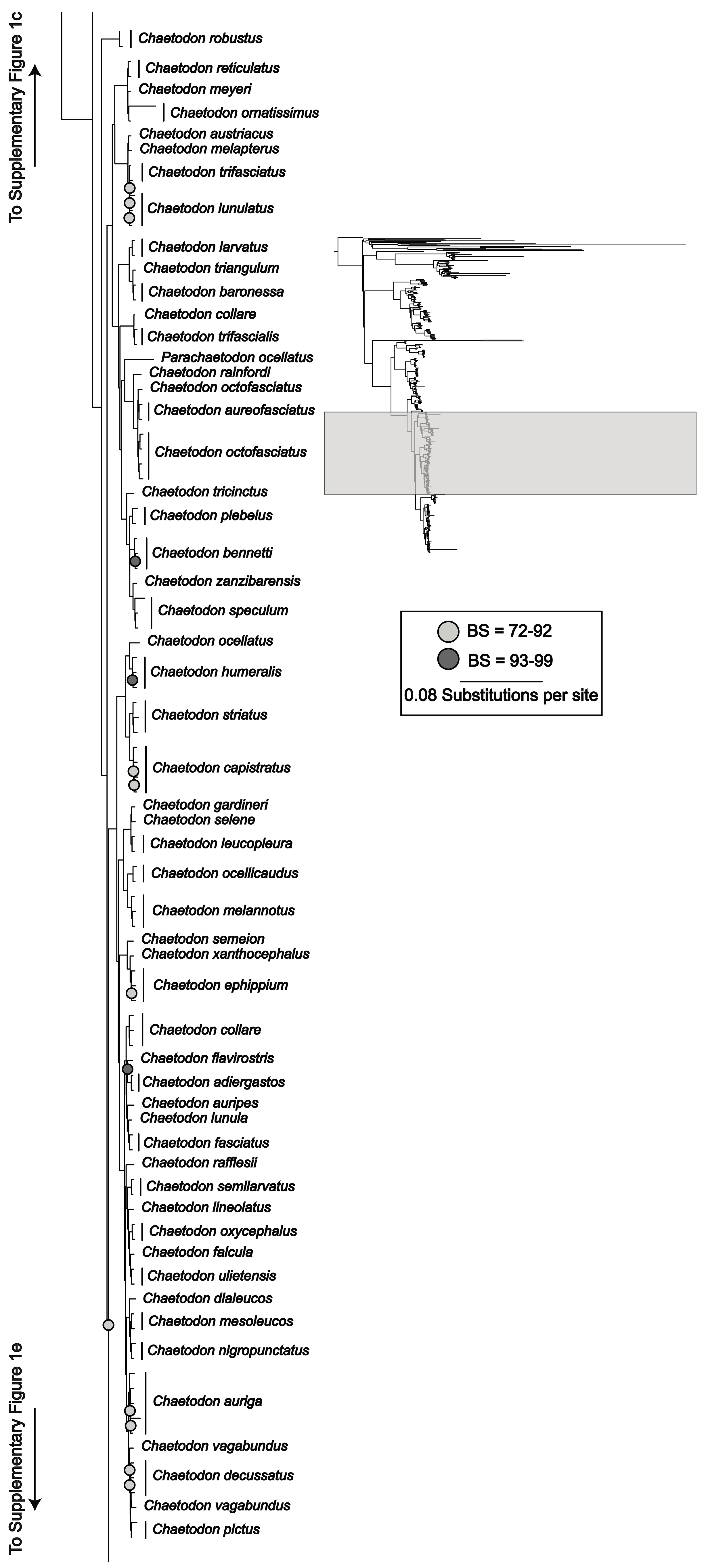
To Supplementary Figure 1c
Chaetodon robustus
Chaetodon reticulatus
Chaetodon meyeri
Chaetodon ornatissimus
Chaetodon austriacus
Chaetodon melapterus
Chaetodon trifasciatus
Chaetodon lunulatus
Chaetodon larvatus
Chaetodon triangulum
Chaetodon baronessa
Chaetodon collare
Chaetodon trifascialis
Parachaetodon ocellatus
Chaetodon rainfordi
Chaetodon octofasciatus
Chaetodon aureofasciatus
Chaetodon octofasciatus
Chaetodon tricinctus
Chaetodon plebeius
Chaetodon bennetti
Chaetodon zanzibarensis
Chaetodon speculum
Chaetodon ocellatus
Chaetodon humeralis
Chaetodon striatus
Chaetodon capistratus
Chaetodon gardineri
Chaetodon selene
Chaetodon leucopleura
Chaetodon ocellicaudus
Chaetodon melannotus
Chaetodon semeion
Chaetodon xanthocephalus
Chaetodon ephippium
Chaetodon collare
Chaetodon flavirostris
Chaetodon adiergastos
Chaetodon auripes
Chaetodon lunula
Chaetodon fasciatus
Chaetodon rafflesii
Chaetodon semilarvatus
Chaetodon lineolatus
Chaetodon oxycephalus
Chaetodon falcula
Chaetodon ulietensis
Chaetodon dialeucos
Chaetodon mesoleucos
Chaetodon nigropunctatus
Chaetodon auriga
Chaetodon vagabundus
Chaetodon decussatus
Chaetodon vagabundus
Chaetodon pictus
To Supplementary Figure 1e
BS = 72-92
BS = 93-99
0.08 Substitutions per site

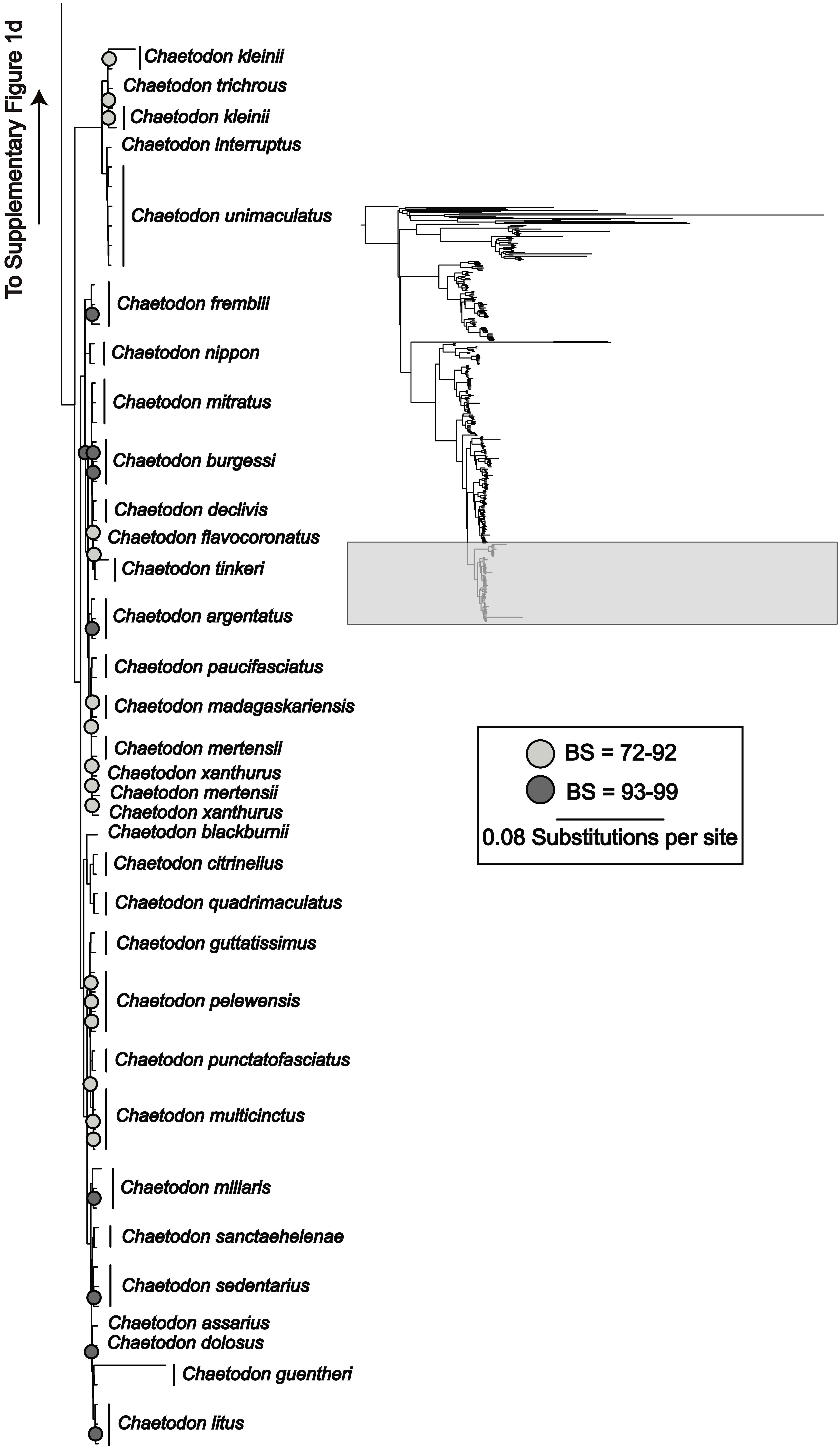

To Supplementary Figure 1d
Chaetodon kleinii
Chaetodon trichrous
Chaetodon kleinii
Chaetodon interruptus
Chaetodon unimaculatus
Chaetodon fremblii
Chaetodon nippon
Chaetodon mitratus
Chaetodon burgessi
Chaetodon declivis
Chaetodon flavocoronatus
Chaetodon tinkeri
Chaetodon argentatus
Chaetodon paucifasciatus
Chaetodon madagaskariensis
Chaetodon mertensii
Chaetodon xanthurus
Chaetodon mertensii
Chaetodon xanthurus
Chaetodon blackburnii
Chaetodon citrinellus
Chaetodon quadrimaculatus
Chaetodon guttatissimus
Chaetodon pelewensis
Chaetodon punctatofasciatus
Chaetodon multicinctus
Chaetodon miliaris
Chaetodon sanctaehelenae
Chaetodon sedentarius
Chaetodon assarius
Chaetodon dolosus
Chaetodon guentheri
Chaetodon litus
BS = 72-92
BS = 93-99
0.08 Substitutions per site

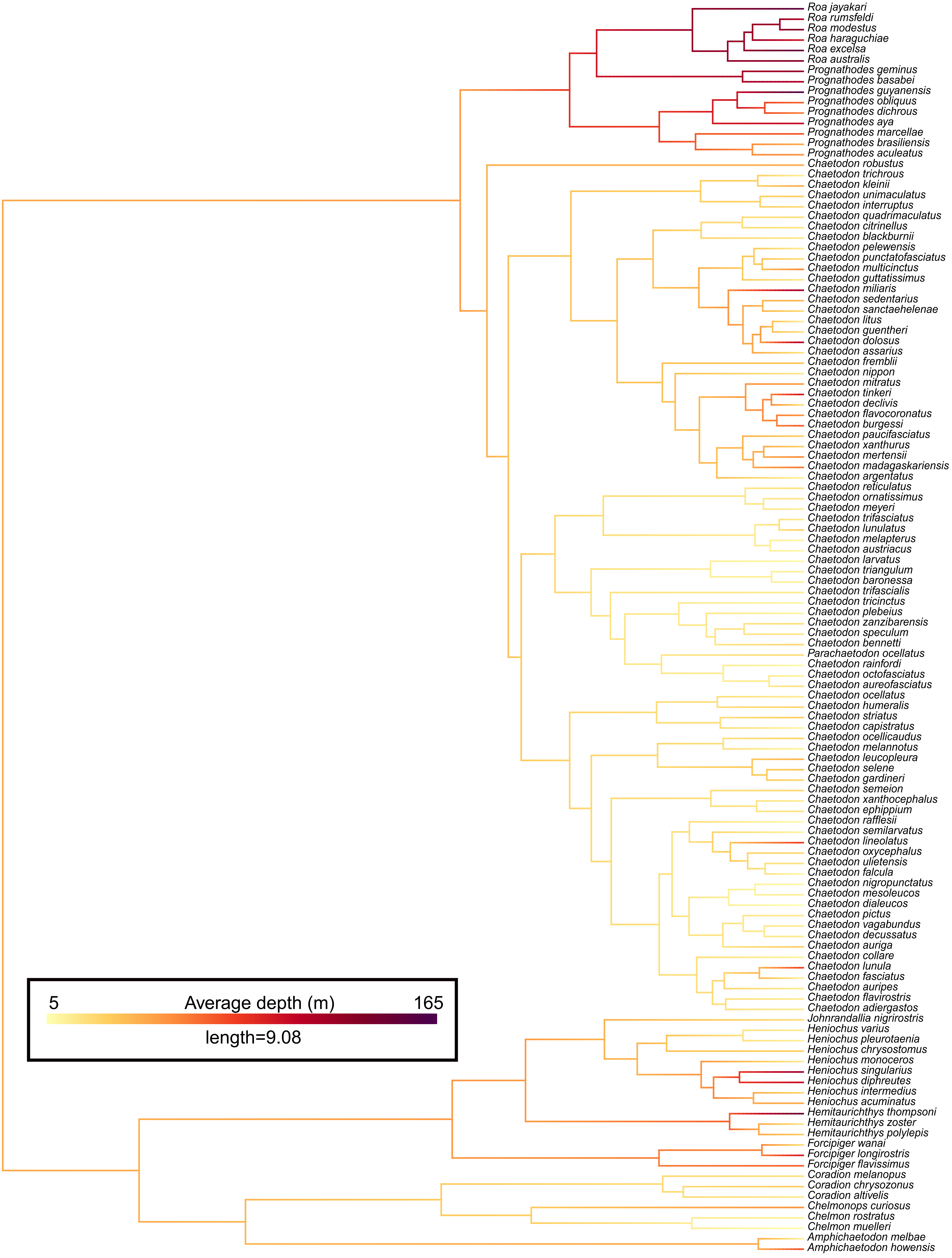
Roa jayakari
Roa rumsfeldi
Roa modestus
Roa haraguchiae
Roa excelsa
Roa australis
Prognathodes geminus
Prognathodes basabei
Prognathodes guyanensis
Prognathodes obliquus
Prognathodes dichrous
Prognathodes aya
Prognathodes marcellae
Prognathodes brasiliensis
Prognathodes aculeatus
Chaetodon robustus
Chaetodon trichrous
Chaetodon kleinii
Chaetodon unimaculatus
Chaetodon interruptus
Chaetodon quadrimaculatus
Chaetodon citrinellus
Chaetodon blackburnii
Chaetodon pelewensis
Chaetodon punctatofasciatus
Chaetodon multicinctus
Chaetodon guttatissimus
Chaetodon miliaris
Chaetodon sedentarius
Chaetodon sanctaehelenae
Chaetodon litus
Chaetodon guentheri
Chaetodon dolosus
Chaetodon assarius
Chaetodon fremblii
Chaetodon nippon
Chaetodon mitratus
Chaetodon tinkeri
Chaetodon declivis
Chaetodon flavocoronatus
Chaetodon burgessi
Chaetodon paucifasciatus
Chaetodon xanthurus
Chaetodon mertensii
Chaetodon madagaskariensis
Chaetodon argentatus
Chaetodon reticulatus
Chaetodon ornatissimus
Chaetodon meyeri
Chaetodon trifasciatus
Chaetodon lunulatus
Chaetodon melapterus
Chaetodon austriacus
Chaetodon larvatus
Chaetodon triangulum
Chaetodon baronessa
Chaetodon trifascialis
Chaetodon tricinctus
Chaetodon plebeius
Chaetodon zanzibarensis
Chaetodon speculum
Chaetodon bennetti
Parachaetodon ocellatus
Chaetodon rainfordi
Chaetodon octofasciatus
Chaetodon aureofasciatus
Chaetodon ocellatus
Chaetodon humeralis
Chaetodon striatus
Chaetodon capistratus
Chaetodon ocellicaudus
Chaetodon melannotus
Chaetodon leucopleura
Chaetodon selene
Chaetodon gardineri
Chaetodon semeion
Chaetodon xanthocephalus
Chaetodon ephippium
Chaetodon rafflesii
Chaetodon semilarvatus
Chaetodon lineolatus
Chaetodon oxycephalus
Chaetodon ulietensis
Chaetodon falcula
Chaetodon nigropunctatus
Chaetodon mesoleucos
Chaetodon dialeucos
Chaetodon pictus
Chaetodon vagabundus
Chaetodon decussatus
Chaetodon auriga
Chaetodon collare
Chaetodon lunula
Chaetodon fasciatus
Chaetodon auripes
Chaetodon flavirostris
Chaetodon adiergastos
Johnrandallia nigrirostris
Heniochus varius
Heniochus pleurotaenia
Heniochus chrysostomus
Heniochus monoceros
Heniochus singularius
Heniochus diphreutes
Heniochus intermedius
Heniochus acuminatus
Hemitaurichthys thompsoni
Hemitaurichthys zoster
Hemitaurichthys polylepis
Forcipiger wanai
Forcipiger longirostris
Forcipiger flavissimus
Coradion melanopus
Coradion chrysozonus
Coradion altivelis
Chelmonops curiosus
Chelmon rostratus
Chelmon muelleri
Amphichaetodon melbae
Amphichaetodon howensis
5
Average depth (m)
165
length=9.08

**Supplementary Table 1.** Sample information for *Chaetodontidae* taxa included in the phylogenomic analysis. For each sample, we report species identity, museum voucher number, collection locality, and the GenBank accession number.

| Sample No. | Species | Voucher No. | Collection Locality (Country/State/Parish) | GenBank Accession No. |
|---|---|---|---|---|
| 1. | *Amphichaetodon howensis* | M966 | Malabar, Lord Howe Island, Australia | |
| 2. | *Amphichaetodon howensis* | M967 | Malabar, Lord Howe Island, Australia | |
| 3. | *Amphichaetodon melbae* | FMNH107370 | San Felix Island, Chile | |
| 4. | *Chaetodon adiergastos* | CHD01 | Philippines | |
| 5. | *Chaetodon adiergastos* | SJL036 | Philippines | |
| 6. | *Chaetodon argentatus* | M1589 | Okinawa, Japan | |
| 7. | *Chaetodon argentatus* | M1590 | Okinawa, Japan | |
| 8. | *Chaetodon argentatus* | M1591 | Okinawa, Japan | |
| 9. | *Chaetodon assarius* | M1684 | Perth, Australia | |
| 10. | *Chaetodon aureofasciatus* | M1637 | Lizard Island, Australia | |
| 11. | *Chaetodon aureofasciatus* | M1656 | Rib Reef, Queensland, Australia | |
| 12. | *Chaetodon auriga* | CAZ01 | Saudi Arabia | |
| 13. | *Chaetodon auriga* | CAZ04 | Taiwan | |
| 14. | *Chaetodon auriga* | | GenBank sequence database (NCBI) | SAMN48779023 |
| 15. | *Chaetodon auripes* | CEU01 | Taiwan | |
| 16. | *Chaetodon austriacus* | CAI02 | Saudi Arabia | |
| 17. | *Chaetodon baronessa* | CBN01 | Philippines | |
| 18. | *Chaetodon baronessa* | SJL027 | Philippines | |
| 19. | *Chaetodon bennetti* | CNN01 | Socotra Island, Yemen | |
| 20. | *Chaetodon bennetti* | PW315 | Palau | |
| 21. | *Chaetodon bennetti* | PW316 | Palau | |
| 22. | *Chaetodon blackburnii* | PW4271 | Purchased from the pet trade | |
| 23. | *Chaetodon burgessi* | JRH80 | Koon Island, Watubelas Islands, Indonesia | |
| 24. | *Chaetodon burgessi* | JRH81 | Cenderawasih Bay, West Papua, Indonesia | |
| 25. | *Chaetodon burgessi* | PW4135 | Purchased from the pet trade | |
| 26. | *Chaetodon capistratus* | PW3429 | Curacao | |
| 27. | *Chaetodon capistratus* | PW3430 | Curacao | |
| 28. | *Chaetodon capistratus* | PW3467 | Curacao | |
| 29. | *Chaetodon capistratus* | PW4844 | Purchased from the pet trade | |
| 30. | *Chaetodon citrinellus* | FMNH119310 | Babelthuap Island, Palau | |
| 31. | *Chaetodon citrinellus* | FMNH120324 | Lillian's reef, Papua New Guinea | |

| Sample No. | Species | Voucher No. | Collection Locality (Country/State/Parish) | GenBank Accession No. |
|---|---|---|---|---|
| 32. | *Chaetodon collare* | CHCO01 | Somaliland, Somalia | |
| 33. | *Chaetodon collare* | CHCO02 | Maldives | |
| 34. | *Chaetodon collare* | PW3052 | Purchased from the pet trade | |
| 35. | *Chaetodon collare* | PW3054 | Purchased from the pet trade | |
| 36. | *Chaetodon declivis* | CDE01 | Marquesas, French Polynesia | |
| 37. | *Chaetodon declivis* | CDE02 | Marquesas, French Polynesia | |
| 38. | *Chaetodon decussatus* | M1546 | Rowley Shoals, Australia | |
| 39. | *Chaetodon decussatus* | M1547 | Rowley Shoals, Australia | |
| 40. | *Chaetodon decussatus* | PW4845 | Purchased from the pet trade | |
| 41. | *Chaetodon dialeucos* | CDA01 | Oman | |
| 42. | *Chaetodon dolosus* | KU5066 | Aliwal Shoal, South Africa | |
| 43. | *Chaetodon ephippium* | CEH01 | Taiwan | |
| 44. | *Chaetodon ephippium* | PW3015 | Purchased from the pet trade | |
| 45. | *Chaetodon ephippium* | PW3016 | Purchased from the pet trade | |
| 46. | *Chaetodon falcula* | CHL01 | Chagos Archipelago | |
| 47. | *Chaetodon fasciatus* | CFT01 | Saudi Arabia | |
| 48. | *Chaetodon fasciatus* | CFT06 | Somaliland, Somalia | |
| 49. | *Chaetodon flavirostris* | USNM424207 | French Polynesia | |
| 50. | *Chaetodon flavocoronatus* | MCE847 | Caroline Islands, Federated States of Micronesia | |
| 51. | *Chaetodon fremblii* | PW1517 | Hawaii, United States | |
| 52. | *Chaetodon fremblii* | PW1518 | Hawaii, United States | |
| 53. | *Chaetodon fremblii* | PW4303 | Kona, Hawaii, United States | |
| 54. | *Chaetodon gardineri* | PW4074 | Purchased from the pet trade | |
| 55. | *Chaetodon guentheri* | FMNH128374 | Philippines | |
| 56. | *Chaetodon guentheri* | CGN01 | Philippines | |
| 57. | *Chaetodon guttatissimus* | CGU14 | Chagos Archipelago | |
| 58. | *Chaetodon guttatissimus* | CGU15 | Maldives | |
| 59. | *Chaetodon humeralis* | PW1006 | Los Perlas, Panama | |
| 60. | *Chaetodon humeralis* | PW1007 | Los Perlas, Panama | |
| 61. | *Chaetodon humeralis* | CHU04 | Baja California, Mexico | |
| 62. | *Chaetodon interruptus* | PW4842E | Purchased from the pet trade | |
| 63. | *Chaetodon interruptus* | PW4842A | Purchased from the pet trade | |
| 64. | *Chaetodon kleinii* | CKL25 | Chagos Archipelago | |
| 65. | *Chaetodon kleinii* | PW1870 | Western outer reef , Palau | |
| 66. | *Chaetodon kleinii* | PW4831 | Purchased from the pet trade | |
| 67. | *Chaetodon larvatus* | CLA01 | Saudi Arabia | |
| 68. | *Chaetodon larvatus* | CLA07 | Somaliland, Somalia | |

| Sample No. | Species | Voucher No. | Collection Locality (Country/State/Parish) | GenBank Accession No. |
|---|---|---|---|---|
| 69. | *Chaetodon leucopleura* | CTL02 | Oman | |
| 70. | *Chaetodon leucopleura* | PW4078 | Purchased from the pet trade | |
| 71. | *Chaetodon lineolatus* | FMNH123553 | Palau | |
| 72. | *Chaetodon litus* | PW4846 | Anakena, Rapa Nui, Chile | |
| 73. | *Chaetodon litus* | PW4847 | Anakena, Rapa Nui, Chile | |
| 74. | *Chaetodon litus* | PW4848 | Anakena, Rapa Nui, Chile | |
| 75. | *Chaetodon lunula* | CHLU11 | Socotra Island, Yemen | |
| 76. | *Chaetodon lunulatus* | SJL061 | Philippines | |
| 77. | *Chaetodon lunulatus* | PW1896 | Palau | |
| 78. | *Chaetodon madagaskariensis* | CMD10 | Chagos Archipelago | |
| 79. | *Chaetodon madagaskariensis* | CMD11 | Maldives | |
| 80. | *Chaetodon melannotus* | CMN01 | Saudi Arabia | |
| 81. | *Chaetodon melannotus* | CMN08 | Maldives | |
| 82. | *Chaetodon melannotus* | PW4833 | Fiji | |
| 83. | *Chaetodon melapterus* | CMP05 | Somaliland, Somalia | |
| 84. | *Chaetodon mertensii* | PW3002 | Purchased from the pet trade | |
| 85. | *Chaetodon mertensii* | PW3003 | Purchased from the pet trade | |
| 86. | *Chaetodon mesoleucos* | CMS01 | Saudi Arabia | |
| 87. | *Chaetodon mesoleucos* | CMS10 | Somaliland, Somalia | |
| 88. | *Chaetodon meyeri* | FMNH118571 | Tobi Island, Palau | |
| 89. | *Chaetodon miliaris* | PW1521 | Hawaii, United States | |
| 90. | *Chaetodon miliaris* | PW1535 | Hawaii, United States | |
| 91. | *Chaetodon miliaris* | PW4861 | | |
| 92. | *Chaetodon mitratus* | CHI01 | Chagos Archipelago | |
| 93. | *Chaetodon mitratus* | CHI02 | Maldives | |
| 94. | *Chaetodon mitratus* | JRH82 | Barren Island, Andaman Islands, India | |
| 95. | *Chaetodon multicinctus* | PW1520 | Hawaii, United States | |
| 96. | *Chaetodon multicinctus* | PW4346 | South Point, Big Island, Hawaii, United States | |
| 97. | *Chaetodon multicinctus* | PW5531 | South Point, Big Island, Hawaii, United States | |
| 98. | *Chaetodon multicinctus* | PW5532 | South Point, Big Island, Hawaii, United States | |
| 99. | *Chaetodon nigropunctatus* | CNC01 | | |
| 100. | *Chaetodon nigropunctatus* | CNC02 | | |
| 101. | *Chaetodon nigropunctatus* | CNC03 | | |
| 102. | *Chaetodon nippon* | M1853 | Taipei, Taiwan | |
| 103. | *Chaetodon nippon* | M1854 | Taipei, Taiwan | |

| Sample No. | Species | Voucher No. | Collection Locality (Country/State/Parish) | GenBank Accession No. |
|---|---|---|---|---|
| 104. | *Chaetodon ocellatus* | COC01 | Bahamas | |
| 105. | *Chaetodon ocellicaudus* | FMNH128372 | Philippines | |
| 106. | *Chaetodon ocellicaudus* | CLL01 | Philippines | |
| 107. | *Chaetodon octofasciatus* | COT20 | Palau | |
| 108. | *Chaetodon octofasciatus* | COT24A | Philippines | |
| 109. | *Chaetodon octofasciatus* | COT24B | Philippines | |
| 110. | *Chaetodon octofasciatus* | PW3013 | Purchased from the pet trade | |
| 111. | *Chaetodon octofasciatus* | PW3014 | Purchased from the pet trade | |
| 112. | *Chaetodon ornatissimus* | PW5538 | South Point, Big Island, Hawaii, United States | |
| 113. | *Chaetodon ornatissimus* | PW5539 | South Point, Big Island, Hawaii, United States | |
| 114. | *Chaetodon oxycephalus* | COY01 | Maldives | |
| 115. | *Chaetodon oxycephalus* | FMNH119318 | Tabletop Reef, Kayangel, Palau | |
| 116. | *Chaetodon paucifasciatus* | CPF01 | Saudi Arabia | |
| 117. | *Chaetodon paucifasciatus* | CPF02 | Saudi Arabia | |
| 118. | *Chaetodon pelewensis* | PW3004 | Purchased from the pet trade | |
| 119. | *Chaetodon pelewensis* | PW3005 | Purchased from the pet trade | |
| 120. | *Chaetodon pelewensis* | PW4835A | South Pacific | |
| 121. | *Chaetodon pelewensis* | PW4835B | South Pacific | |
| 122. | *Chaetodon pictus* | CPS01 | Oman | |
| 123. | *Chaetodon pictus* | CPS02 | Somaliland, Somalia | |
| 124. | *Chaetodon plebeius* | CPB01 | Philippines | |
| 125. | *Chaetodon plebeius* | FMNH120344 | Lion Island, Papua, New Guinea | |
| 126. | *Chaetodon punctatofasciatus* | PW4843 | Purchased from the pet trade | |
| 127. | *Chaetodon punctatofasciatus* | SJL013 | Philippines | |
| 128. | *Chaetodon quadrimaculatus* | FNMH128429 | Whaler's Cove, South Shore, Kauai, Hawaii, United States | |
| 129. | *Chaetodon quadrimaculatus* | PW4068 | Kona, Hawaii, United States | |
| 130. | *Chaetodon rafflesii* | CRL01 | Philippines | |
| 131. | *Chaetodon rainfordi* | FMNH128538 | Purchased from the pet trade | |
| 132. | *Chaetodon reticulatus* | FMNH123545 | Seventy Islands, Koror, Palau | |
| 133. | *Chaetodon reticulatus* | PW2267 | Palau | |
| 134. | *Chaetodon robustus* | CRB01 | São Tomé Island, Gulf of Guinea | |
| 135. | *Chaetodon robustus* | CRB12 | Taiwan | |
| 136. | *Chaetodon robustus* | PW3965 | Purchased from the pet trade | |
| 137. | *Chaetodon sanctaehelenae* | CHSA01 | Ascension Island, South Atlantic Ocean | |

| Sample No. | Species | Voucher No. | Collection Locality (Country/State/Parish) | GenBank Accession No. |
|---|---|---|---|---|
| 138. | *Chaetodon sanctaehelenae* | CHSA02 | Ascension Island, South Atlantic Ocean | |
| 139. | *Chaetodon sedentarius* | PW1245 | Florida Keys, United States | |
| 140. | *Chaetodon sedentarius* | PW1246 | Florida Keys, United States | |
| 141. | *Chaetodon sedentarius* | PW1247 | Florida Keys, United States | |
| 142. | *Chaetodon selene* | CHS01 | Philippines | |
| 143. | *Chaetodon semeion* | FMNH118569 | Pulo Anna, Sonsorol, Palau | |
| 144. | *Chaetodon semilarvatus* | CSE01 | Saudi Arabia | |
| 145. | *Chaetodon semilarvatus* | CSE02 | Saudi Arabia | |
| 146. | *Chaetodon speculum* | CHP01 | Philippines | |
| 147. | *Chaetodon speculum* | PW319 | Palau | |
| 148. | *Chaetodon speculum* | PW320 | Palau | |
| 149. | *Chaetodon striatus* | CHR01 | Curacao | |
| 150. | *Chaetodon striatus* | PW3468 | Curacao | |
| 151. | *Chaetodon striatus* | PW4854 | Purchased from the pet trade | |
| 152. | *Chaetodon tinkeri* | PW1516 | Big Island, Hawaii, United States | |
| 153. | *Chaetodon tinkeri* | PW1545 | Hawaii, United States | |
| 154. | *Chaetodon triangulum* | CTG01 | Maldives | |
| 155. | *Chaetodon trichrous* | PW4270 | Purchased from the pet trade | |
| 156. | *Chaetodon tricinctus* | M965 | Lagoon, Lord Howe Island, Australia | |
| 157. | *Chaetodon trifascialis* | CTR05 | Chagos Archipelago | |
| 158. | *Chaetodon trifascialis* | | GenBank sequence database (NCBI) | SAMN41253346 |
| 159. | *Chaetodon trifasciatus* | | GenBank sequence database (NCBI) | SAMN46283988 |
| 160. | *Chaetodon trifasciatus* | FP52 | Moorea, French Polynesia | |
| 161. | *Chaetodon trifasciatus* | CFT01 | Maldives | |
| 162. | *Chaetodon trifasciatus* | CFT47 | Socotra Island, Yemen | |
| 163. | *Chaetodon ulietensis* | FMNH119327 | Babelthuap Island, Palau | |
| 164. | *Chaetodon ulietensis* | PW4837 | Fiji | |
| 165. | *Chaetodon unimaculatus* | FP62 | Moorea, French Polynesia | |
| 166. | *Chaetodon unimaculatus* | CUN01 | Philippines | |
| 167. | *Chaetodon unimaculatus* | PW1522 | Hawaii, United States | |
| 168. | *Chaetodon unimaculatus* | PW1523 | Hawaii, United States | |
| 169. | *Chaetodon unimaculatus* | PW4836 | Fiji | |
| 170. | *Chaetodon vagabundus* | FMNH120353 | Lion Island, Papua New Guinea | |
| 171. | *Chaetodon vagabundus* | PW4860 | Purchased from the pet trade | |

| Sample No. | Species | Voucher No. | Collection Locality (Country/State/Parish) | GenBank Accession No. |
|---|---|---|---|---|
| 172. | *Chaetodon xanthocephalus* | CXN01 | Chagos Archipelago | |
| 173. | *Chaetodon xanthurus* | CXH01 | Philippines | |
| 174. | *Chaetodon xanthurus* | PW4859 | Purchased from the pet trade | |
| 175. | *Chaetodon xanthurus* | SJL044 | Philippines | |
| 176. | *Chaetodon zanzibarensis* | CZA01 | Chagos Archipelago | |
| 177. | *Chelmon muelleri* | JRH83 | Queensland, Australia | |
| 178. | *Chelmon muelleri* | JRH84 | Queensland, Australia | |
| 179. | *Chelmon muelleri* | M1611 | Bay Rock, Queensland, Australia | |
| 180. | *Chelmon muelleri* | M1612 | Bay Rock, Queensland, Australia | |
| 181. | *Chelmon rostratus* | JRH85 | Mesempta, Southeast Misool, Raja Ampat, West Papua, Indonesia | |
| 182. | *Chelmon rostratus* | PW3099 | Purchased from the pet trade | |
| 183. | *Chelmon rostratus* | PW3100 | Purchased from the pet trade | |
| 184. | *Chelmon rostratus* | PW3101 | Purchased from the pet trade | |
| 185. | *Chelmon rostratus* | PW4838 | Philippines | |
| 186. | *Chelmonops curiosus* | M1682 | Perth, Australia | |
| 187. | *Chelmonops curiosus* | M1683 | Perth, Australia | |
| 188. | *Coradion altivelis* | PW4073 | Purchased from the pet trade | |
| 189. | *Coradion chrysozonus* | COH01 | Philippines | |
| 190. | *Coradion chrysozonus* | SJL029 | Philippines | |
| 191. | *Coradion melanopus* | FMNH124029 | Purchased from the pet trade | |
| 192. | *Coradion melanopus* | COM01 | Philippines | |
| 193. | *Forcipiger flavissimus* | FFL01 | Maldives | |
| 194. | *Forcipiger flavissimus* | FFL02 | Socotra Island, Yemen | |
| 195. | *Forcipiger flavissimus* | PW5534 | South Point, Big Island, Hawaii, United States | |
| 196. | *Forcipiger flavissimus* | PW5535 | South Point, Big Island, Hawaii, United States | |
| 197. | *Forcipiger longirostris* | | GenBank sequence database (NCBI) | SAMN09765373 |
| 198. | *Forcipiger longirostris* | FMNH120361 | Lillian's Reef, Papua New Guinea | |
| 199. | *Forcipiger longirostris* | SJL018 | Philippines | |
| 200. | *Forcipiger wanai* | JRH86 | Kali Lemon, Cenderawasih Bay, West Papua, Indonesia | |
| 201. | *Forcipiger wanai* | JRH87 | Kali Lemon, Cenderawasih Bay, West Papua, Indonesia | |
| 202. | *Forcipiger wanai* | JRH88 | Kali Lemon, Cenderawasih Bay, West Papua, Indonesia | |

| Sample No. | Species | Voucher No. | Collection Locality (Country/State/Parish) | GenBank Accession No. |
|---|---|---|---|---|
| 203. | *Hemitaurichthys polylepis* | PW1596 | Philippines | |
| 204. | *Hemitaurichthys polylepis* | PW2024 | Palau | |
| 205. | *Hemitaurichthys polylepis* | PW3095 | Purchased from the pet trade | |
| 206. | *Hemitaurichthys polylepis* | PW4629 | South Point, Big Island, Hawaii, United States | |
| 207. | *Hemitaurichthys thompsoni* | PW4631 | South Point, Big Island, Hawaii, United States | |
| 208. | *Hemitaurichthys thompsoni* | PW4652 | South Point, Big Island, Hawaii, United States | |
| 209. | *Hemitaurichthys thompsoni* | PW5536 | South Point, Big Island, Hawaii, United States | |
| 210. | *Hemitaurichthys thompsoni* | PW5537 | South Point, Big Island, Hawaii, United States | |
| 211. | *Hemitaurichthys zoster* | HZO01 | Chagos Archipelago | |
| 212. | *Hemitaurichthys zoster* | PW4841 | Purchased from the pet trade | |
| 213. | *Hemitaurichthys zoster* | PW4855 | Purchased from the pet trade | |
| 214. | *Heniochus acuminatus* | HAC05 | Philippines | |
| 215. | *Heniochus acuminatus* | PW4514 | Purchased from the pet trade | |
| 216. | *Heniochus acuminatus* | PW4560 | Purchased from the pet trade | |
| 217. | *Heniochus chrysostomus* | PW2923 | Purchased from the pet trade | |
| 218. | *Heniochus chrysostomus* | PW2924 | Purchased from the pet trade | |
| 219. | *Heniochus chrysostomus* | SJL020 | Philippines | |
| 220. | *Heniochus diphreutes* | HDP01 | Maldives | |
| 221. | *Heniochus diphreutes* | FMNH124021 | Bagalayag Point, Occidental Mindoro, Philippines | |
| 222. | *Heniochus intermedius* | HIN01 | Saudi Arabia | |
| 223. | *Heniochus intermedius* | HIN02 | Saudi Arabia | |
| 224. | *Heniochus monoceros* | PW3173 | Purchased from the pet trade | |
| 225. | *Heniochus monoceros* | FMNH128371 | Dimipac Island, Busuanga, Palawan, Philippines | |
| 226. | *Heniochus pleurotaenia* | HPR01 | Maldives | |
| 227. | *Heniochus singularius* | HSI01 | Philippines | |
| 228. | *Heniochus varius* | HVA01 | Philippines | |
| 229. | *Heniochus varius* | PW2921 | Purchased from the pet trade | |
| 230. | *Heniochus varius* | PW2922 | Purchased from the pet trade | |
| 231. | *Johnrandallia nigrirostris* | JNI01 | Baja California, Mexico | |
| 232. | *Johnrandallia nigrirostris* | JNI06 | Baja California, Mexico | |
| 233. | *Johnrandallia nigrirostris* | PW4857 | Purchased from the pet trade | |
| 234. | *Johnrandallia nigrirostris* | PW933 | Saboga Island, Pearl Islands, Panama | |

| Sample No. | Species | Voucher No. | Collection Locality (Country/State/Parish) | GenBank Accession No. |
|---|---|---|---|---|
| 235. | *Parachaetodon ocellatus* | PW1643 | Purchased from the pet trade | |
| 236. | *Prognathodes aculeatus* | PW648 | Purchased from the pet trade | |
| 237. | *Prognathodes aculeatus* | PW3487 | Curacao | |
| 238. | *Prognathodes aculeatus* | CAC04 | Curacao | |
| 239. | *Prognathodes aya* | PW1042 | Florida Keys, United States | |
| 240. | *Prognathodes aya* | PW1043 | Florida Keys, United States | |
| 241. | *Prognathodes aya* | PW1044 | Florida Keys, United States | |
| 242. | *Prognathodes aya* | PW1045 | Florida Keys, United States | |
| 243. | *Prognathodes basabei* | MCE2742 | Palau Islands | |
| 244. | *Prognathodes brasiliensis* | PW3997 | Purchased from the pet trade | |
| 245. | *Prognathodes dichrous* | PDH01 | Ascension Island, South Atlantic Ocean | |
| 246. | *Prognathodes dichrous* | PDH02 | Ascension Island, South Atlantic Ocean | |
| 247. | *Prognathodes geminus* | MCE878 | Northwestern Hawaiian Islands | |
| 248. | *Prognathodes guyanensis* | USNM414870 | Curacao | |
| 249. | *Prognathodes marcellae* | PMR01 | São Tomé Island, Gulf of Guinea | |
| 250. | *Prognathodes obliquus* | POQ01 | São Tomé Island, Gulf of Guinea | |
| 251. | *Prognathodes obliquus* | POQ02 | Saint Peter and Saint Paul Archipelago, Brazil | |
| 252. | *Roa australis* | M2598 | Leveque, Northwestern Australia | |
| 253. | *Roa excelsa* | Roa sp. 02 | Philippines | |
| 254. | *Roa haraguchiae* | KAUM35795 | East China Sea, Japan | |
| 255. | *Roa jayakari* | JDD98 | Red Sea | |
| 256. | *Roa modesta* | CMO01 | Philippines | |
| 257. | *Roa rumsfeldi* | Roa sp. 01 | Philippines | |

**Supplementary Table 2.** Feeding ecology and intramandibular joint (IMJ) state for *Chaetodontidae* species included in this study. For each species, we report presence or absence of the IMJ, diet mode (non-coral feeder, facultative corallivore, or obligate corallivore), a summary of diet composition, and the literature source(s) supporting these classifications.

| Sample No. | Species | IMJ | Diet Mode | Diet Composition | Reference |
|---|---|---|---|---|---|
| 1. | *Amphichaetodon howensis* | Absent | Non-coral feeder | Non-coral benthic invertivores | Hodge et al. (2021) |
| 2. | *Amphichaetodon melbae* | Absent | Non-coral feeder | Non-coral benthic invertivores | Hodge et al. (2021) |
| 3. | *Chaetodon adiergastos* | Present | Facultative corallivore | Corals, crabs, worms, and other invertebrates | Cole et al. (2008); Nagelkerken et al. (2009); Froese and Pauly (2008) |
| 4. | *Chaetodon argentatus* | Present | Facultative corallivore | Scleractinians (hard corals), Alcyonarians (soft corals), filamentous algae, polychaetes and Hydroids | Cole et al. (2008); Sano (1989) |
| 5. | *Chaetodon assarius* | Present | Non-coral feeder | Filamentous algae and zooplankton | Froese and Pauly (2008) |
| 6. | *Chaetodon aureofasciatus* | Present | Obligate corallivore | Scleractinians (hard corals) almost exclusively | Cole et al. (2008); Pratchett (2005); Pratchett (2007) |
| 7. | *Chaetodon auriga* | Present | Facultative corallivore | Benthic invertebrates, coral polyps (Scleractinians), polychaetes and sea anemones | Cole et al. (2008); Sano (1989); Nagelkerken et al. (2009); Harmelin-Vivien & Bouchon-Navaro (1983) |
| 8. | *Chaetodon auripes* | Present | Facultative corallivore | Scleractinians (hard corals), polychaetes, other benthic invertebrates, filamentous algae | Cole et al. (2008); Sano (1989) |
| 9. | *Chaetodon austriacus* | Present | Obligate corallivore | Scleractinians (hard corals) exclusively; particularly *Acropora spp.*; strongly associated with *Acropora* colonies | Cole et al. (2008); Bouchon-Navaro & Bouchon (1989); Alwany et al. (2003) |
| 10. | *Chaetodon baronessa* | Present | Obligate corallivore | Scleractinians (hard corals) exclusively, highly selective for *Acropora hyacinthus* | Cole et al. (2008); Nagelkerken et al. (2009); Sano (1989); Pratchett (2005); Pratchett (2007) |
| 11. | *Chaetodon bennetti* | Present | Obligate corallivore | Scleractinians (hard corals) exclusively, feeds primarily on *Acropora* and *Pocillopora* | Cole et al. (2008); Nagelkerken et al. (2009); Sano (1989); Harmelin-Vivien & Bouchon-Navaro (1983) |

| Sample No. | Species | IMJ | Diet Mode | Diet Composition | Reference |
|---|---|---|---|---|---|
| 12. | *Chaetodon blackburnii* | Present | Facultative corallivore | Alcyonarians (soft corals) and other non-coral invertebrates | Cole et al. (2008); Harmelin-Vivien (1989) |
| 13. | *Chaetodon burgessi* | Present | Non-coral feeder | | Michael (2004); Pratchett et al. (2013) |
| 14. | *Chaetodon capistratus* | Present | Facultative corallivore | 75% of the diet is derived from Scleractinians (hard corals), also polychaetes and small invertebrates | Cole et al. (2008); Randall (1967) |
| 15. | *Chaetodon citrinellus* | Present | Facultative corallivore | Scleractinians (hard corals), filamentous algae, polychaetes, hydroids | Cole et al. (2008); Sano (1989); Harmelin-Vivien & Bouchon-Navaro (1983); Pratchett (2005) |
| 16. | *Chaetodon collare* | Present | Facultative corallivore | Coral polyps, worms, crustaceans and algae | Cole et al. (2008); Froese and Pauly (2008) |
| 17. | *Chaetodon declivis* | Present | Non-coral feeder | | Michael (2004); Pratchett et al. (2013) |
| 18. | *Chaetodon decussatus* | Present | Facultative corallivore | Coral polyp and filamentous algae | Cole et al. (2008) |
| 19. | *Chaetodon dialeucos* | Present | Non-coral feeder | | Michael (2004); Pratchett et al. (2013) |
| 20. | *Chaetodon dolosus* | Present | Non-coral feeder | | Michael (2004); Pratchett et al. (2013) |
| 21. | *Chaetodon ephippium* | Present | Facultative corallivore | Filamentous algae, Scleractinians (hard colars), sponges, polychaetes, Ascidians (Tunicata) | Cole et al. (2008); Sano (1989); Nagelkerken et al. (2009); Harmelin-Vivien & Bouchon-Navaro (1983) |
| 22. | *Chaetodon falcula* | Present | Facultative corallivore | Coral polyps and benthic invertebrates | Cole et al. (2008); Froese and Pauly (2008); McClanahan et al. (2005) |
| 23. | *Chaetodon fasciatus* | Present | Facultative corallivore | Coral polyps and benthic invertebrates | Cole et al. (2008) |
| 24. | *Chaetodon flavirostris* | Present | Facultative corallivore | Algae, coral polyp, and small benthic invertebrates | Cole et al. (2008); Froese and Pauly (2008) |
| 25. | *Chaetodon flavocoronatus* | Present | Non-coral feeder | Small benthic invertebrates, zoantharians, polychaete worms, gorgonians and tunicates, | Florida Museum (2025): https://www.floridamuseum.ufl.edu/discover-fish/species- |

| Sample No. | Species | IMJ | Diet Mode | Diet Composition | Reference |
|---|---|---|---|---|---|
| | | | | | profiles/foureye-butterflyfish/ |
| 26. | *Chaetodon fremblii* | Present | Non-coral feeder | Preys on wide variety of benthic organisms | Hobson (1974) |
| 27. | *Chaetodon gardineri* | Present | Facultative corallivore | | Cole et al. (2008); Pratchett et al. (2013) |
| 28. | *Chaetodon guentheri* | Present | Non-coral feeder | Zooplankton; selective zooplanktivore | Hodge et al. (2021) |
| 29. | *Chaetodon guttatissimus* | Present | Facultative corallivore | Polychaetes, coral polyps, and algae | Cole et al. (2008); Harmelin-Vivien (1989); Froese and Pauly (2008) |
| 30. | *Chaetodon humeralis* | Present | Non-coral feeder | Algae and benthic invertebrates | Froese and Pauly (2008) |
| 31. | *Chaetodon interruptus* | Present | Facultative corallivore | | Cole et al. (2008); Pratchett et al. (2013) |
| 32. | *Chaetodon kleinii* | Present | Facultative corallivore | Filamentous algae, Scleractinians (hard corals), Alcyonarians (soft corals), sponges, Hydroids, polychaetes | Cole et al. (2008); Nagelkerken et al. (2009); Sano (1989); Pratchett (2005) |
| 33. | *Chaetodon larvatus* | Present | Obligate corallivore | Scleractinians (hard corals) almost exclusively | Cole et al. (2008); Zekeria et al. (2002); |
| 34. | *Chaetodon leucopleura* | Present | Facultative corallivore | Coral polyp and other benthic invertebrates | Cole et al. (2008); |
| 35. | *Chaetodon lineolatus* | Present | Non-coral feeder | Non-coral macro-invertebrates; often feeds on small colonial zoanthids including *Zoanthus spp*. | Sano (1989); Pratchett (2005) |
| 36. | *Chaetodon litus* | Present | Non-coral feeder | Tube worms, sea anemones, and crustaceans | Florida Museum (2025) |
| 37. | *Chaetodon lunula* | Present | Facultative corallivore | Scleractinians (hard corals), polychaetes, and benthic invertebrates | Cole et al. (2008); Nagelkerken et al. (2009); Harmelin-Vivien & Bouchon-Navaro (1983); Pratchett (2005) |
| 38. | *Chaetodon lunulatus* | Present | Obligate corallivore | Scleractinians (hard corals) almost exclusively; mainly *Acropora* and *Pocillopora* | Cole et al. (2008); Nagelkerken et al. (2009); Pratchett (2005) |
| 39. | *Chaetodon madagaskariensis* | Present | Facultative corallivore | Scleractinians (hard corals), Alcyonarians (soft corals), filamentous algae, polychaetes | Harmelin-Vivien (1989) |

| Sample No. | Species | IMJ | Diet Mode | Diet Composition | Reference |
|---|---|---|---|---|---|
| 40. | *Chaetodon melannotus* | Present | Obligate corallivore | Alcyonarians/ Octocorallia (soft corals) almost exclusively | Cole et al. (2008); Bouchon-Navaro & Bouchon (1989) |
| 41. | *Chaetodon melapterus* | Present | Obligate corallivore | Alcyonarians (soft corals) almost exclusively | Cole et al. (2008); Froese and Pauly (2008) |
| 42. | *Chaetodon mertensii* | Present | Facultative corallivore | Coral, algae and small benthic invertebrates | Cole et al. (2008); Froese and Pauly (2008) |
| 43. | *Chaetodon mesoleucos* | Present | Facultative corallivore | Coral polyps, polychaetes, nematode | Cole et al. (2008); Zekeria et al. (2002) |
| 44. | *Chaetodon meyeri* | Present | Obligate corallivore | Scleractinians (hard corals) almost exclusively | Cole et al. (2008); Sano (1989) |
| 45. | *Chaetodon miliaris* | Present | Non-coral feeder | Zooplankton; selective zooplanktivore | Hodge et al. (2021); Hobson (1974) |
| 46. | *Chaetodon mitratus* | Present | Non-coral feeder | | Pratchett et al. (2013) |
| 47. | *Chaetodon multicinctus* | Present | Obligate corallivore | Scleractinians (hard corals) almost exclusively; highly selective for *Pocillipora meandrina* | Cole et al. (2008); Tricas (1989); Aeby (2002) |
| 48. | *Chaetodon nigropunctatus* | Present | Facultative corallivore | Partially coral polyps and benthic invertebrates and algae | Cole et al. (2008); Froese and Pauly (2008) |
| 49. | *Chaetodon nippon* | Present | Facultative corallivore | Polychaetes, filamentous algae, sponges, sea anemones, hydroids and some corals | Cole et al. (2008); Sano (1989) |
| 50. | *Chaetodon ocellatus* | Present | Facultative corallivore | | Cole et al. (2008); Pratchett et al. (2013) |
| 51. | *Chaetodon ocellicaudus* | Present | Obligate corallivore | Alcyonarians (soft corals) almost exclusively | Cole et al. (2008); Froese and Pauly (2008) |
| 52. | *Chaetodon octofasciatus* | Present | Obligate corallivore | Exclusively on coralm polyps; Scleractinians (hard corals) | Cole et al. (2008); Froese and Pauly (2008) |
| 53. | *Chaetodon ornatissimus* | Present | Obligate corallivore | Scleractinians (hard corals) almost exclusively | Cole et al. (2008); Nagelkerken et al. (2009); Sano (1989); Harmelin-Vivien & Bouchon-Navaro (1983) |
| 54. | *Chaetodon oxycephalus* | Present | Facultative corallivore | Coral polyps and anemones | Cole et al. (2008); Froese and Pauly (2008) |
| 55. | *Chaetodon paucifasciatus* | Present | Facultative corallivore | Scleractinians (hard corals) and other benthic invertebrates and polychaetes; | Cole et al. (2008); Bouchon-Navaro & Bouchon (1989) |

| Sample No. | Species | IMJ | Diet Mode | Diet Composition | Reference |
|---|---|---|---|---|---|
| | | | | strong association with Acropora among Red Sea species | |
| 56. | *Chaetodon pelewensis* | Present | Obligate corallivore | Scleractinians (hard corals), Alcyonarians (soft corals) | Harmelin-Vivien & Bouchon-Navaro (1983); Harmelin-Vivien (1989) |
| 57. | *Chaetodon pictus* | Present | Facultative corallivore | Tube worms, sea anemones, corals, and occasionally crustaceans | Florida Museum (2025): [https://www.floridamuseum.ufl.edu/discover-fish/species-profiles/banded-butterflyfish/](https://www.floridamuseum.ufl.edu/discover-fish/species-profiles/banded-butterflyfish/) |
| 58. | *Chaetodon plebeius* | Present | Obligate corallivore | Scleractinians (hard corals) almost exclusively; predominantly *Acropora* and *Pocillopora* | Kuiter (2002); Cole et al. (2008) Sano (1989); Pratchett (2005) |
| 59. | *Chaetodon punctatofasciatus* | Present | Obligate corallivore | Scleractinians (hard corals) almost exclusively; feeds primarily on *Acropora, Montipora spp*., *Pocillopora damicornis* and *Pachyseris speciosa*. | Cole et al. (2008); Nagelkerken et al. (2009) |
| 60. | *Chaetodon quadrimaculatus* | Present | Facultative corallivore | Scleractinians (hard corals), Alcyonarians (soft corals), filamentous algae and hydroids | Cole et al. (2008); Vivien & Bouchon-Navaro (1983) |
| 61. | *Chaetodon rafflesii* | Present | Facultative corallivore | Alcyonarians (soft corals); Scleractinians (hard corals), and other non-coral macro-invertebrates; in regards to hard corals, it particularly feed on one specific *Acropora tenuis* | Cole et al. (2008); Pratchett (2005); Sano (1989) |
| 62. | *Chaetodon rainfordi* | Present | Obligate corallivore | Scleractinians (hard corals) almost exclusively | Cole et al. (2008); Pratchett (2005); Pratchett (2007) |
| 63. | *Chaetodon reticulatus* | Present | Obligate corallivore | Scleractinians (hard corals) almost exclusively | Cole et al. (2008); Vivien & Bouchon-Navaro (1983) |
| 64. | *Chaetodon robustus* | Present | Non-coral feeder | | Pratchett et al. (2013) |
| 65. | *Chaetodon sanctaehelenae* | Present | Non-coral feeder | | Pratchett et al. (2013) |
| 66. | *Chaetodon sedentarius* | Present | Non-coral feeder | Polychaetes, small crustaceans, amphipods, hydroids | Randall (1967) |
| 67. | *Chaetodon selene* | Present | Non-coral feeder | Small benthic invertebrates | Froese and Pauly (2008) |
| 68. | *Chaetodon semeion* | Present | Non-coral feeder | Non-coral invertebrates; they prefer colonial ascidians | Nagelkerken et al. (2009); Sano (1989); Pratchett (2005) |

| Sample No. | Species | IMJ | Diet Mode | Diet Composition | Reference |
|---|---|---|---|---|---|
| 69. | *Chaetodon semilarvatus* | Present | Obligate corallivore | Scleractinians (hard corals) almost exclusively | Cole et al. (2008); Zekeria et al. (2002) |
| 70. | *Chaetodon speculum* | Present | Facultative corallivore | Primarily consumes Scleractinians (hard corals), specifically prefers *Hydonophora exesa*, but also feed on annelid worms, crustaceans and algae. | Cole et al. (2008); Pratchett (2005); Pratchett (2007); Nagelkerken et al. (2009); |
| 71. | *Chaetodon striatus* | Present | Facultative corallivore | Primarily polychaetes but also Anthozoans, including *Bunodactis stelloides* (sea anemonies) and *Zoanthus sp.* (soft corals) | Randall (1967) |
| 72. | *Chaetodon tinkeri* | Present | Non-coral feeder | Planktonic and small benthic invertabrates | Froese and Pauly (2008) |
| 73. | *Chaetodon triangulum* | Present | Obligate corallivore | Scleractinians (hard corals) almost exclusively | Cole et al. (2008); Froese and Pauly (2008) |
| 74. | *Chaetodon trichrous* | Present | Facultative corallivore | | Cole et al. (2008) |
| 75. | *Chaetodon tricinctus* | Present | Facultative corallivore | Corals and other non-coral invertebrates | Cole et al. (2008); Kuiter (1996) |
| 76. | *Chaetodon trifascialis* | Present | Obligate corallivore | Scleractinians (hard corals) almost exclusively; particularly *Acropora* | Cole et al. (2008); Sano (1989); Bouchon-Navaro & Bouchon (1989); Harmelin-Vivien & Bouchon-Navaro (1981); Alwany et al. (2003) |
| 77. | *Chaetodon trifasciatus* | Present | Obligate corallivore | Scleractinians (hard corals) almost exclusively | Cole et al. (2008); Sano (1989); Harmelin-Vivien & Bouchon-Navaro (1981) |
| 78. | *Chaetodon ulietensis* | Present | Facultative corallivore | Alcyonarians (soft corals), Scleractinians (hard corals), filamentous algae, other non-coral intertebrates | Cole et al. (2008); Harmelin-Vivien & Bouchon-Navaro (1981); Pratchett (2005) |
| 79. | *Chaetodon unimaculatus* | Present | Facultative corallivore | Alcyonarians (soft corals), Scleractinians (hard corals), crustacean fragments, filamentous algae | Sano (1989); Harmelin-Vivien & Bouchon-Navaro (1981) |
| 80. | *Chaetodon vagabundus* | Present | Facultative corallivore | Primarily filamentous algae, cephalopods, and polychaetes but also consume Scleractinians (hard corals) and some Alcyonarians (soft corals) | Cole et al. (2008); Harmelin-Vivien & Bouchon-Navaro (1981); Sano (1989); Nagelkerken et al. (2009) |

| Sample No. | Species | IMJ | Diet Mode | Diet Composition | Reference |
|---|---|---|---|---|---|
| 81. | *Chaetodon xanthocephalus* | Present | Facultative corallivore | Ascidians, sponges, Scleractinians (hard corals), filamentous algae, polychaetes | Cole et al. (2008); Harmelin-Vivien (1989) |
| 82. | *Chaetodon xanthurus* | Present | Facultative corallivore | Coral polyps, small bentic invertebrates and algae | Cole et al. (2008); Froese and Pauly (2008) |
| 83. | *Chaetodon zanzibarensis* | Present | Obligate corallivore | Scleractinians (hard corals) almost exclusively; particularly hard corals of the genus *Porites* | Cole et al. (2008) |
| 84. | *Chelmon muelleri* | Absent | Non-coral feeder | Small bentic non-coral invertebrates | Froese and Pauly (2008) |
| 85. | *Chelmon rostratus* | Absent | Non-coral feeder | Small bentic non-coral invertebrates | Michael (2004); Pratchett et al. (2013) |
| 86. | *Chelmonops curiosus* | Absent | Non-coral feeder | Worms and crustaceans | Froese and Pauly (2008) |
| 87. | *Coradion altivelis* | Absent | Non-coral feeder | Variety of invertebrates, in particular small amphipods that are often abundant on sponges | Kuiter (2002) |
| 88. | *Coradion chrysozonus* | Absent | Non-coral feeder | Variety of invertebrates, in particular small amphipods that are often abundant on sponges | Nagelkerken et al. (2009); Kuiter (2002) |
| 89. | *Coradion melanopus* | Absent | Non-coral feeder | Variety of invertebrates, in particular small amphipods that are often abundant on sponges | Kuiter (2002) |
| 90. | *Forcipiger flavissimus* | Absent | Facultative corallivore | Primarily polychaetes but also sponges and hydroids | Harmelin-Vivien & Bouchon-Navaro (1981); Nagelkerken et al. (2009) |
| 91. | *Forcipiger longirostris* | Absent | Non-coral feeder | Small benthic animals; mainly decapod shrimp | Hobson (1974) |
| 92. | *Forcipiger wanai* | Absent | Non-coral feeder | Small benthic animals | |
| 93. | *Hemitaurichthys polylepis* | Absent | Non-coral feeder | Zooplankton; selective zooplanktivore | Hodge et al. (2021); Kuiter (2002) |
| 94. | *Hemitaurichthys thompsoni* | Absent | Non-coral feeder | Zooplankton; selective zooplanktivore | Hodge et al. (2021); Kuiter (2002) |
| 95. | *Hemitaurichthys zoster* | Absent | Non-coral feeder | Zooplankton; selective zooplanktivore | Kuiter (2002) |
| 96. | *Heniochus acuminatus* | Absent | Non-coral feeder | Plankton | Froese and Pauly (2008) |
| 97. | *Heniochus chrysostomus* | Absent | Facultative corallivore | Alcyonarians (soft corals), and polychaetes | Cole et al. (2008); Froese and Pauly (2008); Bouchon-Navaro (1986) |

| Sample No. | Species | IMJ | Diet Mode | Diet Composition | Reference |
|---|---|---|---|---|---|
| 98. | *Heniochus diphreutes* | Absent | Non-coral feeder | Zooplankton; selective zooplanktivore | Hodge et al. (2021) |
| 99. | *Heniochus intermedius* | Absent | Facultative corallivore | Polychaetes, Scleractinians (hard corals), larvaceans, crabs and amphipods | Cole et al. (2008); Zekeria et al. (2002) |
| 100. | *Heniochus monoceros* | Absent | Non-coral feeder | Small benthic invertebrates | Froese and Pauly (2008) |
| 101. | *Heniochus pleurotaenia* | Absent | Non-coral feeder | Zooplankton and small benthic invertebrates | Michael (2004) |
| 102. | *Heniochus singularius* | Absent | Facultative corallivore | Coral polyps, benthic invertebrates and algae | Australian Museum (2019): https://australian.museum/learn/animals/fishes/singular-bannerfish-heniochus-singularius/ |
| 103. | *Heniochus varius* | Absent | Facultative corallivore | Coral polyps and other invertebrates | Cole et al. (2008); Froese and Pauly (2008) |
| 104. | *Johnrandallia nigrirostris* | Absent | Non-coral feeder | Small invertebrates and algae | Kuiter (2002) |
| 105. | *Parachaetodon ocellatus* | Present | Non-coral feeder | Small invertebrates, usually picked from narrow crevices or rock-surfaces with tubeworms | Kuiter (2002) |
| 106. | *Prognathodes aculeatus* | Absent | Non-coral feeder | Small invertebrates, usually picked from narrow crevices or rock-surfaces with tubeworms | Kuiter (2002) |
| 107. | *Prognathodes aya* | Absent | Non-coral feeder | Small invertebrates, usually picked from narrow crevices or rock-surfaces with tubeworms | Kuiter (2002) |
| 108. | *Prognathodes basabei* | Absent | Non-coral feeder | Small invertebrates, usually picked from narrow crevices or rock-surfaces with tubeworms | Kuiter (2002) |
| 109. | *Prognathodes brasiliensis* | Absent | Non-coral feeder | Small invertebrates, usually picked from narrow crevices or rock-surfaces with tubeworms | Kuiter (2002) |
| 110. | *Prognathodes dichrous* | Absent | Non-coral feeder | Small invertebrates, usually picked from narrow crevices or rock-surfaces with tubeworms | Kuiter (2002) |
| 111. | *Prognathodes geminus* | Absent | Non-coral feeder | Small benthic non-coral invertebrates | |
| 112. | *Prognathodes guyanensis* | Absent | Non-coral feeder | Small invertebrates, usually picked from narrow crevices or rock-surfaces with tubeworms | Kuiter (2002) |
| 113. | *Prognathodes marcellae* | Absent | Non-coral feeder | Small invertebrates, usually picked from narrow crevices or rock-surfaces with tubeworms | Kuiter (2002) |

| Sample No. | Species | IMJ | Diet Mode | Diet Composition | Reference |
|---|---|---|---|---|---|
| 114. | *Prognathodes obliquus* | Absent | Non-coral feeder | Small invertebrates, usually picked from narrow crevices or rock-surfaces with tubeworms | Kuiter (2002) |
| 115. | *Roa australis* | Absent | Non-coral feeder | Small benthic non-coral invertebrates | |
| 116. | *Roa excelsa* | Absent | Non-coral feeder | Small benthic non-coral invertebrates | Allen et al. (1998); Pratchett et al. (2013); Hodge et al. (2021) |
| 117. | *Roa haraguchiae* | Absent | Non-coral feeder | Small benthic non-coral invertebrates | |
| 118. | *Roa jayakari* | Absent | Non-coral feeder | Small benthic non-coral invertebrates | Pratchett et al. (2013); Hodge et al. (2021) |
| 119. | *Roa modesta* | Absent | Non-coral feeder | Small benthic non-coral invertebrates | Michael (2004); Pratchett et al. (2013); Hodge et al. (2021) |
| 120. | *Roa rumsfeldi* | Absent | Non-coral feeder | Small benthic non-coral invertebrates | Hodge et al. (2021) |

**Supplementary Table 3.** Average dietary composition of 36 *Chaetodontidae* species, compiled from published feeding studies based on gut-content analysis. Dietary categories reported in the original sources were consolidated into three groups: Scleractinians (hard corals), Alcyonarians (soft corals), and Other (combining non-coral benthic invertebrates, algae, and any other prey reported in the source). Category consolidation was performed within each individual source prior to averaging. For species with more than one published source, values for each category were averaged across sources (equal weighting per source). One source (*Chaetodon pelewensis*) summed to 96.9% rather than 100% as originally published and is reproduced here unadjusted.

| Species Name | Scleractinians (stony corals) (%) | Alcyonarians (soft corals) (%) | Other (%) | Sources |
|---|---|---|---|---|
| *Chaetodon auriga* | 43.57 | 2.35 | 54.08 | (Harmelin-Vivien, 1989; Harmelin-Vivien & Bouchon-Navaro, 1983; Pratchett, 2005) |
| *Chaetodon citrinellus* | 32.61 | 18.42 | 48.97 | (Harmelin-Vivien, 1989; Harmelin-Vivien & Bouchon-Navaro, 1983; Pratchett, 2005) |
| *Chaetodon kleinii* | 44.59 | 4.84 | 50.57 | (Harmelin-Vivien, 1989; Pratchett, 2005; Sano, 1989) |
| *Chaetodon lunula* | 65.10 | 1.25 | 33.65 | (Harmelin-Vivien, 1989; Harmelin-Vivien & Bouchon-Navaro, 1983; Pratchett, 2005) |
| *Chaetodon madagascariensis* | 55.00 | 15.00 | 30.00 | (Harmelin-Vivien, 1989) |
| *Chaetodon trifascialis* | 99.99 | 0.00 | 0.01 | (Bouchon-Navaro, 1986; Harmelin-Vivien, 1989; Pratchett, 2005) |
| *Chaetodon trifasciatus* | 100.00 | 0.00 | 0.00 | (Bouchon-Navaro, 1986; Harmelin-Vivien, 1989; Pratchett, 2005) |
| *Chaetodon vagabundus* | 33.81 | 4.57 | 61.62 | (Harmelin-Vivien, 1989; Harmelin-Vivien & Bouchon-Navaro, 1983; Pratchett, 2005) |
| *Chaetodon xanthocephalus* | 6.00 | 0.00 | 94.00 | (Harmelin-Vivien, 1989) |

| Species Name | Scleractinians (stony corals) (%) | Alcyonarians (soft corals) (%) | Other (%) | Sources |
|---|---|---|---|---|
| *Chaetodon ephippium* | 10.73 | 1.24 | 88.03 | (Harmelin-Vivien & Bouchon-Navaro, 1983; Pratchett, 2005) |
| *Chaetodon pelewensis* | 55.90 | 39.40 | 1.70 | (Harmelin-Vivien & Bouchon-Navaro, 1983) |
| *Chaetodon quadrimaculatus* | 27.50 | 34.10 | 38.40 | (Harmelin-Vivien & Bouchon-Navaro, 1983) |
| *Chaetodon ulietensis* | 45.05 | 23.77 | 31.17 | (Harmelin-Vivien & Bouchon-Navaro, 1983; Pratchett, 2005) |
| *Chaetodon unimaculatus* | 52.12 | 30.98 | 16.90 | (Harmelin-Vivien & Bouchon-Navaro, 1983; Pratchett, 2005; Sano, 1989) |
| *Forcipiger flavissimus* | 3.10 | 2.45 | 94.45 | (Harmelin-Vivien & Bouchon-Navaro, 1983; Sano, 1989) |
| *Heniochus chrysostomus* | 19.55 | 26.25 | 54.20 | (Harmelin-Vivien & Bouchon-Navaro, 1983; Sano, 1989) |
| *Chaetodon argentatus* | 17.00 | 0.00 | 83.00 | (Sano, 1989) |
| *Chaetodon melannotus* | 17.87 | 70.64 | 11.48 | (Pratchett, 2005; Sano, 1989) |
| *Chaetodon nippon* | 5.00 | 1.00 | 94.00 | (Sano, 1989) |
| *Chaetodon punctatofasciatus* | 94.00 | 5.00 | 1.00 | (Sano, 1989) |
| *Chaetodon rafflesii* | 20.32 | 48.98 | 30.70 | (Pratchett, 2005; Sano, 1989) |
| *Hemitaurichthys polylepis* | 0.00 | 0.00 | 100.00 | (Sano, 1989) |
| *Heniochus varius* | 43.00 | 0.00 | 57.00 | (Sano, 1989) |
| *Chaetodon baronessa* | 99.56 | 0.00 | 0.44 | (Pratchett, 2005) |

| Species Name | Scleractinians (stony corals) (%) | Alcyonarians (soft corals) (%) | Other (%) | Sources |
|---|---|---|---|---|
| *Chaetodon lunulatus* | 97.09 | 0.02 | 2.89 | (Pratchett, 2005) |
| *Chaetodon plebeius* | 95.66 | 0.00 | 4.34 | (Pratchett, 2005) |
| *Chaetodon rainfordi* | 93.07 | 1.73 | 5.20 | (Pratchett, 2005) |
| *Chaetodon speculum* | 61.66 | 0.00 | 38.34 | (Pratchett, 2005) |
| *Chaetodon lineolatus* | 0.00 | 0.00 | 100.00 | (Pratchett, 2005) |
| *Chaetodon semeion* | 0.00 | 0.00 | 100.00 | (Pratchett, 2005) |
| *Chelmon rostratus* | 0.00 | 0.00 | 100.00 | (Pratchett, 2005) |
| *Chaetodon bennetti* | 100.00 | 0.00 | 0.00 | (Bouchon-Navaro, 1986) |
| *Chaetodon ornatissimus* | 100.00 | 0.00 | 0.00 | (Bouchon-Navaro, 1986) |
| *Chaetodon reticulatus* | 100.00 | 0.00 | 0.00 | (Bouchon-Navaro, 1986) |
| *Forcipiger longirostris* | 0.00 | 0.00 | 100.00 | (Bouchon-Navaro, 1986) |
| *Chaetodon austriacus* | 100.00 | 0.00 | 0.00 | (Bouchon-Navaro, 1986) |

**Supplementary Table 4.** Crown-group diversification rates for *Chaetodontinae*, its sister lineage, *Heniochinae*, and 29 comparator reef-associated and marine fish lineages, calculated following the crown-age method-of-moments estimator of Magallón and Sanderson (2001). *N* = total number of valid described species per lineage; crown ages are given in millions of years (Ma); diversification rates (in species per lineage per million years, $Ma^{-1}$) are shown under two contrasting extinction-fraction assumptions, ε=0 (no extinction) and ε=0.9 (high extinction), to bound the range of plausible rate estimates. Species counts follow Eschmeyer's Catalog of Fishes (Fricke et al. 2026); crown ages are drawn from multiple associated phylogenomic sources, as indicated per lineage.

| Lineage | *N* | Crown age (Ma) | Diversification rate of crown when ε=0 | Diversification rate of crown when ε=0.9 | Citations for age estimate |
|---|---|---|---|---|---|
| *Chaetodontinae* | 110 | 7.78 | 0.515 | 0.312 | This paper |
| *Heniochinae* | 28 | 15.07 | 0.175 | 0.083 | This paper |
| *Scarinae* | 100 | 23.7 | 0.165 | 0.099 | Brownstein et al. 2025 |
| *Pomacanthidae* | 90 | 16.13 | 0.236 | 0.139 | Brownstein et al. 2026 |
| *Tetraodontidae* | 193 | 37.5 | 0.122 | 0.079 | Brownstein et al. 2026 |
| *Monacanthidae* | 108 | 17.8 | 0.224 | 0.135 | Brownstein et al. 2026 |
| *Pomacentridae* | 437 | 17.1 | 0.315 | 0.219 | Brownstein et al. 2026 |
| *Balistidae* | 39 | 25.0 | 0.119 | 0.061 | Brownstein et al. 2026 |
| *Gobiidae* | 1462 | 30.9 | 0.213 | 0.160 | Brownstein et al. 2026 |
| *Blenniidae* | 414 | 12.5 | 0.427 | 0.295 | Brownstein et al. 2026 |
| *Mullidae* | 109 | 47.35 | 0.084 | 0.051 | Brownstein et al. 2026 |
| *Syngnathidae* | 338 | 85.25 | 0.060 | 0.041 | Brownstein et al. 2026 |
| *Leiognathidae* | 51 | 16.86 | 0.192 | 0.103 | Brownstein et al. 2026 |
| *Julidinae* | 223 | 21.12 | 0.223 | 0.146 | Brownstein et al. 2025 |

| Lineage | *N* | Crown age (Ma) | Diversification rate of crown when ε=0 | Diversification rate of crown when ε=0.9 | Citations for age estimate |
|---|---|---|---|---|---|
| *Apogonidae* | 380 | 41.2 | 0.127 | 0.088 | Ghezelayagh et al. 2022 |
| *Labridae* | 679 | 56.1 | 0.104 | 0.075 | Brownstein et al. 2025 |
| *Scarini* | 75 | 17.8 | 0.204 | 0.117 | Brownstein et al. 2025 |
| *Sparisomatini* | 25 | 18.6 | 0.136 | 0.063 | Brownstein et al. 2025 |
| *Cirrhilabrinae* | 89 | 27.4 | 0.139 | 0.081 | Brownstein et al. 2025 |
| *Cirrhilabrus* | 52 | 6.7 | 0.486 | 0.262 | Brownstein et al. 2025 |
| *Anthiadidae* | 238 | 31.3 | 0.153 | 0.101 | Brownstein et al. 2026 |
| *Epinephelidae* | 250 | 27.7 | 0.174 | 0.116 | Brownstein et al. 2026 |
| *Serranidae* | 106 | 15.8 | 0.251 | 0.151 | Brownstein et al. 2026 |
| *Acanthuridae* | 85 | 51.7 | 0.073 | 0.042 | Brownstein et al. 2026 |
| *Holocentridae* | 90 | 52.9 | 0.072 | 0.042 | Brownstein et al. 2025 |
| *Cirrhitidae* | 35 | 10 | 0.286 | 0.143 | Ghezelayagh et al. 2022 |
| *Lethrinidae* | 46 | 31.9 | 0.098 | 0.052 | Chen and Borsa 2020 |
| *Lutjanidae* | 139 | 47.9 | 0.089 | 0.055 | Rincon-Sandoval et al. 2020 |
| *Haemulidae* | 138 | 48.3 | 0.088 | 0.055 | Tavera et al. 2018 |
| Siganus | 32 | 25.7 | 0.108 | 0.053 | Siqueira et al. 2019 |
| Antennariidae | 69 | 60.3 | 0.059 | 0.033 | Brownstein et al. 2024 |

**Supplementary Table 5a.** Classification of *Chaetodontinae*, *Heniochinae,* and 29 comparator reef-associated and marine fish lineages relative to the crown diversification rate of *Percomorpha* (N=19,110; crown age=113.0 Ma), following the confidence-interval approach of Magallón and Sanderson (2001). The following two columns represent the lower and upper bounds of the 95% confidence interval on the species richness expected at each lineage's own crown age under *Percomorpha's* estimated diversification rate, assuming high extinction (ε=0.9), the wider, more conservative of the two extinction-fraction scenarios. Lineages are classified as extremely species-rich if observed richness exceeds the upper bound, extremely species-poor if it falls below the lower bound, and within the expected range otherwise.

| Lineage | Lower bound 95% CI (species, ε=0.9) | Upper bound 95% CI (species, ε=0.9) | Classification |
|---|---|---|---|
| *Chaetodontinae* | 1.2 | 30.9 | Extremely species-rich |
| *Heniochinae* | 1.5 | 71.3 | Within expected range |
| *Scarinae* | 2.1 | 152.9 | Within expected range |
| *Pomacanthidae* | 1.5 | 79.0 | Extremely species-rich |
| *Tetraodontidae* | 4.0 | 433.5 | Within expected range |
| *Monacanthidae* | 1.6 | 92.3 | Extremely species-rich |
| *Pomacentridae* | 1.6 | 86.6 | Extremely species-rich |
| *Balistidae* | 2.2 | 169.8 | Within expected range |
| *Gobiidae* | 2.8 | 267.6 | Extremely species-rich |
| *Blenniidae* | 1.4 | 54.8 | Extremely species-rich |
| *Mullidae* | 7.0 | 865.2 | Within expected range |
| *Syngnathidae* | 77.6 | 11109.4 | Within expected range |
| *Leiognathidae* | 1.6 | 84.7 | Within expected range |
| *Julidinae* | 1.8 | 123.5 | Extremely species-rich |
| *Apogonidae* | 4.9 | 563.7 | Within expected range |
| *Labridae* | 11.9 | 1573.9 | Within expected range |
| *Scarini* | 1.6 | 92.3 | Within expected range |
| *Sparisomatini* | 1.7 | 99.2 | Within expected range |
| *Cirrhilabrinae* | 2.4 | 205.0 | Within expected range |
| *Cirrhilabrus* | 1.2 | 26.4 | Extremely species-rich |
| *Anthiadidae* | 2.9 | 275.7 | Within expected range |
| *Epinephelidae* | 2.4 | 209.8 | Extremely species-rich |
| *Serranidae* | 1.5 | 76.6 | Extremely species-rich |

| Lineage | Lower bound 95% Cl (species, ε=0.9) | Upper bound 95% Cl (species, ε=0.9) | Classification |
|---|---|---|---|
| *Acanthuridae* | 9.0 | 1166.5 | Within expected range |
| *Holocentridae* | 9.7 | 1266.0 | Within expected range |
| *Cirrhitidae* | 1.3 | 41.2 | Within expected range |
| *Lethrinidae* | 3.0 | 288.3 | Within expected range |
| *Lutjanidae* | 7.2 | 898.7 | Within expected range |
| *Haemulidae* | 7.4 | 923.8 | Within expected range |
| *Siganus* | 2.2 | 179.5 | Within expected range |
| *Antennariidae* | 15.4 | 2091.2 | Within expected range |
| *Percomorpha* | 486.1 | 70337.0 | Within expected range |

**Supplementary Table 5b.** Classification of *Chaetodontinae*, *Heniochinae,* and 29 comparator reef-associated and marine fish lineages relative to the crown diversification rate of *Eupercaria* (N=7,134; crown age=87.7 Ma), following the confidence-interval approach of Magallón and Sanderson (2001). The following two columns represent the lower and upper bounds of the 95% confidence interval on the species richness expected at each lineage's own crown age under *Eupercaria's* estimated diversification rate, assuming high extinction (ε=0.9), the wider, more conservative of the two extinction-fraction scenarios. Lineages are classified as extremely species-rich if observed richness exceeds the upper bound, extremely species-poor if it falls below the lower bound, and within the expected range otherwise.

| Lineage | Lower bound 95% Cl (species, ε=0.9) | Upper bound 95% Cl (species, ε=0.9) | Classification |
|---|---|---|---|
| *Chaetodontinae* | 1.2 | 35.0 | Extremely species-rich |
| *Heniochinae* | 1.6 | 84.7 | Within expected range |
| *Scarinae* | 2.3 | 191.6 | Within expected range |
| *Pomacanthidae* | 1.6 | 94.5 | Within expected range |
| *Tetraodontidae* | 5.1 | 595.4 | Within expected range |
| *Monacanthidae* | 1.8 | 111.5 | Within expected range |
| *Pomacentridae* | 1.7 | 104.1 | Extremely species-rich |
| *Balistidae* | 2.5 | 214.5 | Within expected range |
| *Gobiidae* | 3.4 | 351.3 | Extremely species-rich |
| *Blenniidae* | 1.4 | 64.1 | Extremely species-rich |
| *Mullidae* | 9.8 | 1275.2 | Within expected range |
| *Syngnathidae* | 151.9 | 21879.4 | Within expected range |
| *Leiognathidae* | 1.7 | 101.6 | Within expected range |
| *Julidinae* | 2.0 | 152.2 | Extremely species-rich |
| *Apogonidae* | 6.5 | 794.8 | Within expected range |
| *Labridae* | 18.1 | 2475.1 | Within expected range |
| *Scarini* | 1.8 | 111.5 | Within expected range |
| *Sparisomatini* | 1.8 | 120.4 | Within expected range |
| *Cirrhilabrinae* | 2.8 | 263.0 | Within expected range |
| *Cirrhilabrus* | 1.2 | 29.7 | Extremely species-rich |
| *Anthiadidae* | 3.5 | 363.0 | Within expected range |
| *Epinephelidae* | 2.9 | 269.7 | Within expected range |
| *Serranidae* | 1.6 | 91.4 | Extremely species-rich |

| Lineage | Lower bound 95% Cl (species, ε=0.9) | Upper bound 95% Cl (species, ε=0.9) | Classification |
|---|---|---|---|
| *Acanthuridae* | 13.2 | 1775.1 | Within expected range |
| *Holocentridae* | 14.4 | 1943.9 | Within expected range |
| *Cirrhitidae* | 1.3 | 47.4 | Within expected range |
| *Lethrinidae* | 3.6 | 381.1 | Within expected range |
| *Lutjanidae* | 10.2 | 1329.8 | Within expected range |
| *Haemulidae* | 10.5 | 1371.0 | Within expected range |
| *Siganus* | 2.6 | 227.8 | Within expected range |
| *Antennariidae* | 24.4 | 3394.7 | Within expected range |
| *Eupercaria* | 182.1 | 26257.3 | Within expected range |

**Supplementary Table 5c.** Classification of *Chaetodontinae, Heniochinae,* and 29 comparator reef-associated and marine fish lineages relative to the crown diversification rate of *Perciformes* (N=3,213; crown age=51.1 Ma), following the confidence-interval approach of Magallón and Sanderson (2001). The following two columns represent the lower and upper bounds of the 95% confidence interval for the species richness expected at each lineage's crown age under *Perciformes'* estimated diversification rate, assuming high extinction (ε=0.9), the wider, more conservative of the two extinction-fraction scenarios. Lineages are classified as extremely species-rich if observed richness exceeds the upper bound, extremely species-poor if it falls below the lower bound, and within the expected range otherwise.

| Lineage | Lower bound 95% Cl (species, ε=0.9) | Upper bound 95% Cl (species, ε=0.9) | Classification |
|---|---|---|---|
| *Chaetodontinae* | 1.4 | 58.6 | Extremely species-rich |
| *Heniochinae* | 2.2 | 175.4 | Within expected range |
| *Scarinae* | 4.6 | 516.9 | Within expected range |
| *Pomacanthidae* | 2.4 | 201.9 | Within expected range |
| *Tetraodontidae* | 18.6 | 2550.9 | Within expected range |
| *Monacanthidae* | 2.7 | 250.5 | Within expected range |
| *Pomacentridae* | 2.6 | 229.0 | Extremely species-rich |
| *Balistidae* | 5.2 | 603.3 | Within expected range |
| *Gobiidae* | 9.3 | 1200.2 | Extremely species-rich |
| *Blenniidae* | 1.8 | 123.1 | Extremely species-rich |
| *Mullidae* | 54.5 | 7757.6 | Within expected range |
| *Syngnathidae* | 3750.7 | 543640.1 | Extremely species-poor |
| *Leiognathidae* | 2.5 | 222.0 | Within expected range |
| *Julidinae* | 3.6 | 378.6 | Within expected range |
| *Apogonidae* | 27.7 | 3878.4 | Within expected range |
| *Labridae* | 144.0 | 20729.0 | Within expected range |
| *Scarini* | 2.7 | 250.5 | Within expected range |
| *Sparisomatini* | 2.9 | 277.2 | Within expected range |
| *Cirrhilabrinae* | 6.5 | 799.9 | Within expected range |
| *Cirrhilabrus* | 1.3 | 48.0 | Extremely species-rich |
| *Anthiadidae* | 9.7 | 1256.7 | Within expected range |
| *Epinephelidae* | 6.7 | 828.4 | Within expected range |
| *Serranidae* | 2.3 | 193.4 | Within expected range |

| Lineage | Lower bound 95% Cl (species, ε=0.9) | Upper bound 95% Cl (species, ε=0.9) | Classification |
|---|---|---|---|
| *Acanthuridae* | 88.2 | 12649.5 | Extremely species-poor |
| *Holocentridae* | 100.8 | 14474.2 | Extremely species-poor |
| *Cirrhitidae* | 1.6 | 84.7 | Within expected range |
| *Lethrinidae* | 10.3 | 1346.5 | Within expected range |
| *Lutjanidae* | 57.9 | 8252.7 | Within expected range |
| *Haemulidae* | 60.5 | 8632.5 | Within expected range |
| *Siganus* | 5.5 | 655.3 | Within expected range |
| *Antennariidae* | 230.0 | 33201.6 | Extremely species-poor |
| *Perciformes* | 82.6 | 11825.1 | Within expected range |

**Supplementary Table 5d.** Classification of *Chaetodontinae, Heniochinae,* and 29 comparator reef-associated and marine fish lineages relative to the crown diversification rate of *Acanthuriformes* (N=2,423; crown age=80.9 Ma), following the confidence-interval approach of Magallón and Sanderson (2001). The following two columns represent the lower and upper bounds of the 95% confidence interval for the species richness expected at each lineage's crown age under *Acanthuriformes'* estimated diversification rate, assuming high extinction (ε=0.9), the wider, more conservative of the two extinction-fraction scenarios. Lineages are classified as extremely species-rich if observed richness exceeds the upper bound, extremely species-poor if it falls below the lower bound, and within the expected range otherwise.

| Lineage | Lower bound 95% CI (species, ε=0.9) | Upper bound 95% CI (species, ε=0.9) | Classification |
|---|---|---|---|
| *Chaetodontinae* | 1.2 | 31.3 | Extremely species-rich |
| *Heniochinae* | 1.5 | 72.7 | Within expected range |
| *Scarinae* | 2.1 | 156.8 | Within expected range |
| *Pomacanthidae* | 1.6 | 80.6 | Extremely species-rich |
| *Tetraodontidae* | 4.1 | 448.9 | Within expected range |
| *Monacanthidae* | 1.6 | 94.3 | Extremely species-rich |
| *Pomacentridae* | 1.6 | 88.4 | Extremely species-rich |
| *Balistidae* | 2.2 | 174.2 | Within expected range |
| *Gobiidae* | 2.9 | 275.7 | Extremely species-rich |
| *Blenniidae* | 1.4 | 55.8 | Extremely species-rich |
| *Mullidae* | 7.2 | 902.8 | Within expected range |
| *Syngnathidae* | 83.5 | 11960.5 | Within expected range |
| *Leiognathidae* | 1.6 | 86.4 | Within expected range |
| *Julidinae* | 1.9 | 126.4 | Extremely species-rich |
| *Apogonidae* | 5.0 | 585.3 | Within expected range |
| *Labridae* | 12.4 | 1653.7 | Within expected range |
| *Scarini* | 1.6 | 94.3 | Within expected range |
| *Sparisomatini* | 1.7 | 101.4 | Within expected range |
| *Cirrhilabrinae* | 2.4 | 210.7 | Within expected range |
| *Cirrhilabrus* | 1.2 | 26.7 | Extremely species-rich |
| *Anthiadidae* | 3.0 | 284.2 | Within expected range |
| *Epinephelidae* | 2.5 | 215.7 | Extremely species-rich |
| *Serranidae* | 1.5 | 78.1 | Extremely species-rich |

| Lineage | Lower bound 95% Cl (species, ε=0.9) | Upper bound 95% Cl (species, ε=0.9) | Classification |
|---|---|---|---|
| *Acanthuridae* | 9.4 | 1221.3 | Within expected range |
| *Holocentridae* | 10.1 | 1326.8 | Within expected range |
| *Cirrhitidae* | 1.3 | 41.8 | Within expected range |
| *Lethrinidae* | 3.0 | 297.3 | Within expected range |
| *Lutjanidae* | 7.5 | 938.1 | Within expected range |
| *Haemulidae* | 7.6 | 964.6 | Within expected range |
| *Siganus* | 2.3 | 184.3 | Within expected range |
| *Antennariidae* | 16.2 | 2204.8 | Within expected range |
| *Acanthuriformes* | 62.5 | 8917.4 | Within expected range |

**Supplementary Table 6.** Brownian motion rate estimates ($\sigma^2$) for seven size-corrected, log-transformed linear morphological traits (maximum body depth, maximum fish width, head depth, lower jaw length, mouth width, minimum caudal peduncle depth, minimum caudal peduncle width) of *Chaetodontidae*. $\sigma^2$ was estimated for each trait under a Brownian motion model via fitContinuous() in the R package geiger on the time-calibrated *Chaetodontidae* phylogeny, using the 82 species with a complete set of measurements. Jaw traits are head depth, lower jaw length and mouth width; the remaining four are body traits.

| **Trait** | **$\sigma^2$** |
|---|---|
| Max body depth | 0.0019 |
| Max fish width | 0.0022 |
| Head depth | 0.0068 |
| Lower jaw length | 0.0390 |
| Mouth width | 0.0363 |
| Min caudal peduncle depth | 0.0018 |
| Min caudal peduncle width | 0.0092 |

**Supplementary Table 7.** Use of four coral genera by 22 species of *Chaetodon* and the tests that used these data. (A) Mean percentage of coral-feeding bites taken from *Acropora*, *Montipora*, *Pocillopora*, *Porites* and other coral genera, averaged across the locations reported for each species (Table 6.1 in ref. 5; values rounded to one decimal; blank = genus not reported for that species, which is not the same as 0%), with each species' class in this study (Supplementary Table 2) and the number of locations averaged. The last two columns show whether the species entered the Friedman test of the obligate corallivores (primary analysis) or of all 22 species (robustness check): Yes, included; No, in the analysis set but excluded from the Friedman test because a genus is not reported (it is still used in the pairwise tests where both genera are reported); –, not part of that analysis. (B) Friedman tests (species as blocks, genus as treatment, 3 degrees of freedom) with Kendall's coefficient of concordance (W) and the median share of coral-feeding bites per genus. (C) Pairwise two-sided Wilcoxon signed-rank tests paired by species, using every species with data for both genera; P (Holm) is adjusted across the six comparisons of each analysis (bold, P < 0.05 after adjustment). V = signed-rank statistic; the higher genus is the one with the larger share of bites; the median difference is the first genus minus the second, in percentage points.

#### A. Share of coral-feeding bites by genus, per species

| Species | Class in this study | Sites (n) | *Acropora* (%) | *Montipora* (%) | *Pocillopora* (%) | *Porites* (%) | Other genera (%) | Friedman test: obligate analysis | Friedman test: all-species analysis |
|---|---|---|---|---|---|---|---|---|---|
| *Chaetodon aureofasciatus* | Obligate corallivore | 1 | 34.2 | 4.5 | 2.3 | 10.3 | 48.3 | Yes | Yes |
| *Chaetodon austriacus* | Obligate corallivore | 1 | 30.9 | 8.6 | 24.9 | 26.8 | 8.8 | Yes | Yes |
| *Chaetodon baronessa* | Obligate corallivore | 2 | 61.0 | 12.2 | 8.2 | 0.2 | 18.5 | Yes | Yes |
| *Chaetodon bennetti* | Obligate corallivore | 3 | 1.0 | 4.3 | 3.0 | 2.4 | 89.3 | Yes | Yes |
| *Chaetodon larvatus* | Obligate corallivore | 1 | | 14.7 | | 15.8 | 69.5 | No | No |
| *Chaetodon lunulatus* | Obligate corallivore | 5 | 25.1 | 14.8 | 6.9 | 31.5 | 26.7 | Yes | Yes |
| *Chaetodon melapterus* | Obligate corallivore | 1 | 9.7 | 8.1 | 23.9 | 31.9 | 26.3 | Yes | Yes |
| *Chaetodon meyeri* | Obligate corallivore | 1 | 47.9 | 6.0 | 12.8 | 19.4 | 14.0 | Yes | Yes |
| *Chaetodon multicinctus* | Obligate corallivore | 1 | | 14.6 | 16.7 | 47.5 | 21.2 | No | No |
| *Chaetodon octofasciatus* | Obligate corallivore | 1 | 10.2 | 2.4 | 6.1 | 33.5 | 47.8 | Yes | Yes |
| *Chaetodon ornatissimus* | Obligate corallivore | 3 | 12.0 | 12.5 | 23.4 | 38.2 | 18.1 | Yes | Yes |

| Species | Class in this study | Sites (n) | *Acropora* (%) | *Montipora* (%) | *Pocillopora* (%) | *Porites* (%) | Other genera (%) | Friedman test: obligate analysis | Friedman test: all-species analysis |
|---|---|---|---|---|---|---|---|---|---|
| *Chaetodon pelewensis* | Obligate corallivore | 1 | 10.0 | 0.0 | 67.5 | 14.0 | 8.5 | Yes | Yes |
| *Chaetodon plebeius* | Obligate corallivore | 1 | 34.8 | 2.7 | 31.6 | 3.9 | 30.1 | Yes | Yes |
| *Chaetodon punctatofasciatus* | Obligate corallivore | 1 | 29.0 | 17.0 | 13.0 | 11.0 | 30.0 | Yes | Yes |
| *Chaetodon rainfordi* | Obligate corallivore | 1 | 29.5 | 0.8 | 17.0 | 3.3 | 49.4 | Yes | Yes |
| *Chaetodon reticulatus* | Obligate corallivore | 1 | 26.3 | 2.8 | 58.7 | 9.9 | 2.4 | Yes | Yes |
| *Chaetodon trifascialis* | Obligate corallivore | 5 | 89.1 | 0.3 | 9.9 | 0.0 | 0.6 | Yes | Yes |
| *Chaetodon trifasciatus* | Obligate corallivore | 3 | 52.0 | 10.2 | 2.3 | 21.0 | 14.5 | Yes | Yes |
| *Chaetodon zanzibarensis* | Obligate corallivore | 2 | 5.3 | 0.7 | 6.6 | 49.1 | 38.3 | Yes | Yes |
| *Chaetodon guttatissimus* | Facultative corallivore | 1 | 41.7 | 2.4 | 1.3 | 17.1 | 37.4 | – | Yes |
| *Chaetodon unimaculatus* | Facultative corallivore | 3 | 49.7 | 54.6 | 2.7 | 0.7 | 8.8 | – | Yes |
| *Chaetodon andamanensis* | Not in phylogeny | 1 | 11.7 | 2.4 | 35.4 | 0.0 | 50.5 | – | Yes |

### B. Friedman tests

| Analysis | Species in set (n) | Species in Friedman test (n) | $\chi^2$ (3 d.f.) | P | Kendall's W | Median share (%): *Acropora* | *Montipora* | *Pocillopora* | *Porites* |
|---|---|---|---|---|---|---|---|---|---|
| Obligate corallivores (primary) | 19 | 17 | 10.76 | 0.013 | 0.211 | 29.0 | 6.0 | 12.9 | 15.8 |
| All 22 species (robustness check) | 22 | 20 | 10.20 | 0.017 | 0.170 | 29.3 | 5.3 | 12.8 | 14.9 |

### C. Pairwise Wilcoxon signed-rank tests

| Analysis | Genus pair | n | V | P | P (Holm) | Higher genus | Median difference (percentage points) |
|---|---|---|---|---|---|---|---|
| **Obligate corallivores** | ***Acropora* vs *Montipora*** | **17** | **149** | **0.00011** | **0.00064** | ***Acropora*** | **22.3** |
| Obligate corallivores | *Acropora* vs *Pocillopora* | 17 | 108 | 0.145 | 0.436 | *Acropora* | 6.0 |

| Analysis | Genus pair | n | V | P | P (Holm) | Higher genus | Median difference (percentage points) |
|---|---|---|---|---|---|---|---|
| Obligate corallivores | *Acropora* vs *Porites* | 17 | 105 | 0.190 | 0.436 | *Acropora* | 16.4 |
| Obligate corallivores | *Montipora* vs *Pocillopora* | 18 | 34 | 0.026 | 0.105 | *Pocillopora* | −6.4 |
| **Obligate corallivores** | ***Montipora* vs *Porites*** | **19** | **22** | **0.002** | **0.010** | ***Porites*** | **−10.8** |
| Obligate corallivores | *Pocillopora* vs *Porites* | 18 | 75 | 0.663 | 0.663 | *Porites* | −4.3 |
| **All 22 species** | ***Acropora* vs *Montipora*** | **20** | **201** | **0.00006** | **0.00038** | ***Acropora*** | **17.2** |
| All 22 species | *Acropora* vs *Pocillopora* | 20 | 150 | 0.097 | 0.224 | *Acropora* | 9.3 |
| All 22 species | *Acropora* vs *Porites* | 20 | 156 | 0.058 | 0.224 | *Acropora* | 17.2 |
| All 22 species | *Montipora* vs *Pocillopora* | 21 | 60 | 0.056 | 0.224 | *Pocillopora* | −5.9 |
| All 22 species | *Montipora* vs *Porites* | 22 | 51 | 0.013 | 0.064 | *Porites* | −8.9 |
| All 22 species | *Pocillopora* vs *Porites* | 21 | 108 | 0.808 | 0.808 | *Porites* | −1.9 |

**Supplementary Table 8.** Results of univariate phylogenetic logistic regressions testing corallivory (obligate or facultative coral-feeding, coded as 1, vs. non-coral feeding, coded as 0) against each of nine log-transformed morphological traits independently (total weight, standard length, maximum body depth, maximum fish width, head depth, lower jaw length, mouth width, minimum caudal peduncle depth, and minimum caudal peduncle width). Each model was fit independently on its own complete-case subset of species (n = 82–83, varying by trait due to missing data). β = coefficient estimate; SE = standard error; z = Wald test statistic; P = Wald-type p-value. None of the nine traits showed a statistically significant univariate relationship with corallivory (all P > 0.53).

| **Predictor** | **n** | **β (Estimate)** | **SE** | **z** | **P** |
|---|---|---|---|---|---|
| log_*Total_weight* | 83 | -0.027 | 0.266 | -0.10 | 0.920 |
| log_Standard_length | 82 | -0.130 | 0.840 | -0.16 | 0.877 |
| log_Max_body_depth | 83 | -0.034 | 0.752 | -0.05 | 0.964 |
| log_Max_fish_width | 83 | -0.067 | 0.739 | -0.09 | 0.928 |
| log_Head_depth | 83 | -0.023 | 0.684 | -0.03 | 0.974 |
| log_Lower_jaw_length | 83 | -0.233 | 0.458 | -0.51 | 0.611 |
| log_Mouth_width | 83 | -0.0004 | 0.393 | -0.001 | 0.999 |
| log_Min_caudalpeduncle_depth | 83 | -0.203 | 0.701 | -0.29 | 0.772 |
| log_Min_caudalpeduncle_width | 83 | -0.339 | 0.545 | -0.62 | 0.534 |
